# Tropical hummingbird pollination networks are resistant to short-term experimental removal of a common flowering plant

**DOI:** 10.1101/2022.02.24.481682

**Authors:** Kara G. Leimberger, Adam S. Hadley, Sarah J.K. Frey, Matthew G. Betts

## Abstract

Theory predicts that the structure of plant-pollination networks should withstand disturbance, but experiments testing this prediction remain uncommon. In this study, we simulated the local extinction of a hummingbird-pollinated understory plant, *Heliconia tortuosa*, from tropical forest fragments using a replicated Before-After-Control-Impact design while quantifying hummingbird abundance and space use (383 hummingbird captures and 72 radio-tagged individuals), floral visitation rates (>19,000 observation hours), and pollination success (529 flowers). We expected that *H. tortuosa* removal would either result in (i) network collapse, in which hummingbirds vacate fragments and compromise the reproductive success of other flowering plants, or (ii) increased hummingbird reliance on alternative resources (rewiring), leading to sustained fragment use. In our experiment, hummingbird behavior and pollination were remarkably resistant to loss of *H. tortuosa*, a locally common plant species representing 30-40% of the available nectar resources on average. The exact mechanisms enabling short-term hummingbird persistence after resource removal remain unclear, as we did not discover evidence of rewiring. We hypothesize that physiological adaptations (e.g., torpor and insectivory) may have allowed hummingbird persistence, perhaps alongside high movement ability. With the important caution that short-term experiments may not emulate natural extinction processes, our study provides support for predictions that pollination networks may be robust to plant species loss.

## INTRODUCTION

Species loss can have cascading consequences for the broader community, including coextinctions of dependent species and declines in ecosystem function (Koh *et al*. 2004, Colwell *et al*. 2012). Linked extinctions may be especially likely for mutualistic interactions, such as those between plants and pollinators (Dunn *et al*. 2009, Aslan *et al*. 2013). For instance, plants that are not adequately pollinated produce fewer seeds, accruing extinction debt that becomes realized when the existing generation cannot be replaced by new recruits (Ashman *et al*. 2004, Vellend *et al*. 2006, Kuussaari *et al*. 2009). This repeated reproductive failure will eventually cause demographic collapse of plant populations (Anderson *et al*. 2011, Phillips *et al*. 2015), which could then hasten the decline of animals that consume pollen or nectar (Biesmeijer *et al*. 2006, Pauw and Hawkins 2011, Weiner *et al*. 2014, Scheper *et al*. 2014). Indeed, plants and pollinators are experiencing declines worldwide (Biesmeijer *et al*. 2006, Potts *et al*. 2010, Regan *et al*. 2015), which not only imperils the essential ecosystem services on which humans depend but also threatens the maintenance of global biodiversity (Allen-Wardell *et al*. 1998, Klein *et al*. 2007, Ollerton *et al*. 2011).

However, parallel declines of pollinators and plants are at odds with theoretical predictions suggesting that species loss should generally not result in coextinction of interaction partners (Memmott *et al*. 2004, Bascompte *et al*. 2006). Tolerance to subsequent extinctions (i.e., robustness) primarily arises from the expectation that interaction networks exhibit strongly nested topologies, such that specialists interact with subsets of partners linked to generalists (Bascompte *et al*. 2003). Under this interaction pattern, connections between specialists are uncommon, so the functional role of a specialist can be absorbed by its more abundant generalist partners (Bascompte *et al*. 2003, Bascompte and Jordano 2007). In addition to the redundancy afforded by a nested structure, models suggest that flexibility in pollinator foraging behavior – i.e., rewiring to use alternative resources – may buffer networks against disturbance (Kaiser-Bunbury *et al*. 2010, Ramos-Jiliberto *et al*. 2012, Valdovinos *et al*. 2013, Timóteo *et al*. 2016).

Despite these overall predictions and mechanisms of robustness, simulation studies have revealed that removal of highly connected, generalist species thwarts a network’s capacity to buffer perturbations, leading to rapid partner extinction (Memmott *et al*. 2004, Kaiser-Bunbury *et al*. 2010, Traveset *et al*. 2017). Thus, if external stressors such as habitat fragmentation threaten highly connected species (‘topological keystones’ sensu Jordán 2009), an extinction cascade could occur – particularly without rewiring by pollinators. Uncertainty about the robustness of real-world plant-pollinator networks still exists, however, because relatively few studies have quantified pollinator responses to deletions of other species (Appendix S1: Table S1).

In this study, we tested whether declines of a hummingbird-pollinated plant *(Heliconia tortuosa)* could have cascading consequences for communities of hummingbirds and flowering plants within a fragmented tropical forest landscape – or whether rewiring might forestall network collapse. In Neotropical forests, the genus *Heliconia* (Heliconiaceae) comprises understory herbs that have been suggested to function as “keystone mutualists” (Gilbert 1980). Large, brightly colored *Heliconia* inflorescences not only provide nectar to large-bodied hermit hummingbirds (Trochilidae: Phaethornithinae) throughout the year, but also support a variety of other hummingbird species during periods of resource scarcity (Stiles 1975, 1985). Because most hummingbird species visit multiple plant species (Rodríguez-Flores *et al*. 2019), loss of *H. tortuosa* and its associated hummingbird partners could jeopardize the reproduction of co-occurring plants, thus generating an extinction cascade (Gilbert 1980). Although *H. tortuosa* is locally common in the mid-elevation, premontane tropical forests of Costa Rica, Hadley *et al*. (2014) found that *H. tortuosa* in smaller forest fragments produced fewer seeds, possibly due to short-distance pollen transfer associated with pollinator movement limitation.

To assess the potential consequences of *H. tortuosa* declines, we conducted one of most highly replicated, large-scale removal experiments conducted within the context of plant-pollinator interaction networks (Appendix S1: Table S1). We measured hummingbird space use, floral visitation rates, and plant pollination success before and after temporary experimental removal of *H. tortuosa* within forest fragments. We developed two alternative hypotheses about how hummingbirds would respond to declines of *H. tortuosa* (hereafter *Heliconia*) and potentially affect the pollination of co-occuring plant species (Table 1). Under the *robust network hypothesis* (**H1**), hummingbirds do not permanently emigrate from forest fragments following *Heliconia* decline, leading to uninterrupted visitation to remaining plants. We hypothesized that hummingbirds compensate for *Heliconia* extinction by increasing their use of non-*Heliconia* floral resources (rewiring). Alternatively, hummingbirds may be highly dependent on *H. tortuosa* and inflexible in their foraging choices; therefore, local *Heliconia* declines should reduce nectar availability such that hummingbirds cannot afford the energetic costs to remain in (or travel to) focal forest fragments, leading to decreases in fragment use and pollination success (**H2**: *parallel extinction hypothesis*). Under both hypotheses, we expected hummingbirds to change their behavior due to their exceptionally rapid metabolisms, high dependency on nectar, and limited fat stores (Suarez 1992, Powers *et al*. 2003). Lastly, we expected that changes in *Heliconia* density would affect green hermits (*Phaethornis guy*) and violet sabrewings (*Campylopterus hemileucurus*) more strongly than other hummingbird species (**H3**). These two species (hereafter ‘*Heliconia* specialists’) have bills that are morphologically specialized for long, curved flowers and are *Heliconia*’s primary pollinators (Stiles 1979, Wolf and Stiles 1989, Taylor and White 2007, Betts *et al*. 2015). Both species inhabit the forest understory and behave as high-reward trapliners (sensu Feinsinger and Colwell 1978), visiting spatially dispersed, nectar-rich floral resources – such as *Heliconia* – in a routine sequence (Borgella *et al*. 2001).

**Table 1.**
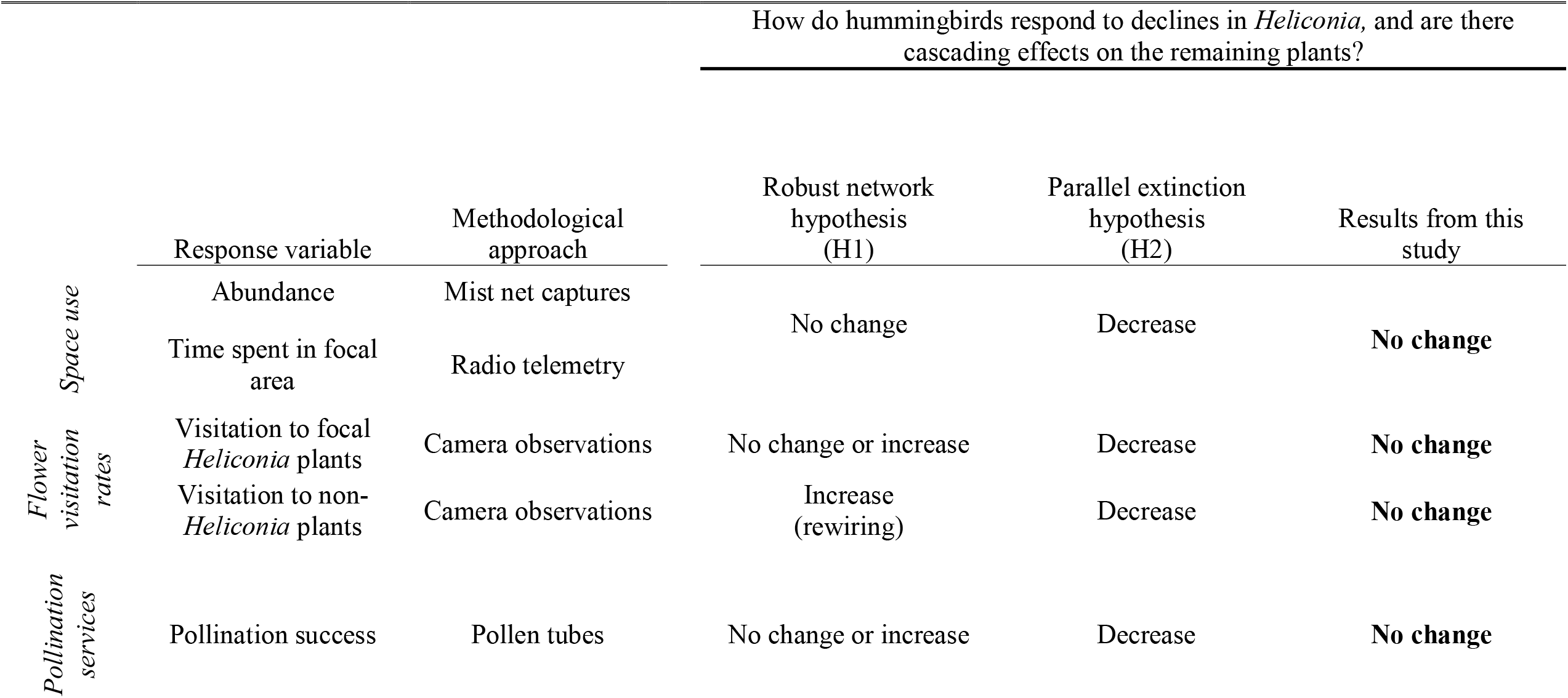
Hypotheses testing whether declines of a locally common, tropical flowering plant (*Heliconia tortuosa*) affect hummingbird space use, flower visitation rates, and plant pollination success. Predicted effects are indicated in the column below each hypothesis. We experimentally removed *H. tortuosa* using a BACI (Before-After-Control-Impact) experimental design (Fig. 2), so predicted effects can be conceptualized as pre-to-post changes in treatment replicates compared to pre-to-post changes in control replicates.

## MATERIAL AND METHODS

### Study system

We conducted this study within a **∼**122 km^2^ area surrounding the Las Cruces Biological Station in Costa Rica (Coto Brus Canton, Puntarenas Province, 8°47’7’’ N, 82°57’32’’ W, Fig. 1: A-B). This study area comprises cattle pastures, coffee plantations, and fragments of tropical premontane wet forest created during a period of extensive land clearing concentrated between 1960 and 1980 (Zahawi *et al*. 2015). Remnant forest fragments (∼30% of the study area) span a wide range of sizes (<1 to >1000 ha), with otherwise isolated fragments often connected by narrow riparian corridors and living fences (Hadley and Betts 2009, Zahawi *et al*. 2015). Mean annual temperature at the Las Cruces Biological Station is ∼21 ºC, mean annual rainfall is 3500-4000 mm, and a dry season occurs from December to March (Borgella *et al*. 2001, Zahawi *et al*. 2015).

**Figure 1.**
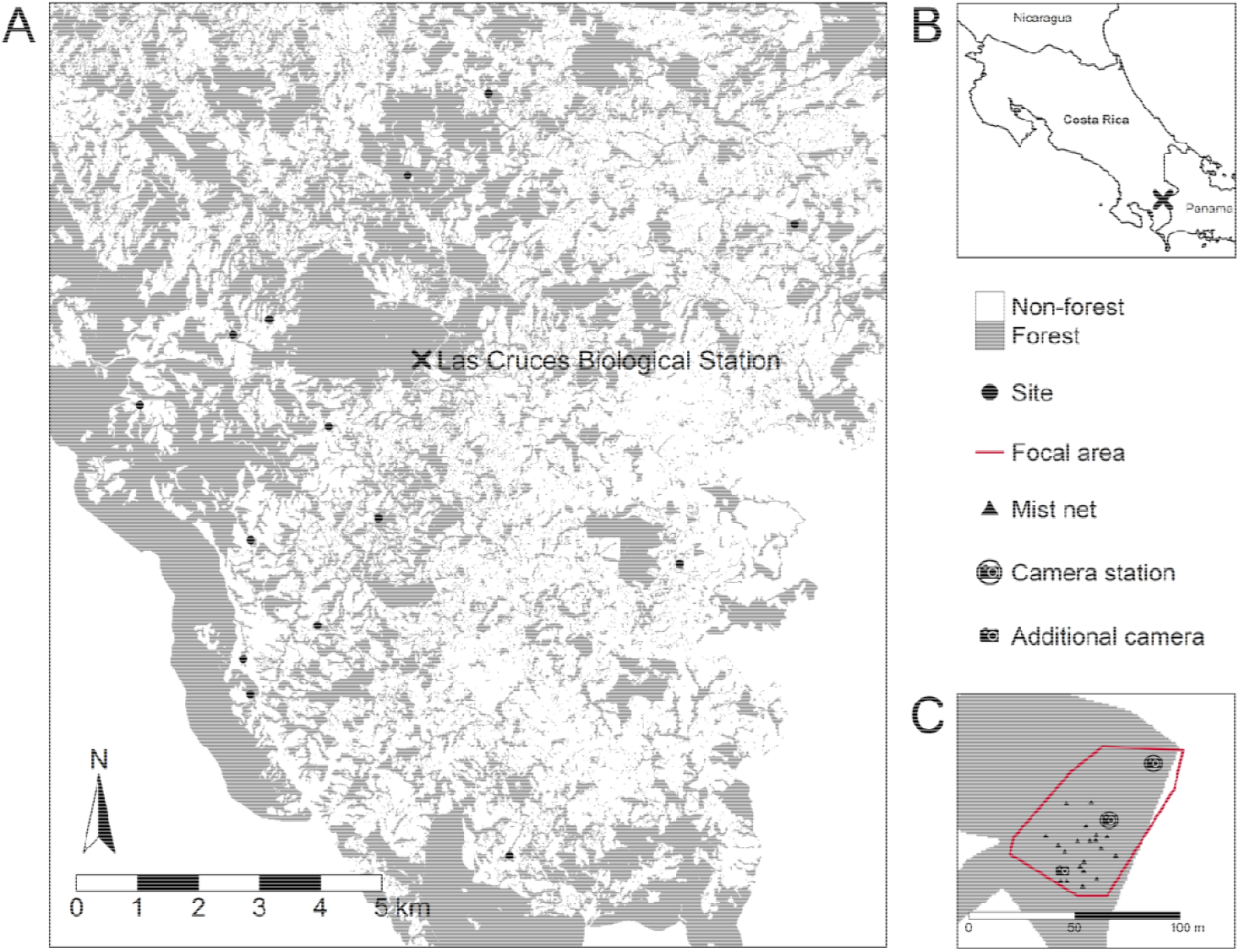
This study was conducted in the fragmented landscape surrounding the Las Cruces Biological Station (A) in southern Costa Rica (B). Within the study system (rectangular extent), we utilized fourteen sites. Forested areas are shown in grey, and non-forested areas (white) primarily comprise pasture. The forest layer is a combination of areas digitized for this study (generally within 1000 m of each site) and those digitized by Mendenhall & Wrona (2018). (C) An example site, with red line representing a focal area [i.e., where we removed *Heliconia* (for treatment replicates) and/or surveyed for floral resources (for treatment and control replicates)]. Within focal areas, we captured hummingbirds using mist nets (triangles) and monitored plant-hummingbird interactions using trail cameras (camera icons).

Based on visual examination of satellite imagery and previous work in this study area (Hadley *et al*. 2014, 2018, Kormann *et al*. 2016), we selected 14 focal fragments (‘sites’, Fig. 1A). Sites were then included based on presence of flowering *Heliconia*, landowner permission, vehicle accessibility, and navigable terrain. To ensure site accessibility, we conducted our study during the dry season and beginning of the rainy season (February – May), because localized flooding can occur during periods of heavy rainfall. Elevation of sites ranged from ∼900-1500 m (Appendix S2: Table S1).

### Experimental design

#### Site details and experimental timeline

This study used a replicated BACI (Before-After-Control-Impact) design. A BACI design requires two paired sites: an Impact site (here, ‘Treatment’) that experiences a disturbance and a Control site that does not. Additionally, each site is monitored concurrently throughout time (here, ‘Pre’ and ‘Post’ disturbance) (Fig. 2). Therefore, the comparison of interest is the pre-to-post change in the treatment site relative to the pre-to-post change in the control site. This design separates treatment effects from natural processes occurring in the study system, either across space (control *versus* treatment) or time (pre *versus* post).

**Figure 2.**
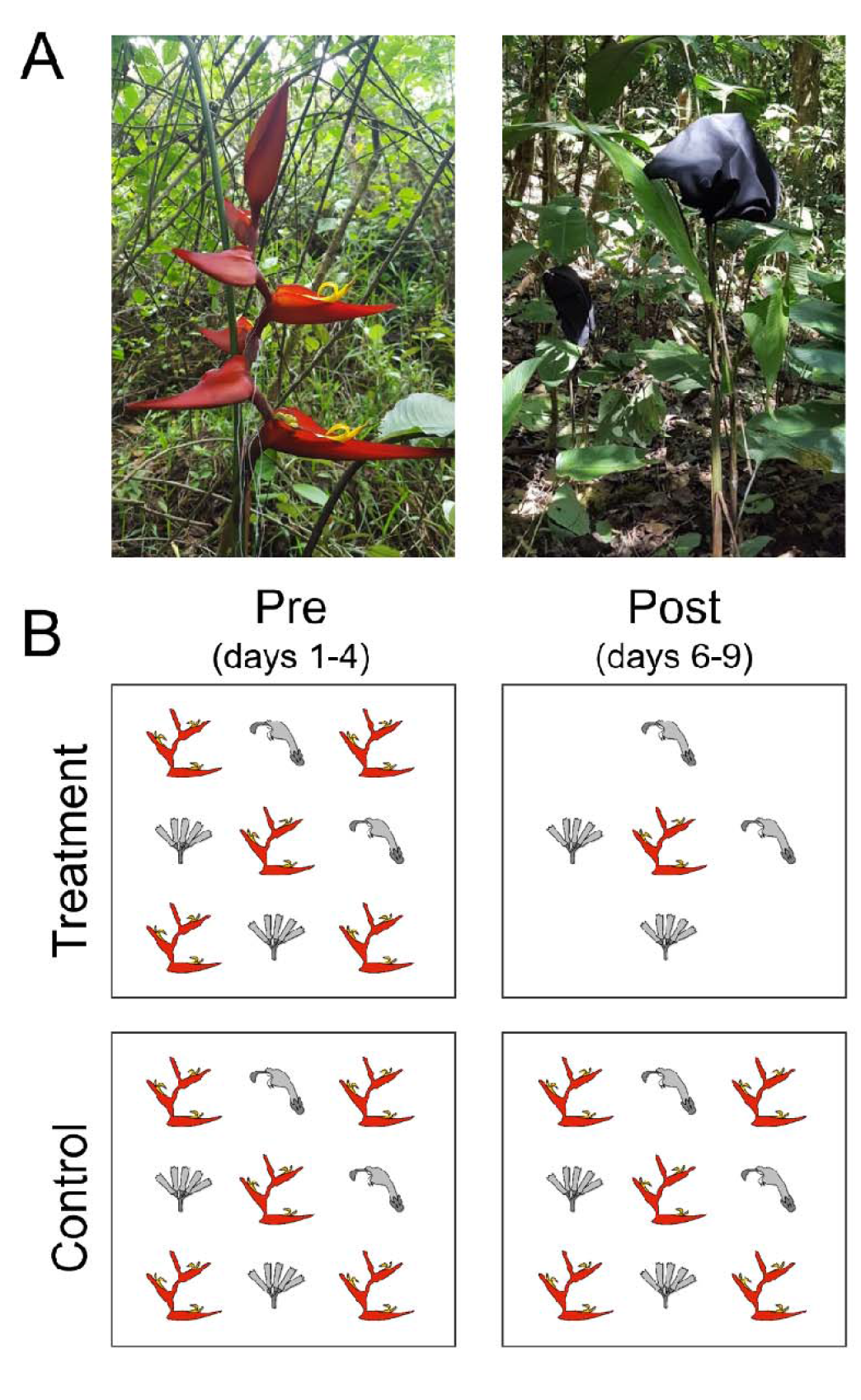
We temporarily removed *Heliconia* by covering inflorescences with fabric bags (A) as part of a Before-After-Control-Impact experimental design (B). Within a year (2016–2018), each site received either *Heliconia* removal (Treatment) or was assigned to be a Control. Treatments were reversed in alternate years for a total maximum sample size of 16 control replicates and 16 treatment replicates. Additionally, sites were sampled during two experimental periods: Before (‘Pre’) and After (‘Post’). With this experimental design, the comparison of interest is the pre-to-post change in treatment replicates compared to the pre-to-post change in control replicates. *Photos provided by M. Donald*.

Here, we paired control and treatment sites based on physical characteristics (elevation and connectivity to additional forest: Appendix S2: Table S1) and ensured that paired sites were at least 1000 m apart. This distance threshold corresponds to the estimated daily movement distance of traplining hummingbirds in this study system (Volpe *et al*. 2014), making it unlikely that hummingbirds from the treatment site would relocate to the control site. Actual distances between sites ranged from 1917-5637 m (mean ± SD: 4256 ± 1528 m, median: 5086 m), and we never detected hummingbird movement between paired sites — both in terms of hummingbird captures and radio telemetry (see *‘Response variables’*, below). Using the 14 sites, we repeated the *Heliconia* removal experiment 16 times across three years (2016-2018), yielding a maximum sample size of 16 control replicates and 16 treatment replicates (Appendix S2: Table S2). Thus, most sites were used multiple times across the three-year study period, but only once per year (Appendix S2; Table S3). Sites used multiple times received the opposite treatment of the previous year, except for 3 of 16 instances (when covering *Heliconia* in that site again would have posed substantial logistical challenges). Initial treatment assignments were random, which reduces the likelihood that any treatment differences would be due to factors associated with individual sites.

Each experimental replicate lasted for 9 days. Starting around noon on the fourth day, we removed *Heliconia* from the treatment site and designated the following 36 hours as a behavioral adjustment period during which no data were collected. Although this adjustment period was necessarily relatively short due to the limited battery lifespan of hummingbird radio-transmitters (see *‘Radio telemetry’*, below), we expected hummingbirds to respond rapidly due to their exceptionally high metabolisms and limited fat stores (Suarez 1992, Powers *et al*. 2003).

#### Heliconia removal (hummingbird exclusion)

Across all study sites and years, we removed between 8 and 526 *Heliconia* inflorescences per treatment replicate (mean ± SD: 172 ± 138, median: 138, Appendix S2: Table 4) by covering *Heliconia* inflorescences (mature and immature) with dark-colored fabric bags (Fig. 2A). This approach prevented hummingbird access and obscured flowers from view while avoiding long-term changes to floral resources associated with destructive sampling methods (e.g., cutting). The covered inflorescences represented, on average, 97% (± 5% SD) of the *Heliconia* inflorescences encountered during removal (range: 80-99%); two focal *Heliconia* plants/site remained uncovered for pre-to-post monitoring, and sometimes plant inaccessibility limited our covering capacity. While covering *Heliconia*, we also collected data on the abundance and location of resources available to hummingbirds (*Heliconia* and non-*Heliconia*). Resource data were used to estimate the percentage of calories removed by our manipulation (see Appendix S3 for methods) and delineate ‘focal areas’ (see below). In 2017-2018, we also collected resource data in the control sites, mimicked the trampling effect experienced by the treatment sites.

#### Focal areas

Within each site, we focused our observations within designated ‘focal areas’. Focal areas encompassed locations where we captured hummingbirds, installed trail cameras, and set up plant arrays (Fig. 1C, see *‘Response variables’*). We delineated focal areas as the 100% minimum convex polygons of the plant GPS points. Treatment focal areas ranged from 0.27 to 7.49 ha (mean ± SD: 1.7 ± 1.9, median: 1.25, Appendix S2: Table S4), depending on the size of the forest fragment and the navigability of the terrain. We acknowledge that our focal areas were smaller, on average, than hummingbird home ranges in this study system (e.g., mean home range size of 2.2 ha for green hermits: Volpe *et al*. 2016). However, here we were interested in any behavioral changes occurring *within* the focal area, following similar removal studies (e.g., 20 *x* 20-m plots within the home range of a bumblebee: Brosi & Briggs 2013).

### Response variables

#### Hummingbird captures

Within each focal area, we captured hummingbirds at the beginning and end of the experimental period (Days 1 and 9). During each capture session, positioned 5-20 mist nets (12-m and 6-m in length) near flowering plants visited by hummingbirds (e.g., the genera *Heliconia, Centropogon, Palicourea, Musa*). Nets were opened 30 minutes after sunrise and remained open for 2-5 hours per session. Capture effort (net-hours) and net locations were kept constant for both capture sessions, except in one instance (Appendix S2: Fig. S1). Following the North American Banding Council’s hummingbird protocol (Russell and Russell 2001), we tagged hummingbirds with an aluminum tarsus band, measured body mass to the nearest 0.01 g using an electronic balance, and measured wing length to the nearest 0.5 mm using a wing rule. Tarsus bands allowed detection of recaptured hummingbirds within and between capture sessions; individuals recaptured during the same capture session were only counted once. In 2017-2018, we also marked hummingbirds’ heads with unique color combinations of nail polish following Hadley *et al*. (2014); this additional mark allowed us to identify individual hummingbirds seen on camera (see *‘Flower visitation’*). All birds were handled in accordance with Oregon State University Animal Care and Use Protocols (ACUP #4655, #5015).

#### Radio telemetry

We attached miniaturized radio transmitters (0.15-0.25 g, Blackburn Transmitters, Nacogdoches, TX) to 72 hummingbirds according to the methods of Hadley & Betts (2009). We then followed hummingbirds on foot using portable receivers (Wildlife Materials Inc TRX 1000S) and handheld Yagi antennae, recorded locations continuously when the bird was found within receiver range (∼250 m in the absence of topographical barriers). In total, we completed 450 hours of telemetry observation across Days 2-4 and Days 6-8 of the experiment, averaging 3.1 ± 0.9 hours per session (typically ∼8:00-11:00) and 5.4 ± 0.9 sessions per site (mean ± SD). From the recorded locations, we estimated the proportion of observation time that radio-tagged birds spent in the focal area. Thirty-six hummingbirds spent time in the focal area during the ‘pre’ period and were included in analyses of the *Heliconia* removal experiment; for full methods and a comprehensive summary of radio-tracking outcomes, see Appendix S2.

#### Flower visitation

To understand hummingbird resource use within each site and to monitor for the presence of marked individuals, we positioned trail cameras (PlotWatcher Pro, Day 6 Outdoors) near inflorescences of plant species known or suspected to be visited by hummingbirds (*N* = 34, Appendix S2: Table S6). Effort within each site was concentrated at two locations (‘stations’, Fig. 1C) located on roughly opposite ends of the focal area (mean ± SD distance between stations: 60 ± 44 m). Stations were centered around a focal *Heliconia* plant expected to flower throughout the experimental period, not in an extremely steep area, and >10 m from the fragment edge. The two focal *Heliconia* plants remained uncovered for the entire experimental period, which allowed us to observe whether visitation rates changed after declines in surrounding *Heliconia* density. To partially standardize plant species composition between control and treatment replicates, we added a floral array comprising ∼3-5 potted plant species to each station (see Appendix S2 and *‘Pollination success’*).

Trail cameras took one photograph per second during daylight hours (5:30-17:30 on Days 2-4 and Days 6-8) and automatically combined these photographs into a time-lapse video. We reviewed videos for hummingbird sightings using the motion detection program MotionMeerkat (Weinstein 2015) or by manually watching the sped-up video. When hummingbirds were detected on camera, we identified the species, recorded any color marks, and noted whether they visited any flowers. From video, we also recorded the number of open flowers available per day. Dates with no available flowers were excluded from analysis, yielding 20,735 total hours of video. We used subsets of these data for a variety of purposes, including estimating the total floral resources available for hummingbirds under ‘normal’ (non-experimental) circumstances. The analysis for the experiment used 19,870 video hours (Appendix S2: Table S7).

#### Pollination success

To investigate whether *Heliconia* declines influenced pollination success, we examined styles for the presence of pollen tubes, which indicate pollen grain germination and subsequent movement toward the flower ovule(s) (Kearns and Inouye 1993). We focused our pollen tube investigation on the two focal *Heliconia* plants per replicate, as well as several species from the focal arrays (Appendix S2: Table S6). Throughout the experimental period, we collected flowers from these focal plants every 1-2 days, which corresponds to the maximum floral longevity for most of the selected species (*Heliconia tortuosa, Hamelia patens*: Stratton 1989, *Stachytarpheta frantzii*: Thomas-Granger 2003). Styles were fixed in formalin-acetic acid-alcohol for at least 24 hours, stained with aniline blue, and examined for pollen tubes using epifluorescence microscopy (Kearns and Inouye 1993, Betts *et al*. 2015). Due to low pollen tube presence and/or challenges with pollen tube visualization, we were only able to analyze two plant species: *H. tortuosa* (*N* = 327 styles) and *H. patens* (*N* = 202 styles). Additional details about style collection and lab procedures are available in Appendix S2.

#### Additional response variables

To supplement the primary response variables described above, we examined several additional metrics: (1) recapture probability (i.e., proportion of hummingbirds recaptured in the ‘post’ period), (2) floral visitation rates from individual hummingbirds marked with nail polish, and (3) body mass of hummingbirds captured during both experimental periods (pre and post). The first two response variables assessed the possibility that hummingbirds caught during the initial capture sessions were rapidly replaced by new individuals, potentially obscuring treatment effects. Body mass measurements allowed us to explore whether birds that remained in (or returned to) treatment focal areas paid an energetic cost. To compare mass changes across hummingbirds of different structural sizes, we calculated a relative body mass measure: observed mass divided by the mass predicted by species-specific allometric equations (i.e., Ln(body mass) ∼ Ln(wing length), Appendix S2: Fig. S4).

### Statistical analysis

We used linear mixed models (LMMs) and generalized linear mixed models (GLMMs) to analyze how experimental *Heliconia* extinction influenced (i) hummingbird abundance, (ii) proportion of time that radio-tagged hummingbirds spent in focal area, (iii) visitation rates to flowering plants (*Heliconia* and non-*Heliconia*), (iv) plant pollination success, and (v) the presence and condition of individual hummingbirds (see ‘*Additional response variables’*). For visitation to non-*Heliconia*, our main analysis examined overall visitation pattern across all plant species; however, we also analyzed individual species when sufficient data were available (nine species had >10 replicates each). We modeled each response variable using appropriate statistical distributions and controlled for variable effort when applicable (Appendix S2: Tables S8-S10). All statistical analyses were conducted in R version 4.2.0 (R Core Team 2022) using ‘glmmTMB’ (Brooks *et al*. 2017). We checked statistical assumptions by visually inspecting plots from the ‘performance’ package (Lüdecke *et al*. 2021) for LMMs and ‘DHARMa’ (Hartig 2020) for GLMMs.

In a BACI experimental design, an overall treatment effect is indicated by a statistically significant interaction between the variables representing treatment and experimental period (Underwood 1994, McDonald *et al*. 2000). Therefore, we tested the *parallel extinction hypothesis* (**H1**) and *robust network hypothesis* (**H2**) using a two-way interaction between *Heliconia* removal and period (pre *versus* post) for all response variables measured during both experimental periods (recapture probability was measured in the post period only). All models included a random intercept for ‘site’. Except when modeling recapture probability, we also included a nested random effect for ‘replicate’ (site-year), which paired pre-to-post observations within each replicate. Additional nested random effects for plant species and individual birds were included where applicable (Appendix S2: Tables S8-S9). Because we were interested in pre-to-post changes in our response variables, we only included covariates that varied between periods. Specifically, when analyzing flower visitation, we accounted for the attractiveness of a given focal plant by including a covariate for the number of open flowers on camera per day, averaged across the experimental period.

For most response variables, we implemented more than one statistical model. Since the magnitude of *Heliconia* removal varied widely (Appendix S2: Table S3), we conducted analyses with a categorical treatment variable (control/treatment) and a quantitative treatment variable, Ln(calories removed/hectare + 1). Results were qualitatively similar, so only results from the categorical approach are presented in the main text. Additionally, to determine whether declines in *Heliconia* affected *Heliconia* specialists more strongly than other hummingbird species (**H3**), we analyzed hummingbird-centric response variables using (i) the full dataset containing all species, and (ii) a dataset including green hermits and violet sabrewings. The one exception was when analyzing flower visitation to individual non-*Heliconia* plant species; here, we focused on the full dataset because *Heliconia* specialists visited some species at very low rates (or not at all) and only used the categorical treatment variable (to limit the total number of models run).

## RESULTS

Across all sites and years, we detected 19 hummingbird species in the forest fragments surrounding the Las Cruces Biological Station (Appendix S4: Table S1). We estimate that, on average, the covered *Heliconia* plants supplied 30% (± 25%, range: 0.4 – 87%) of the calories available for this entire hummingbird community and 37% (± 28%, range: 0.4 – 94%) of the calories available for green hermits and violet sabrewings (Appendix S4: Table S2). Although green hermits and violet sabrewings visited >20 plant species, highest visitation rates were observed for *Pitcairnia imbricata, Erythrina costaricensis, Musa x paradisiaca, Heliconia tortuosa*, and *Costus barbatus* (Appendix S4: Fig. S1). *Heliconia* inflorescences were visited by hummingbirds 6.4 ± 5.7 times per day (range: 0-25, median: 5), mostly by green hermits (75% of 1,055 visits, Appendix S4: Table S3).

### Mist net captures & radio telemetry

We found little evidence that *Heliconia* removal reduced captured hummingbird abundance or decreased the time that radio-tagged hummingbirds spent in the focal area. In treatment replicates, we observed a pre-to-post decrease in hummingbird captures that was, on average, 19% greater relative to control replicates but confidence intervals suggest a wide range of possible effects (95% CI: -47% to +20%, Fig. 3A, Appendix S4: Table S4). Radio-tagged hummingbirds from treatment replicates showed a similar pre-to-post decrease in the percentage of time spent in the focal area (−18% relative to control replicates, 95% CI: -53% to +40%, Fig. 3B, Appendix S4: Table S4). Results were consistent when examining all species together and *Heliconia* specialists separately (Appendix S4: Fig. S2, Tables S5-S6).

**Figure 3.**
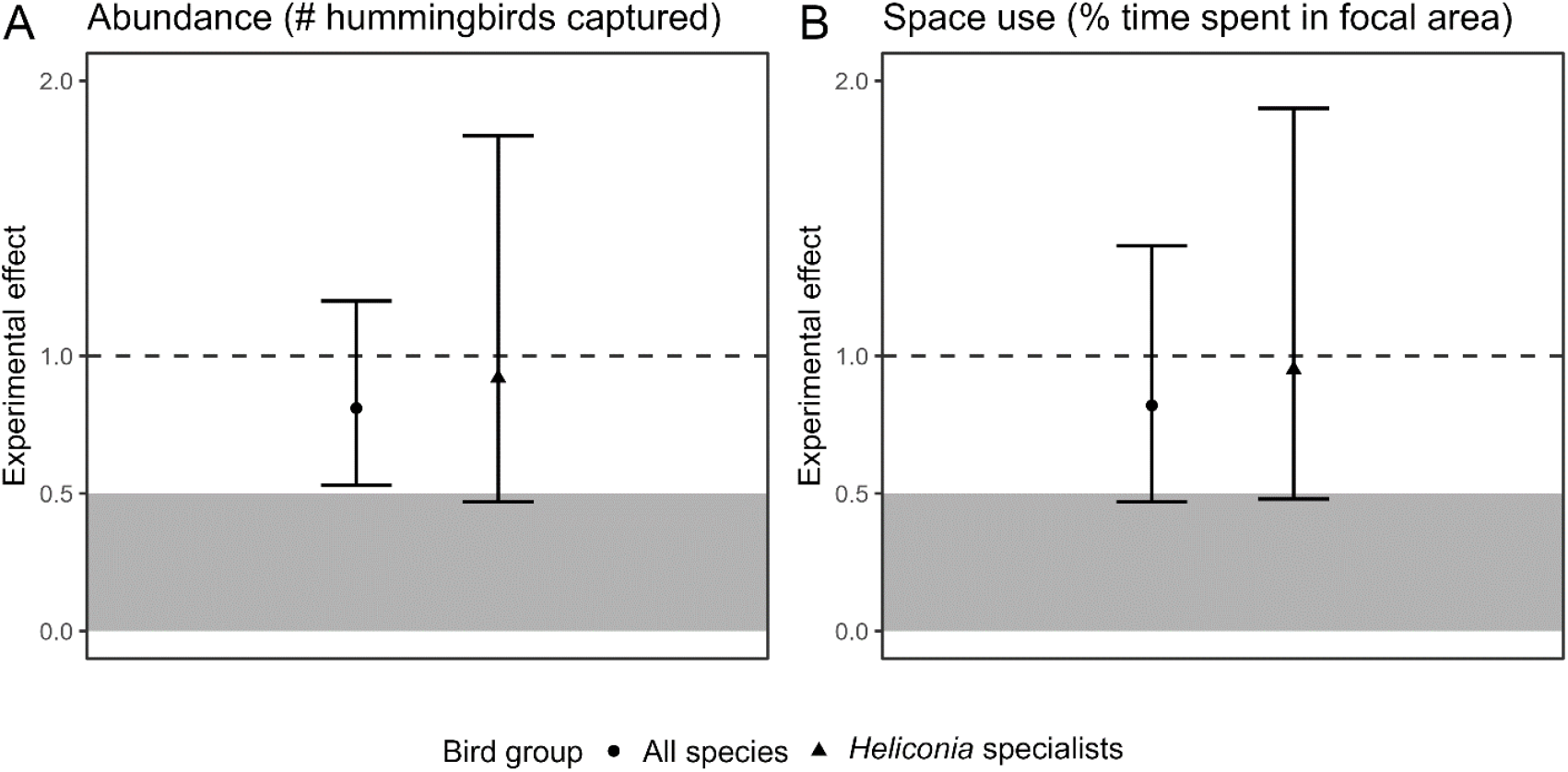
Effects of experimental *Heliconia* removal on hummingbird abundance (A) and space use (B). Hummingbird abundance was quantified as the number of hummingbirds captured in mist nets, controlling for capture effort (net-hours). Space use was quantified as the percentage of time that radio-tagged hummingbirds spent in the focal area. Following the Before-After-Control-Impact experimental design, the *y*-axis reflects the pre-to-post change in treatment replicates relative to pre-to-post change in control replicates. Estimates and 95% confidence intervals are contrasts calculated in ‘emmeans’ (Lenth 2020). The dashed line at 1 indicates no experimental effect; the grey area indicates strong support for the *network collapse hypothesis* (>50% decrease in treatment replicates relative to control replicates). *Heliconia* specialists refers to the two hummingbird species expected to rely most strongly on *H. tortuosa*: green hermits (*P. guy*) and violet sabrewings (*C. hemileucurus*).

### Flower visitation & pollination success

We continuously monitored hummingbird visits to the two focal *Heliconia* inflorescences that remained uncovered throughout the experimental period, as well as numerous other flowering plant species (mean ± SD: 7.8 ± 2 species/replicate; range: 2-11). Under the *robust network hypothesis*, we expected hummingbirds to rewire and increasingly use non-*Heliconia* resources. However, pre-to-post changes in hummingbird visitation rate were non-distinguishable between control and treatment replicates, both when examining the focal *Heliconia* plants (95% CI: -19% to +80%) and non-*Heliconia* plant species analyzed together (−29% to +10%) (Fig. 4A). Responses by *Heliconia* specialists mirrored the responses observed for all species (Fig. 4A, Appendix S4: Tables S7-S8), although these species visited non-*Heliconia* species relatively infrequently (<1 visit/day, on average: Appendix S4: Fig. S3). When analyzing non-*Heliconia* plant species individually, treatment replicates showed greater pre-to-post declines (on average) for 7 of 9 species, but confidence intervals encompassed a range of possible effects (Fig. 4B, Appendix S3: Table S9). We also found no evidence that experimental *Heliconia* removal influenced pollination success for the focal *Heliconia* plants (*z* = 0.15, *P* = 0.88) or another plant species from the floral arrays (*Hamelia patens: z* = 1.03, *P* = 0.30) (Fig. 4C, Appendix S3: Fig. S4, Table S10).

**Figure 4.**
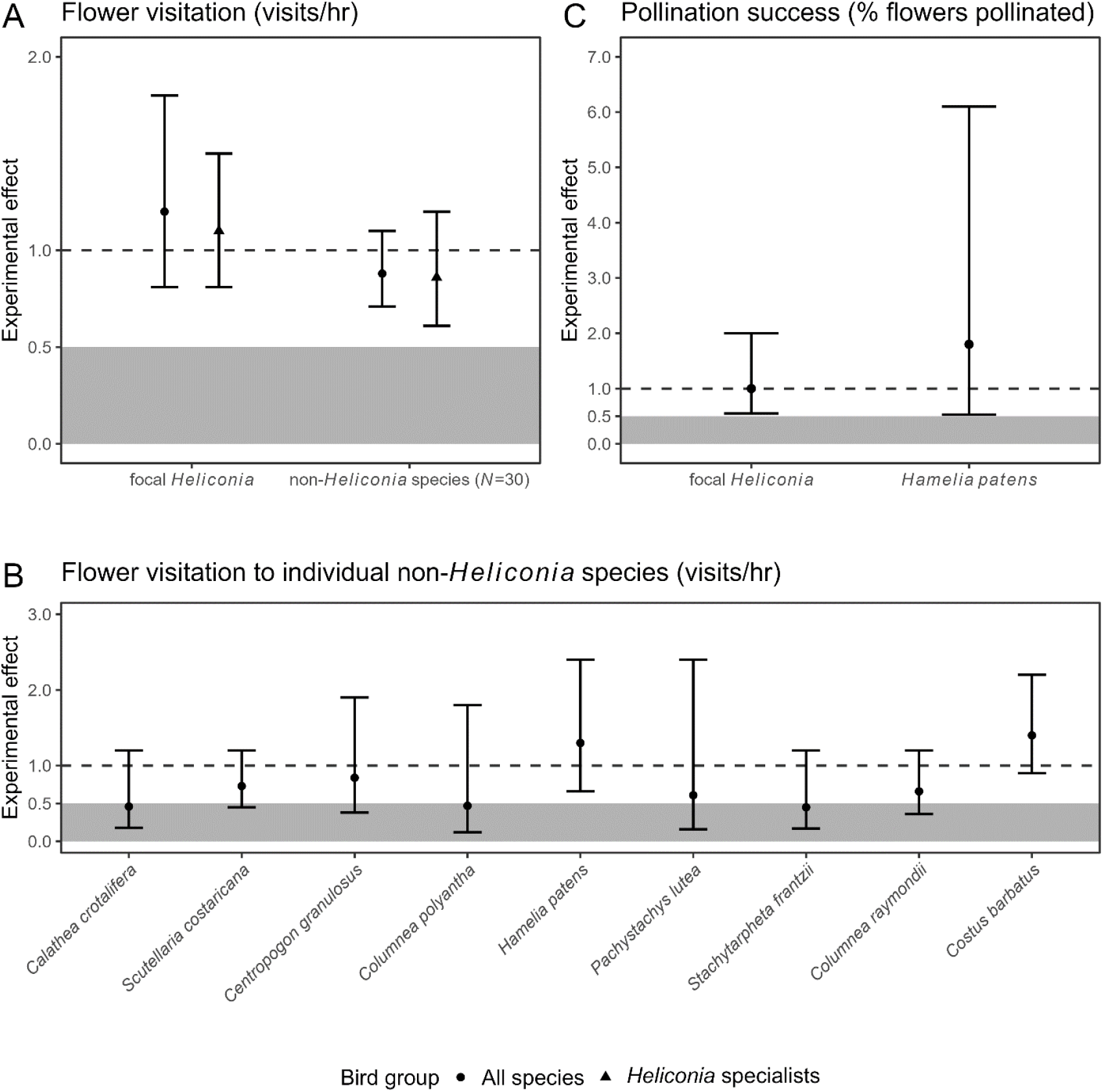
Effects of experimental *Heliconia* removal on flower visitation (A-B) and pollination success (C). Flower visitation was quantified using trail cameras that recorded hummingbird visits to focal *Heliconia* (2 inflorescences/site) and a 30 non-*Heliconia* plant species (Appendix S2: Table S6). Pollination success was quantified as the percentage of flowers with pollen tubes. We analyzed non-*Heliconia* visitation across all plant species (A) and, when sufficient data were available, to individual plant species (B). Following the Before-After-Control-Impact experimental design, the *y*-axis reflects the pre-to-post change in treatment replicates relative to pre-to-post change in control replicates. Estimates and 95% confidence intervals are contrasts calculated in ‘emmeans’ (Lenth 2020). The dashed line at 1 indicates no experimental effect; the grey area indicates strong support for the *network collapse hypothesis* (>50% decrease in treatment replicates relative to control replicates). *Heliconia* specialists refers to the two hummingbird species expected to rely most strongly on *H. tortuosa*: green hermits (*P. guy*) and violet sabrewings (*C. hemileucurus*).

### Additional response variables

When focusing further on the responses of marked individual hummingbirds, we also did not observe an effect of *Heliconia* removal. On average, the estimated probability of recapturing a hummingbird from the ‘pre’ capture session was 18% in control replicates (95% CI: 11% to 29%), compared to 14% in treatment replicates (95% CI: 8% to 23%). This difference was not statistically significant, and recaptures of *Heliconia* specialists showed qualitatively similar results (Fig. 5A, Appendix S4: Table S11). Results were consistent when examining pre-to-post changes in flower visitation for individual color-marked hummingbirds (Fig. 5B, Appendix S4: Fig. S5, Tables S12-13). Finally, we found no evidence that pre-to-post changes in hummingbird body condition differed between control and treatment replicates (Fig. 5C), either for all hummingbird species (z = -0.49, *P* = 0.62) or *Heliconia* specialists (*z* = -0.22, *P* = 0.83) (Appendix S4: Table S14).

**Figure 5.**
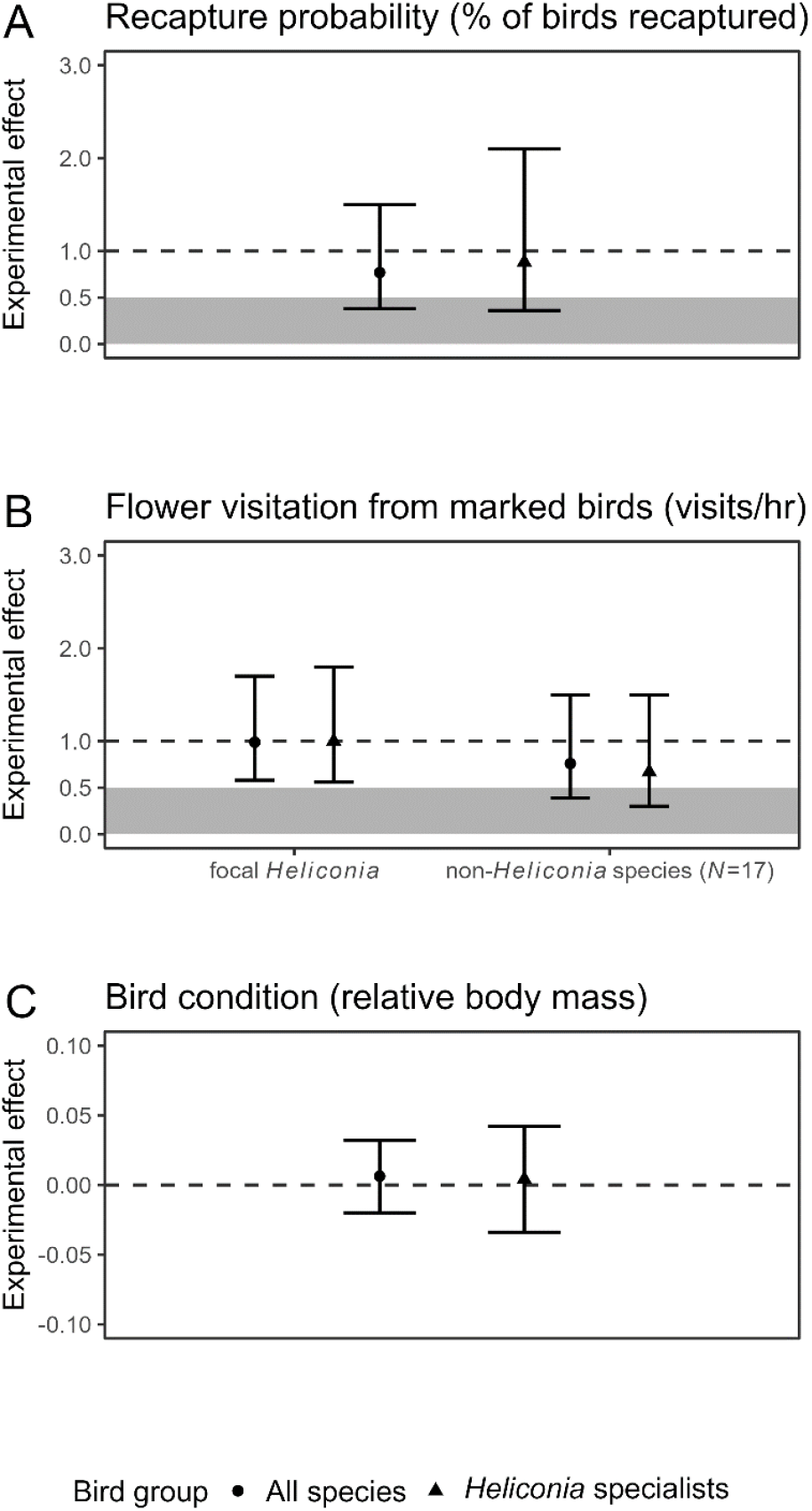
Effects of experimental *Heliconia* removal on additional response variables, which investigate whether hummingbirds present at the start of the experiment were replaced by new individuals (A-B, mist net recaptures and flower visitation of marked birds) or suffered an energetic cost by remaining in the focal area (C, bird condition). Following the Before-After-Control-Impact experimental design, the *y*-axis reflects treatment replicates relative to control replicates (A) or the pre-to-post change in treatment replicates relative to pre-to-post change in control replicates (B, C). Estimates and 95% confidence intervals are contrasts calculated with ‘emmeans’ (Lenth 2020). The dashed lines at 1 and 0 indicate no experimental effect; the grey area indicates strong support for the *network collapse hypothesis* (>50% decrease in treatment replicates relative to control replicates). *Heliconia* specialists refers to the two hummingbird species expected to rely most strongly on *H. tortuosa*: green hermits (*P. guy*) and violet sabrewings (*C. hemileucurus*).

## DISCUSSION

In this study, we experimentally induced the local extinction of a locally abundant tropical plant, *Heliconia tortuosa*, to assess the robustness of plant-hummingbird interaction networks in a fragmented tropical landscape. Based on suite of complementary response variables we found that, even when high proportions of total available calories were experimentally removed (∼30-40% on average), hummingbird responses fell within the natural variability of control replicates. As might be expected with such small changes in hummingbird behavior, we did not observe any cascading effects of experimental manipulation on pollination success of flowering plants, which supports the *robust network hypothesis* (**Hl**). However, the mechanism underlying the robust network hypothesis — hummingbirds expanding their diet to include to alternative floral resources — did not receive strong support, suggesting other explanations for hummingbird persistence.

Our finding that that experimental removal of *Heliconia* did not elicit parallel extinctions in pollinators aligns with theoretical predictions (Memmott *et al*. 2004, Kaiser-Bunbury *et al*. 2010) and several other experiments that removed plant species from mutualistic networks (Ferrero *et al*. 2013, Goldstein and Zych 2016, Costa *et al*. 2018). If hummingbirds remained in forest fragments despite *Heliconia* declines, we assumed that they would rewire to use alternative floral resources; hummingbirds as a taxonomic group are highly opportunistic foragers (Arizmendi and Ornelas 1990, Maruyama *et al*. 2013) and the most frequent *Heliconia* visitor, the green hermit, has a relatively wide niche breadth (Borgella *et al*. 2001, Betts *et al*. 2015). Unexpectedly, we found no evidence that hummingbirds increasingly visited non-*Heliconia* resources when *Heliconia* became less available, suggesting that the mechanisms underlying network robustness in this study may not necessarily align with the rewiring observed in previous empirical tests (e.g., Timóteo *et al*. 2016, Kaiser-Bunbury *et al*. 2017, Costa *et al*. 2018). This apparent lack of foraging flexibility may emerge from various constraints, such as morphological trait-matching, search costs, and/or competition from other hummingbirds (see also Weinstein and Graham 2017). For example, high levels of exploitative competition for non-defendable resources may have discouraged hummingbirds from modifying their trapline and risking overlap with another bird’s foraging route (Young 1971, Gill 1988, Ohashi and Thomson 2009). Lastly, is possible that fine-scale rewiring did occur but was not detected using our analytical approach, which aggregated visitation rates to the level of hummingbird species; further analysis examining the diet of individual hummingbirds may shed light on the possibility of rewiring at finer scales.

In the absence of rewiring, the resilience in hummingbird behavior, abundance, and pollination services following experimental *Heliconia* removal is somewhat surprising. Brief pauses in nectar intake can disrupt a hummingbird’s daily energy balance, given their high metabolic rate and limited energy reserves (Calder and Booser 1973, Tooze and Gass 1985, Brice 1992). Moreover, previous nectar manipulations with hummingbirds – albeit within defended territories – led to rapid changes in territory size, foraging patterns, and/or defense behavior, generally within a day (e.g., Hixon *et al*. 1983, Gass and Sutherland 1985, Eberhard and Ewald 1994, Temeles *et al*. 2004). One explanation for our result is that hummingbirds permanently emigrated from fragments after *Heliconia* removal but, because of high intruder pressure, were quickly replaced by new individuals (Stiles and Wolf 1970). However, we monitored individual birds using several different approaches (radio telemetry, recaptures, flower visitation) and found no evidence that *Heliconia* removal caused emigration and subsequent replacement. An alternative possibility is that hummingbirds relied on physiological adaptations, such as a torpor and/or insectivory, to maintain body condition despite large declines in nectar availability – resulting in the stable body mass measurements we observed in recaptured hummingbirds. Torpor is an energy-saving adaptation characterized by declines in body temperature and metabolic rate (Hainsworth *et al*. 1977, Spence and Tingley 2021), and even tropical hummingbirds inhabiting low-to-mid elevations appear to possess this capability, including the green hermit (Krüger *et al*. 1982, Schuchmann and Prinzinger 1988, Bech *et al*. 1997, Shankar *et al*. 2020). Additionally, hummingbirds are highly insectivorous and, although they cannot completely replace nectar with insects (Brice 1992, Stiles 1995), they may supplement their diet with arthropods when nectar becomes scarce (Young 1971, Kuban and Neill 1980, Montgomerie and Redsell 1980, Hazlehurst and Karubian 2018). Finding torpid hummingbirds at night is extremely challenging (Carpenter and Hixon 1988), but future research could explore changes in arthropod consumption using techniques such as DNA metabarcoding (Moran *et al*. 2019).

Another explanation for hummingbird persistence following experimental *Heliconia* removal involves their high movement ability. *H. tortuosa* is a patchily distributed species that necessitates long-distance foraging movements, as a single clump does not make territoriality energetically profitable (Stiles 1975). Densities can also vary substantially across short distances; for instance, we observed a 150-fold difference in *Heliconia* availability across sites and years in the present study (Appendix S4: Table S15). Similarly, Stiles (1979) notes that some Costa Rican *Heliconia* species can be common in one watershed, yet nearly absent in an adjacent watershed < 1 km away (see also Bruna and Kress 2002). In accordance with this natural history, the most frequent *Heliconia* visitor, the green hermit, can travel long distances (>500 m) while foraging within home ranges of >150 ha (although home range sizes of ∼2 ha are more typical: Volpe *et al*. 2014, 2016). Thus, returning to our focal areas to visit the few remaining *Heliconia* resources could be perceived as a relatively low-cost option for long-distance trapliners, at least in the short term. Notably, although highly mobile pollinators might buffer against network collapse (Volpe *et al*. 2016), they may be vulnerable to habitat loss and fragmentation once thresholds are crossed, leading to parallel pollinator declines observed in nature (Andrén 1994, Hadley and Betts 2012). This strategy is also unlikely to be profitable over longer time scales than measured by our experiment, which only removed *Heliconia* for five days.

Although a short adjustment period (<1 day) is standard in similar species removal experiments (e.g., Hixon *et al*. 1983, Brosi and Briggs 2013, Hazlehurst and Karubian 2018), extending the experimental timeline may have more realistically simulated a future extinction and provided more time for behavioral or demographic adjustment. If hummingbirds were not permanent residents of our focal areas – as suggested by our relatively low recapture rates (∼15%) and proportion of time spent in the focal area (∼20%) – then transient individuals might have not immediately noticed changes in *Heliconia* density and changed their foraging patterns accordingly. Likewise, hummingbirds might not have persisted had we removed *Heliconia* over larger areas, because traveling to isolated plants would become even more energetically costly (Heinrich and Raven 1972). Unfortunately, covering (and then uncovering) *Heliconia* plants in steep terrain with dense understories limited the area we could manipulate.

We also acknowledge that our ability to detect subtle experimental effects may have been limited by low statistical power, owing to high individual-level variation and limited sample size. Nevertheless, we can rule out extreme declines in hummingbird space use, flower visitation, and pollination success that would be indicative of complete network collapse; in the main analyses, treatment replicates generally showed no more than a 50% pre-to-post decline compared to control replicates (based on the lower confidence interval of the relevant contrast, e.g., Fig. 3A-B, Fig. 4A, C). We also emphasize that the spatial scale of our experiment and amount of replication surpasses all but two empirical studies simulating species extinctions within plant-pollinator networks (Appendix S1: Table S1). Still, future removal experiments with highly mobile pollinators should consider increasing the amount of replication, and, if possible, manipulate larger areas.

## CONCLUSIONS

Hummingbird pollinators seemed able to persist during our experiments that removed floral resources for a short period of time (several days), but we caution that this persistence may not be sustainable or reflect the outcome of a natural extinction process. Longer-term experiments will be key for predicting consequences of species loss and understanding mechanisms of network robustness – at least for pollinator species that forage at large spatial scales. Our study also suggests that behavioral flexibility to use alternative floral resources might be limited, even in taxonomic groups thought to be highly opportunistic foragers. Thus, even rewiring rules with strong biological underpinnings (e.g., Vizentin-Bugoni *et al*. 2020) may not necessarily predict real-world responses to species loss, perhaps due to additional constraints such as competition. To further understand the mechanisms underlying network robustness or collapse, we encourage future experiments that consider how temporal scale, species traits (e.g., movement ability), competition, and landscape context interact to determine the outcome of species extinctions. These experiments could be especially valuable at geographic range limits, where climate change will likely lead to species replacements (HilleRisLambers *et al*. 2013), and in the tropics where landscape change and species loss are most rapid (Hansen *et al*. 2013).

## ACKNOWLEDGMENTS

This material is based upon work supported by the National Science Foundation (NSF) Graduate Research Fellowship Program under Grant No. 1840998 to KGL. Funding for the experiment was provided primarily by NSF-DEB-1457837 to MGB and ASH, with additional grants to KGL from the American Ornithological Society and Sigma Xi. We also thank the Las Cruces Biological Station and the private landowners who allowed us to work on their property. Field assistance was provided by Jorge Araya Paniagua, Michael Atencio Picado, Marion Donald, Dustin Gannon, Jessica Greer, Matt Hadley, Urs Kormann, Daniel Moreno, Mauricio Paniagua Castro, Ignacio Ruiz Ilama, Esteban Sandi Paniagua, Andrés Solorzano Vargas, Laura Sutcliffe, and Felipe Torres Vanegas. Videos from trail cameras were reviewed with the help of Marion Donald, Colleen Franklin, Jessica Greer, Zoe Griffith, Elena Hart, Billy Hilgert, Zachary Kendall, Jeremy Lee, Ana Medina Roman, Briley Mullin, Amber Newell, Thanh Nguyen, Zoe Sallada, Allison Simoni, Andrew Stokes, Claire Woods, and Mel Xiao. Nozomi Birkett and Marion Donald helped collect nectar measurements, and Chandler Spruill assisted with pollen tube processing. Mark Novak and Berry Brosi provided feedback on an earlier version of the manuscript, and Ariel Muldoon provided valuable statistical advice. We also thank Nickolas Waser, Justin Bain, and three anonymous reviewers for their suggestions.

## REFERENCES

Allen-Wardell, G., P. Bernhardt, R. Bitner, A. Burquez, S. Buchmann, J. Cane, P. A. Cox, V. Dalton, P. Feinsinger, M. Ingram, D. Inouye, C. E. Jones, K. Kennedy, P. Kevan, H. Koopowitz, R. Medellin, S. Medellin-Morales, G. P. Nabhan, B. Pavlik, V. Tepedino, P. Torchio, and S. Walker. 1998. The potential consequences of pollinator declines on the conservation of biodiversity and stability of food crop yields. Conservation Biology 12:8–17.

Anderson, S. H., D. Kelly, J. J. Ladley, S. Molloy, and J. Terry. 2011. Cascading effects of bird functional extinction reduce pollination and plant density. Science 331:1068–1071.

Andrén, H. 1994. Effects of habitat fragmentation on birds and mammals in landscapes with different proportions of suitable habitat: A review. Oikos 71:355–366.

Arizmendi, M. C., and J. F. Ornelas. 1990. Hummingbirds and their floral resources in a tropical dry forest in Mexico. Biotropica 22:172–180.

Ashman, T. L., T. M. Knight, J. A. Steets, P. Amarasekare, M. Burd, D. R. Campbell, M. R. Dudash, M. O. Johnston, S. J. Mazer, and R. J. Mitchell. 2004. Pollen limitation of plant reproduction: ecological and evolutionary causes and consequences. Ecology 85:2408– 2421.

Aslan, C. E., E. S. Zavaleta, B. Tershy, and D. Croll. 2013. Mutualism disruption threatens global plant biodiversity: A systematic review. PLOS ONE 8:e66993.

Bascompte, J., and P. Jordano. 2007. Plant-animal mutualistic networks: the architecture of biodiversity. Annual Review of Ecology, Evolution, and Systematics 38:567–593.

Bascompte, J., P. Jordano, C. J. Melián, and J. M. Olesen. 2003. The nested assembly of plant– animal mutualistic networks. Proceedings of the National Academy of Sciences 100:9383–9387.

Bascompte, J., P. Jordano, and J. M. Olesen. 2006. Asymmetric coevolutionary networks facilitate biodiversity maintenance. Science 312:431–433.

Bech, C., A. S. Abe, J. F. Steffensen, M. Berger, and J. E. P. W. Bicudo. 1997. Torpor in three species of Brazilian hummingbirds under semi-natural conditions. The Condor 99:780– 788.

Betts, M. G., A. S. Hadley, and W. J. Kress. 2015. Pollinator recognition by a keystone tropical plant. Proceedings of the National Academy of Sciences 112:3433–3438.

Biesmeijer, J. C., S. P. M. Roberts, M. Reemer, R. Ohlemüller, M. Edwards, T. Peeters, A. P. Schaffers, S. G. Potts, R. Kleukers, C. D. Thomas, J. Settele, and W. E. Kunin. 2006. Parallel declines in pollinators and insect-pollinated plants in Britain and the Netherlands. Science 313:351–354.

Borgella, R., A. A. Snow, and T. A. Gavin. 2001. Species richness and pollen loads of hummingbirds using forest fragments in southern Costa Rica. Biotropica 33:90–109.

Brice, A. T. 1992. The essentiality of nectar and arthropods in the diet of the Anna’s hummingbird (Calypte anna). Comparative Biochemistry and Physiology Part A: Physiology 101:151–155.

Brooks, M. E., K. Kristensen, K. J. van Benthem, A. Magnusson, C. W. Berg, A. Nielsen, H. J. Skaug, M. Maechler, and B. M. Bolker. 2017. glmmTMB Balances Speed and Flexibility Among Packages for Zero-inflated Generalized Linear Mixed Modeling. The R Journal 9:378–400.

Brosi, B. J., and H. M. Briggs. 2013. Single pollinator species losses reduce floral fidelity and plant reproductive function. Proceedings of the National Academy of Sciences 110:13044–13048.

Bruna, E. M., and W. J. Kress. 2002. Habitat fragmentation and the demographic structure of an Amazonian understory herb (Heliconia acuminata). Conservation Biology 16:1256– 1266.

Calder, W. A., and J. Booser. 1973. Hypothermia of broad-tailed hummingbirds during incubation in nature with ecological correlations. Science 180:751–753.

Carpenter, F. L., and M. A. Hixon. 1988. A new function for torpor: fat conservation in a wild migrant hummingbird. The Condor 90:373–378.

Colwell, R. K., R. R. Dunn, and N. C. Harris. 2012. Coextinction and persistence of dependent species in a changing world. Annual Review of Ecology, Evolution, and Systematics 43:183–203.

Costa, J. M., J. A. Ramos, L. P. da Silva, S. Timóteo, P. Andrade, P. M. Araújo, C. Carneiro, E. Correia, P. Cortez, M. Felgueiras, C. Godinho, R. J. Lopes, C. Matos, A. C. Norte, P. F. Pereira, A. Rosa, and R. H. Heleno. 2018. Rewiring of experimentally disturbed seed dispersal networks might lead to unexpected network configurations. Basic and Applied Ecology 30:11–22.

Dunn, R. R., N. C. Harris, R. K. Colwell, L. P. Koh, and N. S. Sodhi. 2009. The sixth mass coextinction: are most endangered species parasites and mutualists? Proceedings of the Royal Society B: Biological Sciences 276:3037–3045.

Eberhard, J. R., and P. W. Ewald. 1994. Food availability, intrusion pressure and territory size: an experimental study of Anna’s hummingbirds (Calypte anna). Behavioral Ecology and Sociobiology 34:11–18.

Feinsinger, P., and R. K. Colwell. 1978. Community organization among Neotropical nectarfeeding birds. American Zoologist 18:779–795.

Ferrero, V., S. Castro, J. Costa, P. Acuña, L. Navarro, and J. Loureiro. 2013. Effect of invader removal: pollinators stay but some native plants miss their new friend. Biological Invasions 15:2347–2358.

Gass, C. L., and G. D. Sutherland. 1985. Specialization by territorial hummingbirds on experimentally enriched patches of flowers: energetic profitability and learning. Canadian Journal of Zoology 63:2125–2133.

Gilbert, L. E. 1980. Food web organization and conservation of neotropical diversity. Pages 11– 33 in M. E. Soulé and B. A. Wilcox, editors. Conservation Biology: An Evolutionary-Ecological Perspective. Sinauer, Sunderland.

Gill, F. B. 1988. Trapline foraging by hermit hummingbirds: competition for an undefended, renewable resource. Ecology 69:1933–1942.

Goldstein, J., and M. Zych. 2016. What if we lose a hub? Experimental testing of pollination network resilience to removal of keystone floral resources. Arthropod-Plant Interactions 10:263–271.

Hadley, A. S., and M. G. Betts. 2009. Tropical deforestation alters hummingbird movement patterns. Biology Letters 5:207–210.

Hadley, A. S., and M. G. Betts. 2012. The effects of landscape fragmentation on pollination dynamics: absence of evidence not evidence of absence. Biological Reviews 87:526–544.

Hadley, A. S., S. J. K. Frey, W. D. Robinson, and M. G. Betts. 2018. Forest fragmentation and loss reduce richness, availability, and specialization in tropical hummingbird communities. Biotropica 50:74–83.

Hadley, A. S., S. J. K. Frey, W. D. Robinson, W. J. Kress, and M. G. Betts. 2014. Tropical forest fragmentation limits pollination of a keystone understory herb. Ecology 95:2202–2212.

Hainsworth, F. R., B. G. Collins, and L. L. Wolf. 1977. The function of torpor in hummingbirds. Physiological Zoology 50:215–222.

Hansen, M. C., P. V. Potapov, R. Moore, M. Hancher, S. A. Turubanova, A. Tyukavina, D. Thau, S. V. Stehman, S. J. Goetz, T. R. Loveland, A. Kommareddy, A. Egorov, L. Chini, C. O. Justice, and J. R. G. Townshend. 2013. High-resolution global maps of 21st-century forest cover change. Science 342:850–853.

Hartig, F. 2020. DHARMa: Residual Diagnostics for Hierarchical (Multi-Level / Mixed) Regression Models.

Hazlehurst, J. A., and J. O. Karubian. 2018. Impacts of nectar robbing on the foraging ecology of a territorial hummingbird. Behavioural Processes 149:27–34.

Heinrich, B., and P. H. Raven. 1972. Energetics and pollination ecology. Science 176:597–602.

HilleRisLambers, J., M. A. Harsch, A. K. Ettinger, K. R. Ford, and E. J. Theobald. 2013. How will biotic interactions influence climate change–induced range shifts? Annals of the New York Academy of Sciences 1297:112–125.

Hixon, M. A., F. L. Carpenter, and D. C. Paton. 1983. Territory area, flower density, and time budgeting in hummingbirds: an experimental and theoretical analysis. The American Naturalist 122:366–391.

Jordán, F. 2009. Keystone species and food webs. Philosophical Transactions of the Royal Society B: Biological Sciences 364:1733–1741.

Kaiser-Bunbury, C. N., J. Mougal, A. E. Whittington, T. Valentin, R. Gabriel, J. M. Olesen, and N. Blüthgen. 2017. Ecosystem restoration strengthens pollination network resilience and function. Nature 542:223–227.

Kaiser-Bunbury, C. N., S. Muff, J. Memmott, C. B. Müller, and A. Caflisch. 2010. The robustness of pollination networks to the loss of species and interactions: a quantitative approach incorporating pollinator behaviour. Ecology Letters 13:442–452.

Kearns, C. A., and D. W. Inouye. 1993. Techniques for Pollination Biologists. University Press of Colorado, Niwot.

Klein, A.-M., B. E. Vaissière, J. H. Cane, I. Steffan-Dewenter, S. A. Cunningham, C. Kremen, and T. Tscharntke. 2007. Importance of pollinators in changing landscapes for world crops. Proceedings of the Royal Society B: Biological Sciences 274:303–313.

Koh, L. P., R. R. Dunn, N. S. Sodhi, R. K. Colwell, H. C. Proctor, and V. S. Smith. 2004. Species Coextinctions and the Biodiversity Crisis. Science 305:1632–1634.

Kormann, U., C. Scherber, T. Tscharntke, N. Klein, M. Larbig, J. J. Valente, A. S. Hadley, and M. G. Betts. 2016. Corridors restore animal-mediated pollination in fragmented tropical forest landscapes. Proceedings of the Royal Society B: Biological Sciences 283:20152347.

Krüger, K., R. Prinzinger, and K.-L. Schuchmann. 1982. Torpor and metabolism in hummingbirds. Comparative Biochemistry and Physiology Part A: Physiology 73:679– 689.

Kuban, J. F., and R. L. Neill. 1980. Feeding ecology of hummingbirds in the highlands of the Chisos Mountains, Texas. The Condor 82:180–185.

Kuussaari, M., R. Bommarco, R. K. Heikkinen, A. Helm, J. Krauss, R. Lindborg, E. Öckinger, M. Pärtel, J. Pino, F. Roda, and others. 2009. Extinction debt: a challenge for biodiversity conservation. Trends in ecology & evolution 24:564–571.

Lenth, R. 2020. emmeans: Estimated Marginal Means, aka Least-Squares Means.

Lüdecke, D., M. Ben-Shachar, I. Patil, P. Waggoner, and D. Makowski. 2021. performance: An R Package for Assessment, Comparison and Testing of Statistical Models. Journal of Open Source Software 6:3139.

Maruyama, P. K., G. M. Oliveira, C. Ferreira, B. Dalsgaard, and P. E. Oliveira. 2013. Pollination syndromes ignored: importance of non-ornithophilous flowers to Neotropical savanna hummingbirds. Naturwissenschaften 100:1061–1068.

McDonald, T. L., W. P. Erickson, and L. L. McDonald. 2000. Analysis of count data from before-after control-impact studies. Journal of Agricultural, Biological, and Environmental Statistics 5:262–279.

Memmott, J., N. M. Waser, and M. V. Price. 2004. Tolerance of pollination networks to species extinctions. Proceedings of the Royal Society B: Biological Sciences 271:2605–2611.

Mendenhall, C. D., and A. M. Wrona. 2018. Improving tree cover estimates for fine-scale landscape ecology. Landscape Ecology 33:1691–1696.

Montgomerie, R. D., and C. A. Redsell. 1980. A nesting hummingbird feeding solely on arthropods. The Condor 82:463.

Moran, A. J., S. W. J. Prosser, and J. A. Moran. 2019. DNA metabarcoding allows non-invasive identification of arthropod prey provisioned to nestling Rufous hummingbirds (Selasphorus rufus). PeerJ 7:e6596.

Ohashi, K., and J. D. Thomson. 2009. Trapline foraging by pollinators: its ontogeny, economics and possible consequences for plants. Annals of Botany 103:1365–1378.

Ollerton, J., R. Winfree, and S. Tarrant. 2011. How many flowering plants are pollinated by animals? Oikos 120:321–326.

Pauw, A., and J. A. Hawkins. 2011. Reconstruction of historical pollination rates reveals linked declines of pollinators and plants. Oikos 120:344–349.

Phillips, R. D., R. Peakall, B. A. Retter, K. Montgomery, M. H. M. Menz, B. J. Davis, C. Hayes, G. R. Brown, N. D. Swarts, and K. W. Dixon. 2015. Pollinator rarity as a threat to a plant with a specialized pollination system. Botanical Journal of the Linnean Society 179:511– 525.

Potts, S. G., J. C. Biesmeijer, C. Kremen, P. Neumann, O. Schweiger, and W. E. Kunin. 2010. Global pollinator declines: trends, impacts and drivers. Trends in Ecology & Evolution 25:345–353.

Powers, D. R., A. R. Brown, and J. A. Van Hook. 2003. Influence of normal daytime fat deposition on laboratory measurements of torpor use in territorial versus nonterritorial hummingbirds. Physiological and Biochemical Zoology 76:389–397.

R Core Team. 2022. R: A Language and Environment for Statistical Computing. R Foundation for Statistical Computing, Vienna, Austria.

Ramos-Jiliberto, R., F. S. Valdovinos, P. Moisset de Espanés, and J. D. Flores. 2012. Topological plasticity increases robustness of mutualistic networks. Journal of Animal Ecology 81:896–904.

Regan, E. C., L. Santini, L. Ingwall□King, M. Hoffmann, C. Rondinini, A. Symes, J. Taylor, and S. H. M. Butchart. 2015. Global trends in the status of bird and mammal pollinators. Conservation Letters 8:397–403.

Rodríguez-Flores, C. I., J. F. Ornelas, S. Wethington, and M. C. Arizmendi. 2019. Are hummingbirds generalists or specialists? Using network analysis to explore the mechanisms influencing their interaction with nectar resources. PLOS ONE 14:e0211855.

Russell, S. M., and R. O. Russell. 2001. The North American Banders’ Manual for Banding Hummingbirds. North American Banding Council, Point Reyes Station, CA.

Scheper, J., M. Reemer, R. van Kats, W. A. Ozinga, G. T. J. van der Linden, J. H. J. Schaminée, H. Siepel, and D. Kleijn. 2014. Museum specimens reveal loss of pollen host plants as key factor driving wild bee decline in The Netherlands. Proceedings of the National Academy of Sciences 111:17552–17557.

Schuchmann, K.-L., and R. Prinzinger. 1988. Energy metabolism, nocturnal torpor, and respiration frequency in a Green Hermit (Phaethornis guy). Journal für Ornithologie 129:469–472.

Shankar, A., R. J. Schroeder, S. M. Wethington, C. H. Graham, and D. R. Powers. 2020. Hummingbird torpor in context: duration, more than temperature, is the key to nighttime energy savings. Journal of Avian Biology 51:e02305.

Spence, A. R., and M. W. Tingley. 2021. Body size and environment influence both intraspecific and interspecific variation in daily torpor use across hummingbirds. Functional Ecology 35:870–883.

Stiles, F. G. 1975. Ecology, flowering phenology, and hummingbird pollination of some Costa Rican Heliconia species. Ecology 56:285–301.

Stiles, F. G. 1979. Notes on the natural history of Heliconia (Musaceae) in Costa Rica. Notas sobre la historia natural de Heliconia (Musaceae) en Costa Rica. Brenesia 15:151–180.

Stiles, F. G. 1985. Seasonal patterns and coevolution in the hummingbird-flower community of a Costa Rican subtropical forest. Ornithological Monographs 36:757–787.

Stiles, F. G. 1995. Behavioral, ecological and morphological correlates of foraging for arthropods by the hummingbirds of a tropical wet forest. The Condor 97:853–878.

Stiles, F. G., and L. L. Wolf. 1970. Hummingbird territoriality at a tropical flowering tree. The Auk 87:467–491.

Stratton, D. A. 1989. Longevity of individual flowers in a Costa Rican cloud forest: ecological correlates and phylogenetic constraints. Biotropica 21:308–318.

Suarez, R. K. 1992. Hummingbird flight: sustaining the highest mass-specific metabolic rates among vertebrates. Experientia 48:565–570.

Taylor, J., and S. A. White. 2007. Observations of hummingbird feeding behavior at flowers of Heliconia beckneri and H. tortuosa in southern Costa Rica. Ornitologia Neotropical 18:133–138.

Temeles, E. J., A. B. Muir, E. B. Slutsky, and M. N. Vitousek. 2004. Effect of food reductions on territorial behavior of purple-throated caribs. The Condor 106:691–695.

Thomas-Granger, K. 2003. Light sum, nitrogen level and growth retardant effects on flower and nectar production in Stachytarpheta frantzii. Master’s thesis, University of Guelph.

Timóteo, S., J. A. Ramos, I. P. Vaughan, and J. Memmott. 2016. High resilience of seed dispersal webs highlighted by the experimental removal of the dominant disperser. Current Biology 26:910–915.

Tooze, Z. J., and C. L. Gass. 1985. Responses of rufous hummingbirds to midday fasts. Canadian Journal of Zoology 63:2249–2253.

Traveset, A., C. Tur, and V. M. Eguíluz. 2017. Plant survival and keystone pollinator species in stochastic coextinction models: role of intrinsic dependence on animal-pollination. Scientific Reports 7:6915.

Underwood, A. J. 1994. On Beyond BACI: sampling designs that might reliably detect environmental disturbances. Ecological Applications 4:3–15.

Valdovinos, F. S., P. Moisset de Espanés, J. D. Flores, and R. Ramos-Jiliberto. 2013. Adaptive foraging allows the maintenance of biodiversity of pollination networks. Oikos 122:907– 917.

Vellend, M., K. Verheyen, H. Jacquemyn, A. Kolb, H. V. Calster, G. Peterken, and M. Hermy. 2006. Extinction debt of forest plants persists for more than a century following habitat fragmentation. Ecology 87:542–548.

Vizentin-Bugoni, J., V. J. Debastiani, V. A. G. Bastazini, P. K. Maruyama, and J. H. Sperry. 2020. Including rewiring in the estimation of the robustness of mutualistic networks. Methods in Ecology and Evolution 11:106–116.

Volpe, N. L., A. S. Hadley, W. D. Robinson, and M. G. Betts. 2014. Functional connectivity experiments reflect routine movement behavior of a tropical hummingbird species. Ecological Applications 24:2122–2131.

Volpe, N. L., W. D. Robinson, S. J. K. Frey, A. S. Hadley, and M. G. Betts. 2016. Tropical forest fragmentation limits movements, but not occurrence of a generalist pollinator species. PLOS ONE 11:e0167513.

Weiner, C. N., M. Werner, K. E. Linsenmair, and N. Blüthgen. 2014. Land-use impacts on plant–pollinator networks: interaction strength and specialization predict pollinator declines. Ecology 95:466–474.

Weinstein, B. G. 2015. MotionMeerkat: integrating motion video detection and ecological monitoring. Methods in Ecology and Evolution 6:357–362.

Weinstein, B. G., and C. H. Graham. 2017. Persistent bill and corolla matching despite shifting temporal resources in tropical hummingbird-plant interactions. Ecology Letters 20:326– 335.

Wolf, L. L., and F. G. Stiles. 1989. Adaptations for the “fail-safe” pollination of specialized ornithophilous flowers. The American Midland Naturalist 121:1–10.

Young, A. M. 1971. Foraging for insects by a tropical hummingbird. The Condor 73:36–45.

Zahawi, R. A., G. Duran, and U. Kormann. 2015. Sixty-seven years of land-use change in southern Costa Rica. PLOS ONE 10:e0143554.

