## Appendix S1 for "Tropical hummingbird pollination networks are resistant to short-term experimental removal of a common flowering plant"

**Appendix S1:** Compilation of previous removal experiments with mutualistic networks

Supporting Information for:

**Tropical hummingbird pollination networks are resistant to short-term experimental removal of a common flowering plant**

Kara G. Leimberger^1^, Adam S. Hadley^1,2^, Sarah J.K. Frey^1,3^, and Matthew G. Betts^1^

^1^Forest Biodiversity Research Network, Department of Forest Ecosystems and Society, Oregon State University, Corvallis, Oregon, USA

^2^Biodiversity Section, Department of Natural Resources and Energy Development, Fredericton, New Brunswick, Canada

^3^Department of Animal and Rangeland Sciences, Oregon State University, Corvallis, Oregon, USA

**Table S1.** Summary of previous studies in which animals or plants were experimentally removed from mutualistic networks to test predictions about network robustness.

BACI refers to ‘Before-After-Control-Impact’, where ‘Impact’ references the Treatment (i.e., species removal). ‘Spatial scale’ refers to the scale at which the manipulation was implemented.

|  | **Study** | **Network type** | **Location** | **Experimental design** | **#**  **treatment replicates** | **#**  **control replicates** | **Species removed** | **Spatial scale (hectares)** | **Temporal scale** |
| --- | --- | --- | --- | --- | --- | --- | --- | --- | --- |
| **Animal removals** | Brosi & Briggs (2013) | Pollination  (plant–insect) | Gothic, Colorado USA | Before-After | 20 |  | Variable; most abundant *Bombus* sp. | 0.04 | 1 day |
|  | Brosi *et al.* (2017) | Pollination  (plant–insect) | Gothic, Colorado USA | Before-After | 15 |  | Variable; most abundant *Bombus* sp. | 0.04 | 1 day |
|  | Timóteo *et al.* (2016) | Seed dispersal  (plant–ant) | Portugal | Control-Impact^1^ | 18 | 18 | *Messor barbatus* | 0.01 | ~90 days^2^ |
| **Plant removals** | Lopezaraiza-Mikel *et al.* (2007) | Pollination  (plant–insect) | Bristol, Great Britain | Control-Impact^3^ | 8 | 8 | *Impatiens glandulifera*  (non-native) | 0.04 | ~90 days |
|  | Ferrero *et al.* (2013) | Pollination  (plant–insect) | Castelo Viegas, Portugal | Before-After | 1 |  | *Oxalis pes-caprae*  (non-native) | 0.05 | ~4 days before,  ~4 days after |
|  | Goldstein & Zych (2016) | Pollination  (plant–insect) | Kleczkowo, Poland | BACI | 1 | 1 | *Polygonum bistorta* | 0.5 | 2 days before,  2 days after |
|  | Kaiser-Bunbury *et al.* (2017) | Pollination  (plant–insect, primarily) | Mahé, Seychelles | Control-Impact | 2 | 2 | Variable; all non-native species | ~1 | ~240 days |
|  | Costa *et al.* (2018) | Seed dispersal  (plant–bird) | Portugal | Before-After | 1 |  | *Rubus ulmifolius* | 3.14 | 7 days before,  7 days after |
|  | Biella *et al.* (2019)  Biella *et al.* (2020) | Pollination  (plant–insect) | Český Krumlov, Czech Republic | BACI | 3 | 1 | Variable; sequential removal of four most generalist plant species | 0.15 - 0.18 | 2 days before,  2 days after |
|  | Bain *et al.*  (2022) | Pollination  (plant–insect) | Gothic, Colorado USA | Control-Impact | 10 | 10 | *Helianthella quinquenervis* | 0.017 | 21 days |
|  | **Present study** | Pollination  (plant–hummingbird) | San Vito, Costa Rica | BACI  14 sites | 16 | 16 | *Heliconia tortuosa* | Variable;  0.27 - 7.49  Median: 1.25 | 4 days before,  4 days after |

1, 1 site (1700 ha farm), 3 habitat types, 12 plots/habitat type

2, 5 weeks for removal and 6 weeks for data collection

3, 4 sites and 2 plots/site

**Literature cited**

Bain, J. A., R. G. Dickson, A. M. Gruver, and P. J. CaraDonna. 2022. Removing flowers of a generalist plant changes pollinator visitation, composition, and interaction network structure. Ecosphere 13:e4154.

Biella, P., A. Akter, J. Ollerton, A. Nielsen, and J. Klecka. 2020. An empirical attack tolerance test alters the structure and species richness of plant–pollinator networks. Functional Ecology 34:2246–2258.

Biella, P., A. Akter, J. Ollerton, S. Tarrant, Š. Janeček, J. Jersáková, and J. Klecka. 2019. Experimental loss of generalist plants reveals alterations in plant-pollinator interactions and a constrained flexibility of foraging. Scientific Reports 9:7376.

Brosi, B. J., and H. M. Briggs. 2013. Single pollinator species losses reduce floral fidelity and plant reproductive function. Proceedings of the National Academy of Sciences 110:13044–13048.

Brosi, B. J., K. Niezgoda, and H. M. Briggs. 2017. Experimental species removals impact the architecture of pollination networks. Biology Letters 13:20170243.

Costa, J. M., J. A. Ramos, L. P. da Silva, S. Timóteo, P. Andrade, P. M. Araújo, C. Carneiro, E. Correia, P. Cortez, M. Felgueiras, C. Godinho, R. J. Lopes, C. Matos, A. C. Norte, P. F. Pereira, A. Rosa, and R. H. Heleno. 2018. Rewiring of experimentally disturbed seed dispersal networks might lead to unexpected network configurations. Basic and Applied Ecology 30:11–22.

Ferrero, V., S. Castro, J. Costa, P. Acuña, L. Navarro, and J. Loureiro. 2013. Effect of invader removal: pollinators stay but some native plants miss their new friend. Biological Invasions 15:2347–2358.

Goldstein, J., and M. Zych. 2016. What if we lose a hub? Experimental testing of pollination network resilience to removal of keystone floral resources. Arthropod-Plant Interactions 10:263–271.

Kaiser-Bunbury, C. N., J. Mougal, A. E. Whittington, T. Valentin, R. Gabriel, J. M. Olesen, and N. Blüthgen. 2017. Ecosystem restoration strengthens pollination network resilience and function. Nature 542:223–227.

Lopezaraiza–Mikel, M. E., R. B. Hayes, M. R. Whalley, and J. Memmott. 2007. The impact of an alien plant on a native plant–pollinator network: an experimental approach. Ecology Letters 10:539–550.

Timóteo, S., J. A. Ramos, I. P. Vaughan, and J. Memmott. 2016. High resilience of seed dispersal webs highlighted by the experimental removal of the dominant disperser. Current Biology 26:910–915.
