## Appendix S2 for "Tropical hummingbird pollination networks are resistant to short-term experimental removal of a common flowering plant"

**Appendix S2:** Supplemental Methods (Study design, Response variables, Statistical analysis)

Supporting Information for:

**Tropical hummingbird pollination networks are resistant to short-term experimental removal of a common flowering plant**

Kara G. Leimberger^1^, Adam S. Hadley^1,2^, Sarah J.K. Frey^1,3^, and Matthew G. Betts^1^

^1^Forest Biodiversity Research Network, Department of Forest Ecosystems and Society, Oregon State University, Corvallis, Oregon, USA

^2^Biodiversity Section, Department of Natural Resources and Energy Development, Fredericton, New Brunswick, Canada

^3^Department of Animal and Rangeland Sciences, Oregon State University, Corvallis, Oregon, USA

### Study design

#### Table S1. Characteristics of the 14 sites used in this experiment. Site ID refers to the unique site code used by the Oregon State University Tropical Hummingbird Research Project. Elevation values were extracted for each study site from the NASA SRTM Digital Elevation Model (30-m) available in Google Earth Engine (Gorelick *et al.* 2017).

| Site number | Site ID | Elevation (m) | Connectivity to additional forest |
| --- | --- | --- | --- |
| 1 | 200 | 1437 | Low |
| 2 | 137 | 1165 | Low |
| 3 | 204 | 929 | Low |
| 4 | 130 | 1301 | Low |
| 5 | 60 | 1107 | Low |
| 6 | 205 | 1365 | Medium |
| 7 | 58 | 1405 | Medium |
| 8 | 203 | 1457 | Medium |
| 9 | 24 | 1354 | Medium |
| 10 | 10 | 1248 | High |
| 11 | 49 | 996 | High |
| 12 | 29 | 1333 | High |
| 13 | 30 | 1236 | High |
| 14 | 201 | 1187 | High |

#### Table S2. Sample size per analysis. For all metrics except interaction turnover, values were calculated for both experimental periods (pre and post). For example, the analysis of individual specialization (recaptures) included 27 individuals each sampled twice, yielding 54 total observations.

| Analysis | Subanalysis | Bird group | # observations (rows) | # treatment replicates | # control replicates |
| --- | --- | --- | --- | --- | --- |
| Mist net captures | Captures | All species | 59 | 15 | 15 |
|  |  | *Heliconia* specialists | 56 | 15 | 15 |
|  | Recaptures | All species | 28 | 14 | 14 |
|  |  | *Heliconia* specialists | 25 | 12 | 13 |
| Radio telemetry | All replicates | All species | 72 | 10 | 14 |
|  |  | *Heliconia* specialists | 54 | 8 | 14 |
|  | Without outlier replicate | All species | 64 | 10 | 13 |
|  |  | *Heliconia* specialists | 48 | 8 | 13 |
| Flower visitation (*Heliconia*) | All birds | All species | 62 | 15 | 16 |
|  |  | *Heliconia* specialists | 62 | 15 | 16 |
|  | Marked birds | All species | 52 | 8 | 8 |
|  |  | *Heliconia* specialists | 42 | 6 | 7 |
| Flower visitation  (non-*Heliconia*) | All birds | All species | 420 | 16 | 16 |
|  |  | *Heliconia* specialists | 420 | 16 | 16 |
|  | Marked birds | All species | 160 | 7 | 8 |
|  |  | *Heliconia* specialists | 90 | 5 | 6 |
| Flower visitation  (non-*Heliconia,*  individual plant species)  All birds | *Calathea crotalifera* | All species | 48 | 11 | 13 |
|  | *Scutellaria costaricana* | All species | 44 | 11 | 11 |
|  | *Centropogon granulosus* | All species | 42 | 10 | 11 |
|  | *Columnea polyantha* | All species | 36 | 9 | 9 |
|  | *Hamelia patens* | All species | 34 | 8 | 9 |
|  | *Pachystachys lutea* | All species | 32 | 9 | 7 |
|  | *Stachytarpheta frantzii* | All species | 30 | 7 | 8 |
|  | *Columnea raymondii* | All species | 28 | 8 | 6 |
|  | *Costus barbatus* | All species | 22 | 5 | 6 |
| Pollination success | *Hamelia patens* | NA | 24 | 7 | 5 |
|  | *Heliconia* | NA | 50 | 13 | 12 |
| Body condition (mass) |  | All species | 60 | 9 | 9 |
|  |  | *Heliconia* specialists | 32 | 6 | 5 |

#### Table S3. Site pairings and treatment assignments for the *Heliconia* removal experiment. Colors indicate experimental order and site pairings within each year (red = first replicate of field season, blue = last replicate of field season). Despite the paired design, data were not analyzed within the paired framework since data were sometimes unavailable for one site within a pair.

| Replicate | Treatment assignment |
| --- | --- |
| 2016_137 | Treatment |
| 2016_60 | Control |
| 2016_10 | Treatment |
| 2016_201 | Control |
| 2016_58 | Treatment |
| 2016_24 | Control |
| 2016_130 | Treatment |
| 2016_200 | Control |
| 2016_30 | Treatment |
| 2016_203 | Control |
| 2016_204 | Treatment |
| 2016_49 | Control |
| 2017_10 | Control |
| 2017_201 | Treatment |
| 2017_60 | Treatment |
| 2017_137 | Control |
| 2017_24 | Treatment |
| 2017_58 | Control |
| 2017_130 | Control |
| 2017_200 | Treatment |
| 2017_30 | Control |
| 2017_203 | Treatment |
| 2017_49 | Treatment |
| 2017_204 | Control |
| 2018_29 | Control |
| 2018_205 | Treatment |
| 2018_60 | Treatment |
| 2018_137 | Control |
| 2018_10 | Treatment |
| 2018_58 | Control |
| 2018_24 | Control |
| 2018_30 | Treatment |

#### Table S4. Variation in amount of *H. tortuosa* removed in treatment replicates. We not only covered flowering *Heliconia*, but also plants that might flower within the experimental period (i.e., plants with immature bracts). Flowers and calories were estimated based on the number of open bracts. Estimates of calories are based on nectar measurements from covered flowers, i.e., not accounting for nectar refill following hummingbird visitation; the values presented here can thus be considered a minimum estimate of calories depleted. Values are ordered by the number of calories removed per hectare, calculated based on the size of the focal area.

|  |  | Resources removed | | | | | |  |
| --- | --- | --- | --- | --- | --- | --- | --- | --- |
| Year | Site | Total inflorescences | Inflorescences with open bracts^a^ | Inflorescences with ≥3 open bracts^b^ | Flowers (estimated) | Calories (estimated)^c^ | Calories per ha  (estimated) | Size of focal area (ha)^d^ |
| 2017 | 203 | 8 | 6 | 6 | 6 | 330 | 134 | 2.47 |
| 2017 | 201 | 81 | 66 | 36 | 36 | 1983 | 265 | 7.49 ^e^ |
| 2016 | 204 | 66 | 60 | 45 | 53 | 2919 | 646 | 4.52 |
| 2017 | 60 | 21 | 16 | 9 | 9 | 496 | 1510 | 0.33 |
| 2016 | 130 | 82 | 66 | 37 | 42 | 2313 | 1649 | 1.4 |
| 2017 | 24 | 149 | 104 | 56 | 61 | 3360 | 2650 | 1.27 |
| 2017 | 200 | 182 | 114 | 62 | 64 | 3525 | 3092 | 1.14 |
| 2018 | 10 | 370 | 281 | 210 | 260 | 14320 | 5966 | 2.4 |
| 2016 | 58 | 269 | 187 | 127 | 149 | 8206 | 6083 | 1.35 |
| 2017 | 49 | 241 | 189 | 126 | 152 | 8372 | 6848 | 1.22 |
| 2016 | 10 | 290 | 244 | 190 | 234 | 12888 | 7526 | 1.71 |
| 2016 | 30 | 127 | 115 | 77 | 90 | 4957 | 8475 | 0.58 |
| 2016 | 137 | 104 | 89 | 64 | 76 | 4186 | 8801 | 0.48 |
| 2018 | 60 | 88 | 70 | 44 | 48 | 2644 | 9853 | 0.27 |
| 2018 | 30 | 154 | 112 | 83 | 124 | 6829 | 19616 | 0.35 |
| 2018 | 205 | 526 | 264 | 166 | 173 | 9528 | 20480 | 0.47 |
| ***Min*** | | 8 | 6 | 6 | 6 | 330 | 134 | 0.27 |
| ***Max*** | | 526 | 281 | 210 | 260 | 14320 | 20480 | 7.49 |
| ***Median*** | | 138 | 108 | 63 | 70 | 3856 | 6025 | 1.25 |
| ***Mean ± SD*** | | 172 ± 138 | 124 ± 85 | 84 ± 62 | 99 ± 76 | 5421 ± 4194 | 6471 ± 6208 | 1.7 ± 1.9 |

a, *Heliconia* inflorescences have sequentially opening bracts, which produce up to one flower per day once open. Thus, open bracts indicate that the plant was flowering prior to removal.

b, *Heliconia* plants with fewer than three bracts were not assigned any flowers.

c, Unit is calories (not kilocalories).

d, Area in which *Heliconia* plants were covered, delineated by a minimum convex polygon created from all plant locations recorded during resource survey conducted alongside *Heliconia* covering.

e, Site 201 contained a small lake (0.8 ha) within the focal area; lake is excluded from this value.

### Response variables

**Mist net captures**

***Capture effort***

We calculated ‘net-hours’ as the number of nets multiplied by the number of hours it was open, with 6-m nets counted as half a net. Pre-to-post differences in net-hours occasionally arose if nets were closed early due to logistical or safety concerns (e.g., potential predator in the area). Overall, we found no evidence that capture effort differed between capture sessions (Fig. S2; paired Wilcoxon test: V = 201.5, *P* = 0.77). Nevertheless, before analyzing recapture probability we removed the treatment replicate with substantially more effort during the ‘post’ period.

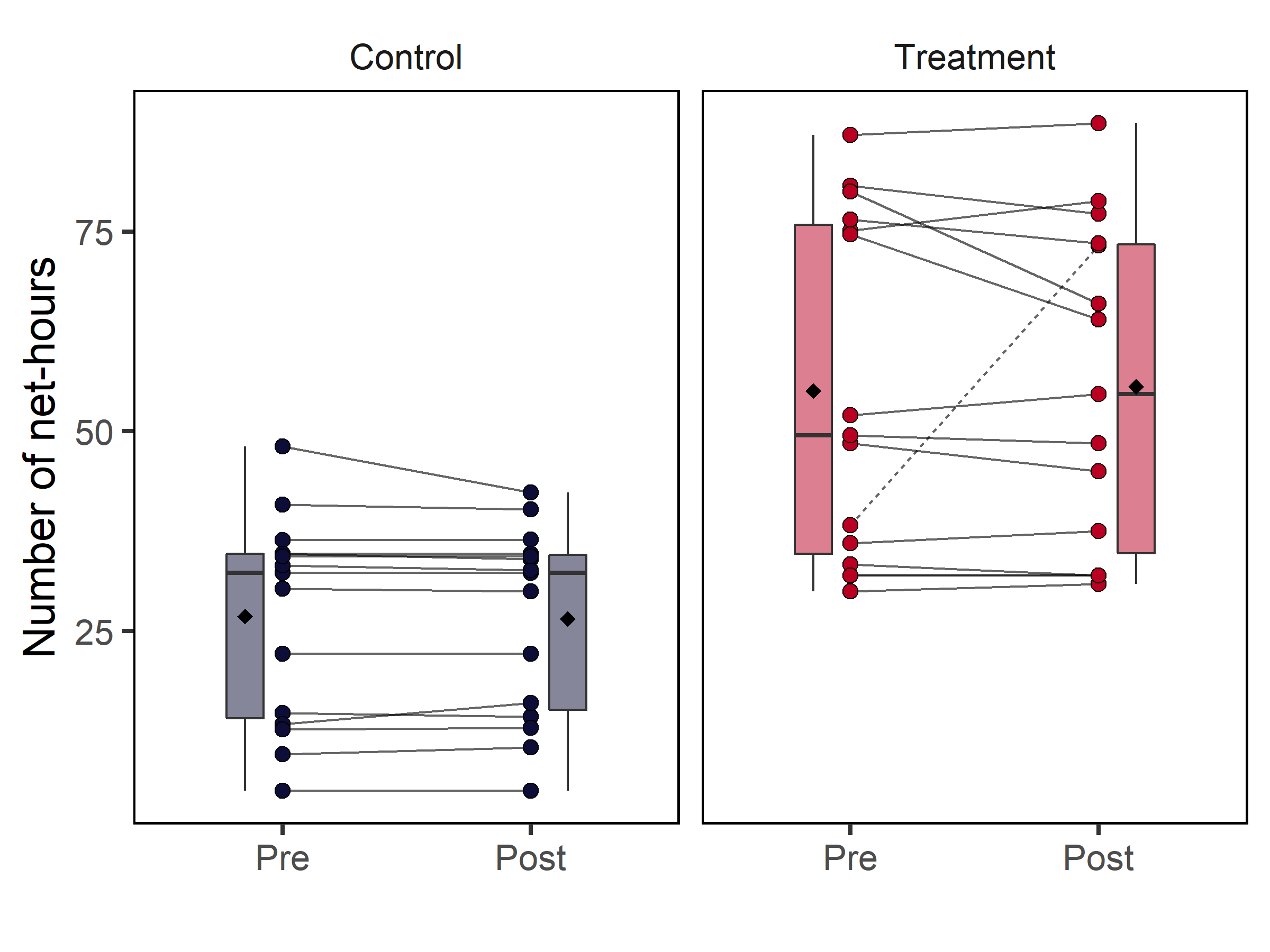

#### Figure S1. Comparisons of capture effort between pre and post capture sessions. Boxplots are presented alongside raw data points, with pre-to-post pairings indicated by solid lines. Black diamonds indicate mean values, while darker black lines indicate medians. Lower and upper lines represent first and third quartiles; whiskers extend to the lowest and highest values within 1.5x the IQR. The one replicate with dramatic pre-to-post differences in capture effort (dashed line) was not included in analysis of recapture probability.

**Radio telemetry**

***Radio transmitter attachment***

Following Hadley & Betts (2009) and Volpe *et al.* (2014, 2016), we attached transmitters with eyelash glue after parting the feathers of the lower back to expose bare skin. Transmitters attached using this technique are estimated to remain on birds for approximately 14 days (Hadley and Betts 2009), which corresponds to the transmitter’s maximum expected battery life.

***Telemetry observations***

Following Hadley & Betts (2009) and Volpe *et al.* (2014, 2016), we recorded locations continuously when the bird was found within receiver range. For each bird location, we recorded the bird’s arrival time, the observer’s GPS coordinates, the bird’s estimated distance from the observer, and the bird’s direction (compass bearing) in relation to the observer. Prior to analysis, we used trigonometric relationships between distance and direction to calculate the bird’s location. We also excluded locations recorded when the bird was more than 100 m away from the observer, because low precision is expected at this distance. To calculate the proportion of time that radio-tagged hummingbirds spent in the focal area, we estimated the amount of spent at each location (see ‘*Estimating time at location’*, below), subset locations to those within the focal area outline (100% MCP) and calculated the total number of minutes spent in the focal area during each experimental period. We also calculated the estimated amount of time dedicated to radio telemetry observations for each experimental period (see ‘*Estimating observation effort’*, below).

***Estimating time at location***

For 1451/2451 of bird locations (59%), observers recorded the bird’s arrival time to that point, but not the bird’s departure time. These situations arose due to a trade-off between detailed data collection and maintaining contact with a rapidly moving animal. Therefore, we estimated ‘time at location’ as the intervening time before the next recorded location, unless the observer noted that the bird was moving and/or signal was poor; for these records, we assumed a time of 1 minute. The distribution of time at location estimated using adjacent time points aligned closely with the distribution of times recorded in the field (Fig. S2).

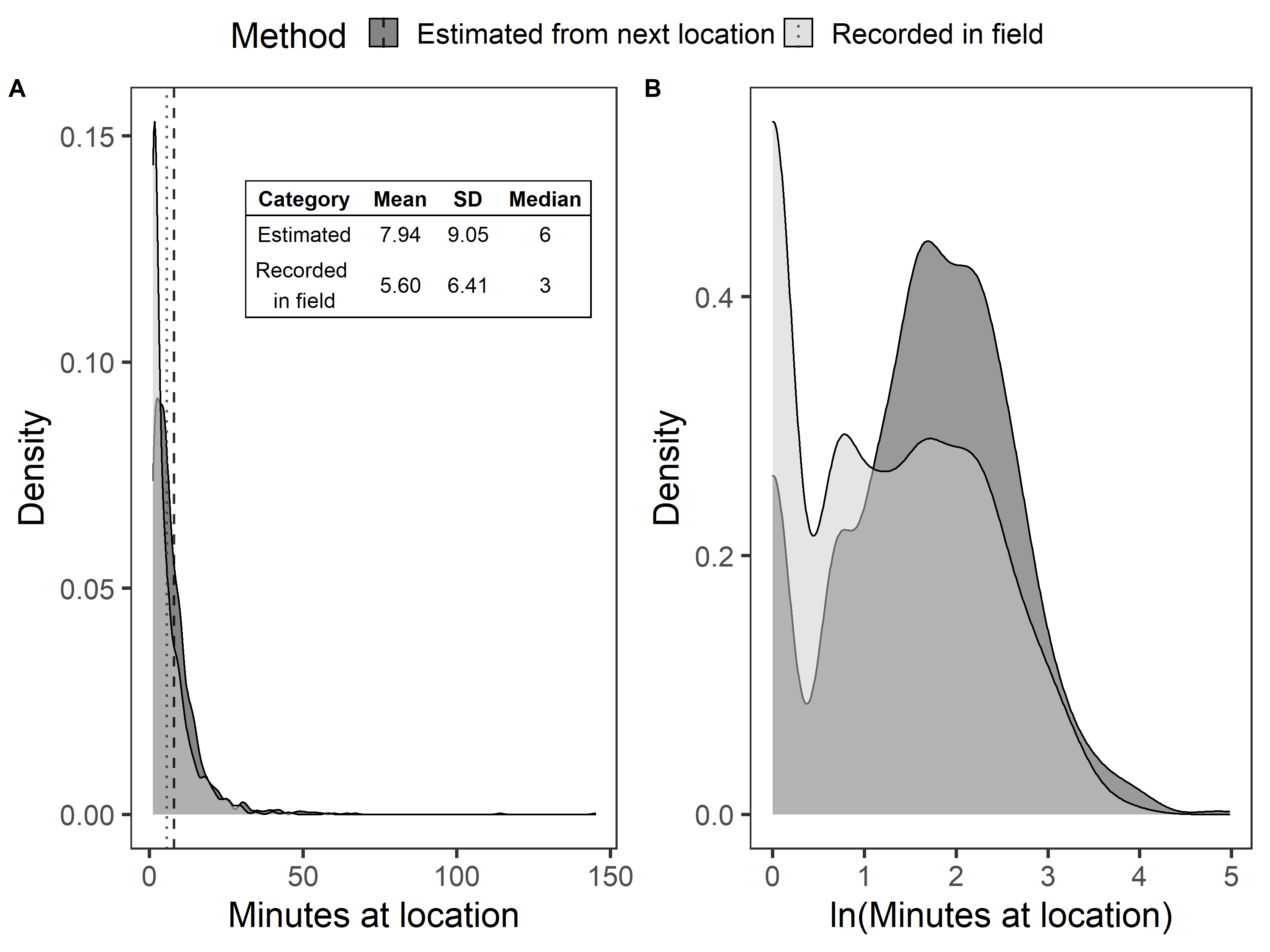

#### Figure S2. Density plots showing the amount of time radio-tagged hummingbirds spent at a given location, as calculated from arrival and departure times recorded in the field (light grey) *versus* the intervening time before the next recorded location (dark grey). Vertical lines indicate mean values.

***Estimating observation effort***

The length of each telemetry session (i.e., amount of daily effort spent following and/or searching for radio-tagged hummingbirds) was not always recorded by individual observers following individual birds. Multiple observers simultaneously followed different birds during each telemetry session per site, so we calculated daily site-level observation effort as the time between (1) the first bird location and/or recorded start time, and (2) the last bird location and/or recorded end time, across all observers following birds at the site. First and last observation times were reasonably close to recorded effort (Fig. S3; 3.06 ± 0.92 hours recorded effort *versus* 2.52 ± 0.96 hours estimated using first and last bird locations).

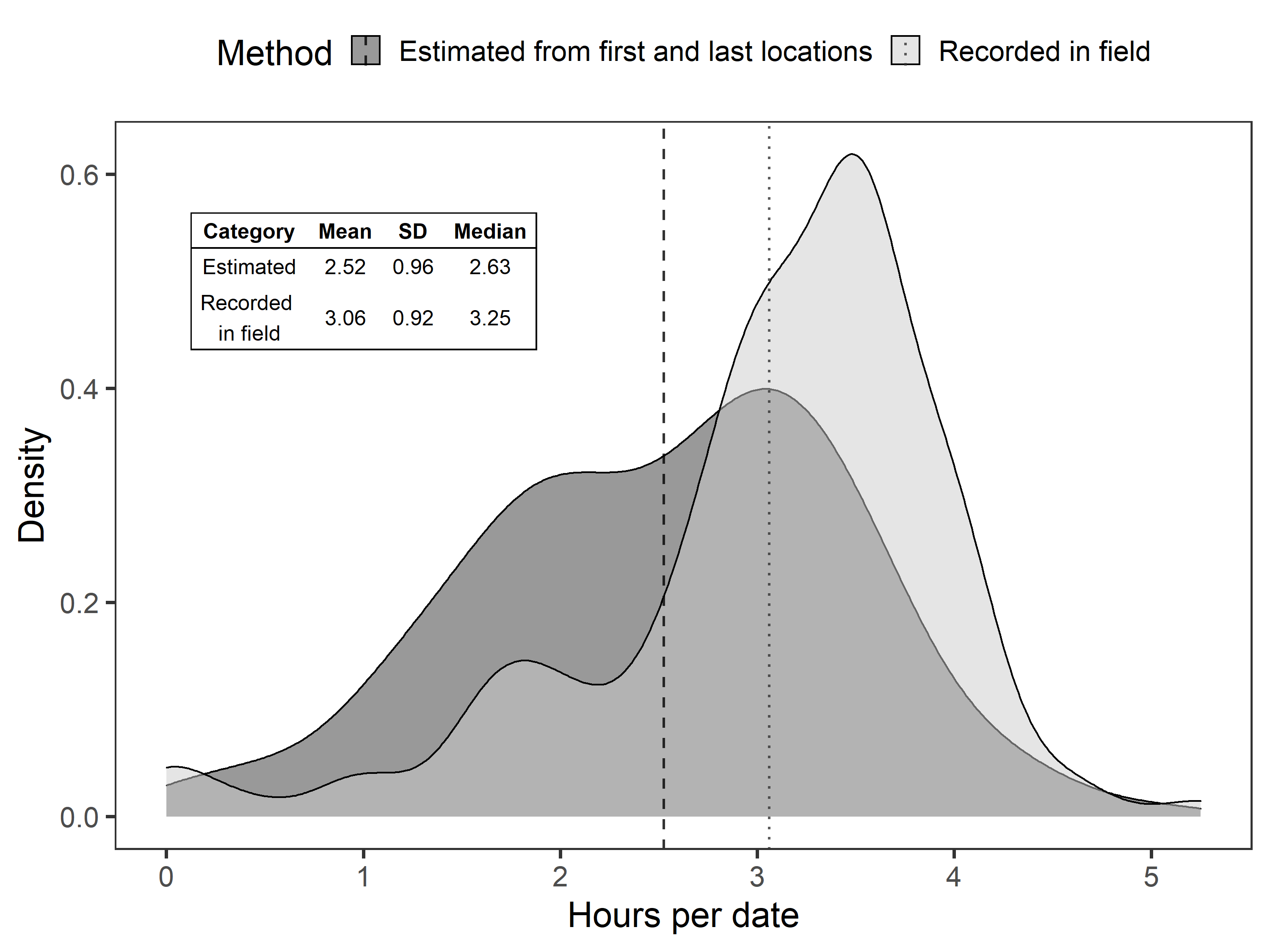

#### Figure S3. Density plots showing the amount of daily observation effort, as calculated from effort start and end times recorded in the field (light grey) *versus* the time between first and last bird locations, across all observers following birds at the site (dark grey). Vertical lines indicate mean values.

#### Table S5. Summary of radio-tracking outcomes for 72 birds that received radio transmitters. Species lacking obvious sexual dimorphism (i.e., rufous-tailed hummingbird and white-tipped sicklebill) were not identified to sex. Thirty-six birds that were present in focal areas during the ‘pre’ period were included in analysis of the *Heliconia* removal experiment.

|  | Green hermit | |  | Violet sabrewing | |  | Rufous-tailed hummingbird |  | Green-crowned brilliant |  | White-tipped sicklebill |  | Total |
| --- | --- | --- | --- | --- | --- | --- | --- | --- | --- | --- | --- | --- | --- |
|  | Male | Female |  | Male | Female |  | All |  | Female |  | All |  |  |
| ***Total birds tagged*** | 9 | 32 |  | 3 | 5 |  | 20 |  | 2 |  | 1 |  | **72** |
| ***Birds included in analysis*** *(Heliconia removal experiment)* | 3 | 21 |  | 2 | 1 |  | 8 |  | 1 |  | 0 |  | **36** |
| Attachment failure^1^ | 0 | 2 |  | 0 | 0 |  | 4 |  | 0 |  | 0 |  | 6 |
| Data reliability^2^ | 1 | 1 |  | 0 | 1 |  | 1 |  | 0 |  | 0 |  | 4 |
| Never detected^3^ | 2 | 1 |  | 0 | 1 |  | 1 |  | 0 |  | 0 |  | 5 |
| Never detected in focal area^4^ | 3 | 6 |  | 0 | 2 |  | 5 |  | 0 |  | 1 |  | 17 |
| Never detected focal area during ‘pre’ period | 0 | 1 |  | 1 | 0 |  | 1 |  | 1 |  | 0 |  | 4 |

1, Confirmed by locating detached transmitters.

2, Birds not detected in focal areas during telemetry observations but appeared on camera during the telemetry observation period.

3, Birds not detected during radio telemetry observations, either by observers or trail cameras.

4, Birds not detected in focal areas during telemetry observations, either by observers or trail cameras.

**Flower visitation**

***Camera installation***

Overall, >80% of cameras were located at a camera station, which comprised a focal *Heliconia* plant, a floral array of potted plants, and any naturally occurring flowering plants within a ~5 m radius of the focal *Heliconia* plant. After installing cameras at these plants, we searched for other flowering plants within each site for up to three person-hours. If any additional species were located, we distributed additional cameras, prioritizing species not already represented at the stations. On average, we obtained data from 10.3 ± 3.4 SD cameras per replicate (range: 2-17 cameras/site).

#### Table S6. List of plant species included in analysis of non-*Heliconia* flower visitation (*N* = 30). *Heliconia tortuosa* and three plant species that received no hummingbird visitation (*Calathea lutea*, *Ctenanthe dasycarpa*, *Maripa nicaraguensis*) are not included in this table. Plant species included in floral arrays are in bold (*N* = 13). Species analyzed for the presence of pollen tubes are denoted with an asterisk (*N* = 4), though only *Hamelia patens* yielded usable data.

| Family | Scientific name | Number of replicates in camera data |
| --- | --- | --- |
| MARANTACEAE | *Calathea crotalifera* | 24 |
| LAMIACEAE | ***Scutellaria costaricana**** | 22 |
| CAMPANULACEAE | ***Centropogon granulosus*** | 21 |
| GESNERIACEAE | ***Columnea polyantha*** | 18 |
| RUBIACEAE | ***Hamelia patens**** | 17 |
| ACANTHACEAE | ***Pachystachys lutea*** | 16 |
| VERBENACEAE | ***Stachytarpheta frantzii**** | 15 |
| GESNERIACEAE | ***Columnea raymondii*** | 14 |
| COSTACEAE | ***Costus barbatus*** | 11 |
| RUBIACEAE | *Palicourea padifolia* | 8 |
| MARANTACEAE | *Calathea guzmanioides* | 6 |
| ACANTHACEAE | ***Justicia oerstedii*** | 6 |
| ERICACEAE | *Cavendishia bracteata* | 4 |
| ZINGIBERACEAE | *Renealmia sp.* | 4 |
| ACANTHACEAE | ***Aphelandra golfodulcensis*** | 4 |
| MUSACEAE | *Musa x paradisiaca* | 3 |
| ZINGIBERACEAE | *Renealmia cernua* | 3 |
| MALVACEAE | ***Malvaviscus achanioides*** | 2 |
| CAMPANULACEAE | *Burmeistera cyclostigmata* | 2 |
| COSTACEAE | ***Costus woodsonii*** | 2 |
| GESNERIACEAE | *Drymonia sp.* | 2 |
| LAMIACEAE | ***Salvia splendens*** | 2 |
| ZINGIBERACEAE | *Elettaria cardamomum* | 1 |
| ERICACEAE | *Satyria warszewiczii* | 1 |
| COSTACEAE | *Costus laevis* | 1 |
| ZINGIBERACEAE | *Etlingera elatior* | 1 |
| RUBIACEAE | *Psychotria poeppigiana* | 1 |
| MYRTACEAE | *Syzygium malaccense* | 1 |
| BROMELIACEAE | *Pitcairnia imbricata* | 1 |
| FABACEAE | *Erythrina costaricensis* | 1 |

#### Table S7. Number of hours of video footage included in each camera dataset.

| Dataset | Purpose | Data excluded | Number of hours |
| --- | --- | --- | --- |
| Full dataset | Divide into subsets for purposes detailed below | Dates without any flowers visible on camera^1^ | 20,735 |
| Experimental | Examine hummingbird responses to experimental *Heliconia* removal | Cameras without videos for both experimental periods (pre and post) | 19,870 |
| Experimental (marked birds) | Examine how individual hummingbirds (marked with nail polish) responded to experimental *Heliconia* removal  Check the reliability of telemetry data | Cameras without videos for both experimental periods (pre and post)  2016 videos^2^ | 14,949 |
| Normal visitation | Understand ‘normal’ visitation patterns in this study system  Tailor estimates of resource availability to each hummingbird species group | Videos from ‘post’ period of treatment replicates | 15,119 |
| Normal visitation (*Heliconia*) | Determine which hummingbird species visited *Heliconia* most frequently | Videos from ‘post’ period of treatment replicates  Non-*Heliconia* species | 1,809 |

1, Some plant species did not produce open flowers every day

2, Birds were only marked with nail polish for the last two years of study (2017-2018)

***Floral arrays***

Floral arrays comprised ~3-5 potted plants of common species present in the study region (Table S6), either growing naturally in premontane forest or ornamentally in pollinator gardens. All plant species were visited by hummingbirds, though the exact species used varied depending on each year and replicate. We obtained plants by transplanting naturally occurring individuals, purchasing from nurseries, or growing plants from seed. Focal arrays were established in sites 0-8 days before the first capture session (mean ± SD: 3.7 ± 1.8 days, median: 4 days) to allow hummingbirds to discover the floral resources prior to data collection. Plants were watered every 1-3 days throughout the experimental period.

**Pollination success**

***Style collection***

At the beginning of the experiment, we marked any open flowers with a permanent marker, which signaled that pollination could have occurred before the plant was transported to the study site. We also used a permanent marker to mark flowers that were open on the cover day (or equivalent day in control replicates), thus distinguishing flowers open during ‘pre’ period from those open during the ‘post’ period. Marked flowers were not included in analysis, nor were flowers from the day after covering in treatment replicates.

***Pollen tube microscopy***

To visualize pollen tubes using epifluorescence microscopy, styles must be dyed with aniline blue, mounted on a microscope slide, crushed into a thin tissue layer. To prepare styles for examination, we first soaked them for 24-48 hours in distilled water. Then, depending on the style thickness of each plant species, we softened styles in an 8 M solution of NaOH for 0-4 days (0 days: *S. costaricana*, 1 day: *H. tortuosa*, *S. frantzii*, 4 days: *H. patens*), short enough to maintain some structural integrity but long enough to prevent the cover slip from breaking when the styles were crushed. After softening, styles were soaked in distilled water for 48 hours before being transferred to a 0.005% solution of aniline blue prepared with 0.05 M K_2_HPO_4_. After at least 6 hours in the aniline blue, we mounted the styles onto microscope slides with a drop of dye, applied a coverslip, and crushed the styles in a fluid motion with constant pressure. A single observer (K.G.L.) examined >80% of all styles, and all observers were naïve to the site and treatment.

***Species included in final analysis***

Pollen tubes were not observed in any *S. frantzii* styles (0/103). Similarly, pollen tubes were present very few *S. costaricana* styles (20/159) — and in no styles from the ‘pre’ period of control replicates—resulting in model non-convergence due to complete separation. The final analysis thus included two species, each analyzed separately: *Heliconia tortuosa* and *Hamelia patens*.

**Bird condition**

To analyze pre-to-post changes in hummingbird body mass, we used a measure of body mass accounting for intraspecific and interspecific variation in structural size (i.e., wing length). Accounting for structural size was necessary because we expected larger hummingbirds to have larger absolute changes in body mass, which could exert undue influence in our analyses. Using eight years of hummingbird banding data from this study system, we modeled the allometric relationship between body mass and wing length (i.e., Ln(mass) ~ Ln(wing length): Gill 1985, Brown and Bowers 1985, Rayner 1988) for eight hummingbird species (Fig. S4). For each recaptured individual, we divided its observed body mass by the body mass predicted for its wing length. Birds with relative body mass measurements >1 are heavier than expected for their structural size; similarly, birds with relative mass <1 are lighter than expected for their structural size.

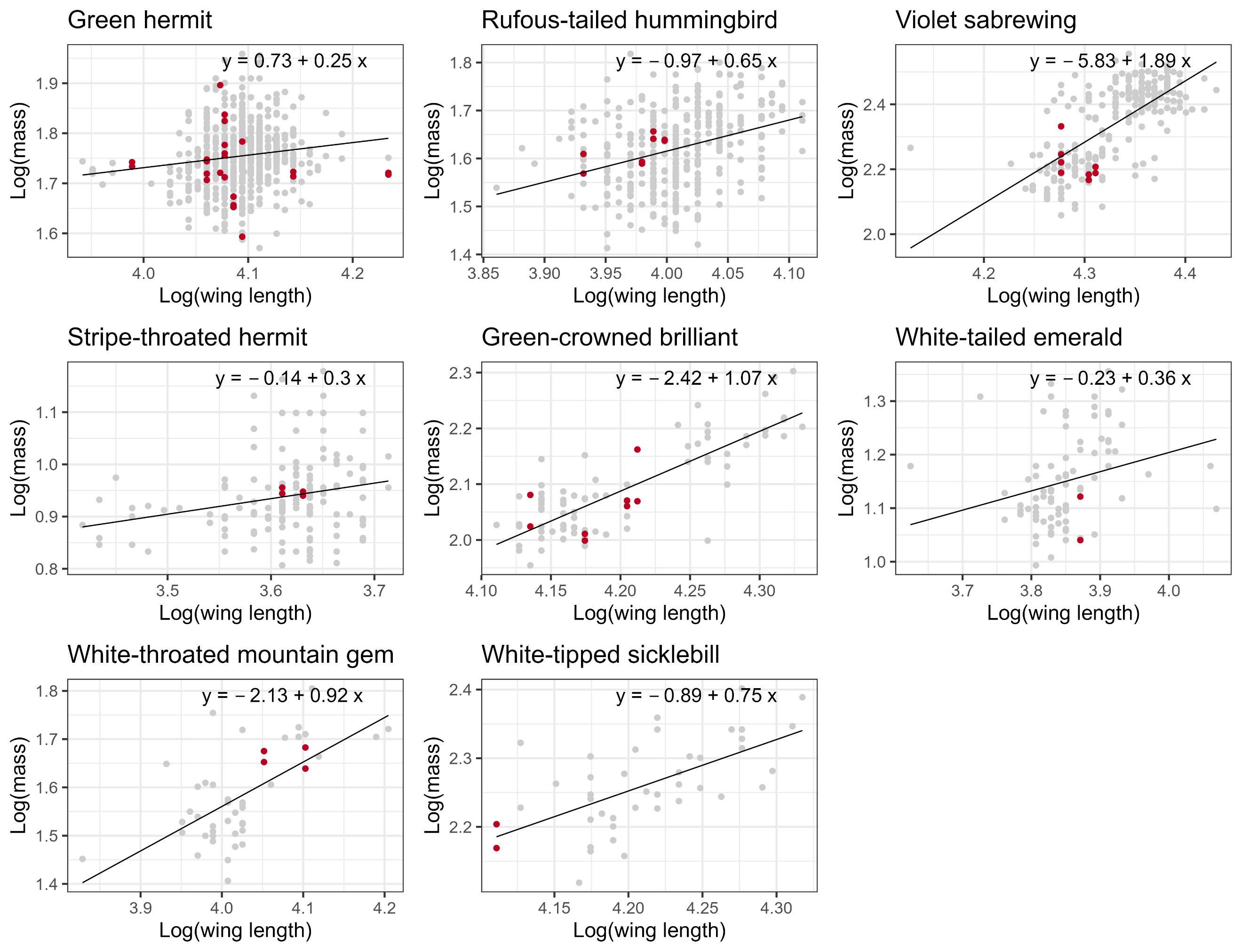

#### Figure S4. Allometric relationships between body mass and wing length (solid line), overlaid with morphological measurements from individual hummingbirds (points). Red points are measurements from the hummingbirds recaptured during both pre and post capture sessions of the *Heliconia* removal experiment (2016-2018). Light grey points are measurements from the individuals in the long-term dataset (2010-2018).

### Statistical analysis

#### Table S8. Summary of modeling approach used for the main response variables in the *Heliconia* removal experiment. ‘Replicate’ refers to each site-year combination and pairs the pre-to-post observations within each replicate. Zero-inflated models were specified as ziformula = ~1 (zero-inflated intercept). Random effects are nested; for example, individual bird ID (or plant species) within replicate and replicate within site.

|  | Response variable | Distribution and link function | Link function | Zero-inflated model | Method used to control for effort | Additional covariates^a^ | Random intercepts |
| --- | --- | --- | --- | --- | --- | --- | --- |
| Captures | Number of birds captured | Poisson | Log | No | Covariate:  Net-hours^b^ |  | Site, Replicate |
| Radio telemetry | Proportion of time spent in focal area | Betabinomial | Logit | No | Weights:  Number minutes of radio telemetry observation |  | Site, Replicate, Bird |
| Visitation rates: *Heliconia* | Number of visits to *Heliconia* | Negative binomial  (nbinom1) | Log | No | Offset:  Number of minutes of video observation | Mean number of open flowers per day | Site, Replicate |
| Visitation rates: non-*Heliconia* | Number of visits to non-*Heliconia* plant species | Negative binomial (nbinom1) | Log | No | Offset:  Number of minutes of video observation | Mean number of open flowers per day | Site, Replicate, Plant species |
| Pollen tubes: *Heliconia* | Proportion of styles with at least one pollen tube | Binomial | Logit | Yes | Weights:  Number of styles analyzed |  | Site, Replicate |
| Pollen tubes: *Hamelia* | Proportion of styles with at least one pollen tube | Binomial | Logit | No | Weights:  Number of styles analyzed |  | Site, Replicate |

a, All models included covariates related to the experiment (e.g., Control/Treatment, Pre/Post).

b, Net-hours as a covariate produced better model fit than net-hours as an offset.

#### Table S9. Summary of modeling approach used for the supplemental response variables in the *Heliconia* removal experiment. ‘Replicate’ refers to each site-year combination and pairs the pre-to-post observations within each replicate. Unless otherwise noted, random effects are nested; for example, individual bird ID (or plant species) within replicate and replicate within site.

|  | Response variable | Distribution and link function | Link function | Zero-inflated model | Method used to control for effort | Additional covariates^a^ | Random intercepts |
| --- | --- | --- | --- | --- | --- | --- | --- |
| Recaptures | Proportion of birds recaptured during ‘post’ period | Binomial | Logit | No | Weights:  Number of birds captured during ‘pre’ period  Did not explicitly control for capture effort^b^ |  | Site |
| Body mass | Relative body mass^c^ | Gaussian |  |  |  |  | Bird^d^ |
| Visitation rates: *Heliconia*  (marked birds) | Number of visits to *Heliconia* from birds marked with nail polish | Poisson | Log | No | Offset:  Number of minutes of video observation | Mean number of open flowers per day | Site, Replicate, Bird |
| Visitation rates: non-*Heliconia* (marked birds) | Number of visits to non-*Heliconia* plant species from birds marked with nail polish | Negative binomial  (nbinom2) | Log | No | Offset:  Number of minutes of video observation | Mean number of open flowers per day | Site, Replicate, Plant species  Bird^e^ |

a, All models included covariates related to the experiment (e.g., Control/Treatment, Pre/Post).

b, Capture effort was kept constant between pre and post sessions, except for 1 instance. This replicate (Site 203, 2017) was removed prior to analysis of recaptures.

c, Observed body mass relative to expected mass for its species and wing length.

d, Due to very limited sample size and problems with model convergence, we only included a random effect for individual Bird ID.

e, Bird ID was not nested within plant species because individual birds visited multiple plant species.

#### Table S10. Summary of modeling approach used to analyze visitation rates to individual non-*Heliconia* species. ‘Replicate’ refers to each site-year combination and pairs the pre-to-post observations within each replicate. In each model, the response variable was ‘number of visits’ and we controlled for effort using an offset (number of minutes of video observation) unless otherwise noted. Each model also included random intercepts for site and replicate. Note that the model for *S. frantzii* required slightly different specifications in terms of effort and covariates.

|  | Distribution and link function | Link function | Method used to control for effort | Additional covariates^a^ |
| --- | --- | --- | --- | --- |
| *Calathea crotalifera* | Negative binomial (nbinom1) | Log | Offset:  Number of minutes of video observation | Mean number of open flowers per day |
| *Scutellaria costaricana* | Negative binomial (nbinom1) | Log | Offset:  Number of minutes of video observation | Mean number of open flowers per day |
| *Centropogon granulosus* | Negative binomial (nbinom2) | Log | Offset:  Number of minutes of video observation | Mean number of open flowers per day |
| *Columnea polyantha* | Negative binomial (nbinom1) | Log | Offset:  Number of minutes of video observation | Mean number of open flowers per day |
| *Hamelia patens* | Negative binomial (nbinom2) | Log | Offset:  Number of minutes of video observation | Mean number of open flowers per day |
| *Pachystachys lutea* | Negative binomial (nbinom1) | Log | Offset:  Number of minutes of video observation | Mean number of open flowers per day |
| *Stachytarpheta frantzii* | Negative binomial (nbinom1) | Log | Covariate:  Number of minutes of video observation^b^ | Mean number of open flowers per day  (quadratic term) |
| *Columnea raymondii* | Poisson | Log | Offset:  Number of minutes of video observation | Mean number of open flowers per day |
| *Costus barbatus* | Poisson | Log | Offset:  Number of minutes of video observation | Mean number of open flowers per day |

a, All models included covariates related to the experiment (e.g., Control/Treatment, Pre/Post).

b, Hours as a covariate produced better model fit than hours as an offset.

### Literature cited

Brown, J. H., and M. A. Bowers. 1985. Community organization in hummingbirds: relationships between morphology and ecology. The Auk 102:251–269.

Gill, F. B. 1985. Hummingbird flight speeds. The Auk 102:97–101.

Gorelick, N., M. Hancher, M. Dixon, S. Ilyushchenko, D. Thau, and R. Moore. 2017. Google Earth Engine: Planetary-scale geospatial analysis for everyone. Remote Sensing of Environment 202:18–27.

Hadley, A. S., and M. G. Betts. 2009. Tropical deforestation alters hummingbird movement patterns. Biology Letters 5:207–210.

Rayner, J. M. V. 1988. Form and function in avian flight. Pages 1–66 *in* R. F. Johnston, editor. Current Ornithology. Springer US, Boston, MA.

Volpe, N. L., A. S. Hadley, W. D. Robinson, and M. G. Betts. 2014. Functional connectivity experiments reflect routine movement behavior of a tropical hummingbird species. Ecological Applications 24:2122–2131.

Volpe, N. L., W. D. Robinson, S. J. K. Frey, A. S. Hadley, and M. G. Betts. 2016. Tropical forest fragmentation limits movements, but not occurrence of a generalist pollinator species. PLOS ONE 11:e0167513.
