## Appendix S3 for "Tropical hummingbird pollination networks are resistant to short-term experimental removal of a common flowering plant"

**Appendix S3:** Supplemental Methods (Quantifying resource availability)

Supporting Information for:

**Tropical hummingbird pollination networks are resistant to short-term experimental removal of a common flowering plant**

Kara G. Leimberger^1^, Adam S. Hadley^1,2^, Sarah J.K. Frey^1,3^, and Matthew G. Betts^1^

^1^Forest Biodiversity Research Network, Department of Forest Ecosystems and Society, Oregon State University, Corvallis, Oregon, USA

^2^Biodiversity Section, Department of Natural Resources and Energy Development, Fredericton, New Brunswick, Canada

^3^Department of Animal and Rangeland Sciences, Oregon State University, Corvallis, Oregon, USA

### Quantifying resource availability

***Resource surveys***

Across all sites, we recorded 75 known and 15 unknown species of flowering plants known or suspected to be visited by hummingbirds (Table S1). We did not exclude any species based solely on pollination syndrome, because hummingbirds can be highly opportunistic foragers (Maruyama *et al.* 2013). Six of the unknown species could be identified to genus and were retained for calorie estimation purposes; the remaining species were excluded from further analysis (*N* = 28 plants), as were four species of Melastomataceae and Myrsinaceae known or suspected to be nectarless (*N* = 9 plants). We also excluded two species known or suspected to be pollinated by bats (*Burmeistera cyclostigmata* and *Mucuna globulifera*). Although hummingbirds occasionally visit night-blooming flowers at dusk or in the early morning (Muchhala 2003), we did not attempt to estimate the amount of nectar that would be present at those times, because nectar availability likely depends on bat visitation rate.

***Nectar measurements***

To estimate the number of calories provided by each plant species, we measured nectar properties for 39 species growing in study sites and/or the Wilson Botanical Garden at the Las Cruces Biological Station. In the afternoon or evening prior to nectar sampling, we covered flowers with mesh bags to prevent pollinators from removing nectar. On the following morning or early afternoon, we collected the covered flowers and transported them to the field station for processing (mean ± SD collection time: 11:00 ± 2.5 hrs). We extracted nectar and measured its volume using microcapillary tubes, then used a handheld refractometer to measure nectar concentration as degrees Brix (i.e., grams of sucrose per 100 grams of solution). A total of 1438 flowers were sampled, 220 flowers in 2014 (February - March) and 1218 flowers in 2018 (February - May). We quantified nectar volume by cutting the nectary open with a razor blade, extracting the nectar with a microcapillary tube, and measuring the height of the nectar in the tube. Nectar height (mm) was then converted to volume (µL) using the conversion factors in Table S2. We measured sugar concentration as degrees Brix (i.e., grams of sucrose per 100 grams of solution) using a handheld refractometer with a measurement range of 0-32 ºBx. Some species produced flowers with nectar concentrations above the maximum limit of the refractometer, such as *Passiflora coccinea* and *Calathea crotalifera*. Flowers above the maximum range of the refractometer were assigned nectar concentrations of 33 ºBx and accounted for approx. 3% of all flowers sampled. We did not correct nectar concentrations for ambient temperature, because 80% of the measurements were collected using a refractometer equipped with automatic temperature compensation (Fisher Scientific 0-32 ºBx with ATC), and temperature correction for the remaining samples would not have changed the concentration more than 1 ºBx.

***Calculating calories per flower***

To assign caloric values per flower, we first calculated average nectar volume and concentration. For the species without nectar measurements, we searched the literature and prioritized data from study sites in Costa Rica when data were available from multiple sources. If no data were unavailable, we substituted measurements from a congeneric species, or—if no congeneric measurements were available—another closely related species within the family. We converted nectar volumes and concentrations to calories per flower following the methods of Bolten *et al.* (1979) and Kearns & Inouye (1993). First, we converted nectar concentrations from degrees Brix (% sucrose, g/g) to g/L using Kearns & Inouye’s (1993) Table 5-2 (included here as Table S3). We then multiplied nectar concentration (g/L) by nectar volume (L) to calculate grams of sucrose per flower. Lastly, this value converted to kilocalories by multiplying by 3.94 (Cal/g). Per-flower nectar measurements and caloric values are presented in Table S4.

***Calculating calories per plant***

To calculate calories per plant, we multiplied calories per flower by flowers per plant. However, the number of flowers was not always recorded resource surveys; rather, larger units, such as the number of inflorescences, flower-producing bracts, or trees were used instead. We converted these observations to ‘number of flowers’ as follows. When surveying plants in the genus *Heliconia*, we recorded the number of bracts that had potential to produce flowers. In practice, this meant only counting open bracts with accumulated flower material; *Heliconia* bracts gradually unfurl as the flowering season begins, and on any given day, a subset of bracts produces single-day flowers that accumulate in the bract after abscission. In 2018, we also recorded the number of freshly opened *Heliconia tortuosa* flowers, and from these data we calculated — for plants with 1-14 open bracts — the median number of open flowers on a given day (Table S5). We used these values to convert number of open bracts to number of flowers for all *Heliconia* species.

To determine flowers per inflorescence for non-*Heliconia* species, we supplemented unpublished data with opportunistic observations gathered during resource counts (2016-2018), nectar sampling (2018), and from focal plants monitored for hummingbird visitation (2017-2018). Despite these efforts, certain uncommon species were not included in our measurements; for these 34 species, an experienced field technician (M. Atencio) provided estimates of flowers per inflorescence based on personal observation (Table S6). Estimations based on personal observation were only applied to approx. 2% of the total plants recorded. At the beginning of the study, resources were also occasionally counted using ‘tree’ as the counting unit. We estimated the number of flowers per tree by combining personal observation with data from the literature. Tree-to-flower conversions (Table S7) were applied to approx. 0.5% of the total plants recorded. Finally, counting units were not available for approx. 10% of plants; for these plants, we assumed observers counted inflorescences, leading to a maximum estimate of non-*Heliconia* resource availability and minimum (conservative) estimate of percentage calories removed. We used the conservative estimate when calculating the percentage of calories removed by our experimental manipulation.
**Supplemental tables associated with this section are provided in a separate Excel file.**

### Table S1. List of 75 plant species recorded in 14 forest fragments surrounding the Las Cruces Biological Research Station. Nectarless or night-blooming species were omitted from calorie estimation (see 'Notes' column).

### Table S2. Conversion factors used to calculate nectar volume (µL) from nectar height (mm) in microcapillary tubes of different sizes (5-100 µL).

### Table S3. Table 5-2 from Kearns & Inouye (1993), used to convert ºBrix (g/g) to g/L for calorie estimation. Before applying these conversions, nectar concentrations less than 10 ºBrix were rounded to the nearest 0.5, concentrations of 10-20 ºBrix were rounded to the nearest 1.0, and concentrations greater than 20 ºBrix were rounded to the nearest 2.0.

### Table S4. Per-flower nectar volume, concentration, and caloric content for the plant species recorded in this study. Nectar measurements were collected from 1438 flowers representing 38 species growing in the Wilson Botanical Garden at the Las Cruces Biological Station and forest fragments in the surrounding landscape. To prevent pollinator access, flowers were bagged the day prior to nectar collection. For species without nectar measurements, we searched the literature for data on nectar volume and concentration. If data were available from multiple sources, we prioritized data from study sites in Costa Rica. If data were unavailable, we substituted measurements from a congeneric species, or—if no congeneric measurements were available—another closely related species within the family.

### Table S5. Number of open flowers per inflorescence for 2219 *Heliconia tortuosa* inflorescences with varying numbers of open bracts. Plants were sampled in 14 forest fragments surrounding the Las Cruces Biological Research Station between February and April 2018. The median number of flowers per inflorescence was used in calorie estimation except for plants with uncommonly high bract numbers; these plants (>9 bracts) were assigned two flowers per inflorescence.

### Table S6. Data used to convert number of non-*Heliconia* inflorescences to number of flowers, for calorie estimation. Data were compiled from a variety of sources: inflorescences sampled at the Las Cruces Biological Garden by C. Birkett between February and March 2014 and opportunistic observations from forest fragments in the surrounding landscape gathered during resource surveys, daily nectar sampling, and from focal plants monitored for hummingbird visitation. For the 31 species with missing data, an experienced field technician provided rough estimations based on personal observation; these estimations were applied to 117 plants, or approx. 2% of the total plants counted.

### Table S7. Data used to convert number of trees to number of flowers, for calorie estimation. Estimates were informed by personal observation and literature research. Because smaller counting units (e.g., inflorescences or flowers) were typically used during resource surveys, these flowers-per-tree estimates were only applied to 34 plants (approx. 0.5% of the total plants counted).

### Table S8. Weights used to create metrics of caloric availability specific to each hummingbird group (e.g., all species detected on camera *versus* green hermits and violet sabrewings *versus* all species detected visiting *Heliconia*). Weights are relative visitation rates, ascertained through 15,119 hours of video observation. Video observations are from control replicates and the ‘pre’ period of treatment replicates, only include videos with at least one flower on camera, and include sightings with confirmed visits of any type (i.e., honest or rob). *Burmeistera cyclostigmata* and *Mucuna globulifera* were not included in calorie estimates due to challenges estimated nectar availability for plant species with night-blooming flowers. Similarly, *Heliconia* was omitted from estimates of non-*Heliconia* resources.

### Literature cited

Bolten, A. B., P. Feinsinger, H. G. Baker, and I. Baker. 1979. On the calculation of sugar concentration in flower nectar. Oecologia 41:301–304.

Kearns, C. A., and D. W. Inouye. 1993. Techniques for Pollination Biologists. University Press of Colorado, Niwot.

Maruyama, P. K., G. M. Oliveira, C. Ferreira, B. Dalsgaard, and P. E. Oliveira. 2013. Pollination syndromes ignored: importance of non-ornithophilous flowers to Neotropical savanna hummingbirds. Naturwissenschaften 100:1061–1068.

Muchhala, N. 2003. Exploring the boundary between pollination syndromes: bats and hummingbirds as pollinators of *Burmeistera cyclostigmata* and *B. tenuiflora* (Campanulaceae). Oecologia 134:373–380.
