## Appendix S4 for "Tropical hummingbird pollination networks are resistant to short-term experimental removal of a common flowering plant"

**Appendix S4:** Supplemental Results

Supporting Information for:

**Tropical hummingbird pollination networks are resistant to short-term experimental removal of a common flowering plant**

Kara G. Leimberger^1^, Adam S. Hadley^1,2^, Sarah J.K. Frey^1,3^, and Matthew G. Betts^1^

^1^Forest Biodiversity Research Network, Department of Forest Ecosystems and Society, Oregon State University, Corvallis, Oregon, USA

^2^Biodiversity Section, Department of Natural Resources and Energy Development, Fredericton, New Brunswick, Canada

^3^Department of Animal and Rangeland Sciences, Oregon State University, Corvallis, Oregon, USA

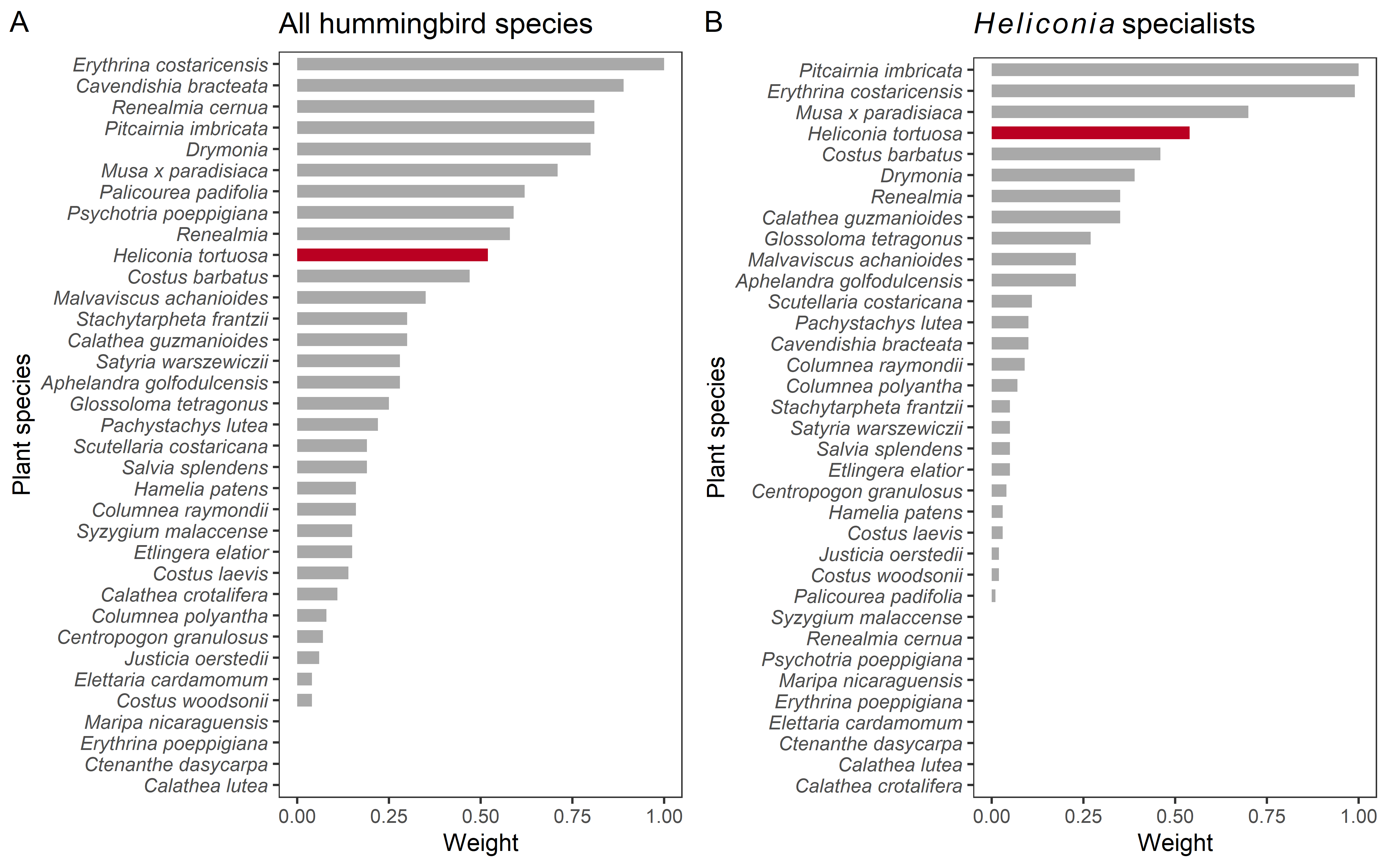

### **Figure S1.** Relative visitation rates (weights) used to tailor our resource availability estimates to different groups of hummingbird species. Within each hummingbird group, the most frequently visited plant species was assigned a weight of one. Visitation rates are summarized from the 15,119 hours of ‘normal’ (i.e., unmanipulated) camera observations collected in the landscape surrounding the Las Cruces Biological Station across 14 study sites, three field seasons (2016–2018), and 35 plant species. (A) Weights summarized across all hummingbird species detected on camera (*N* = 17 species). (B) Weights for green hermits (*Phaethornis guy*) and violet sabrewings (*Campylopterus hemileucurus*), two hummingbird species with morphologically specialized bill shapes that mirror the curvature of Heliconia flowers. These two species are also the most frequent *Heliconia* visitors.

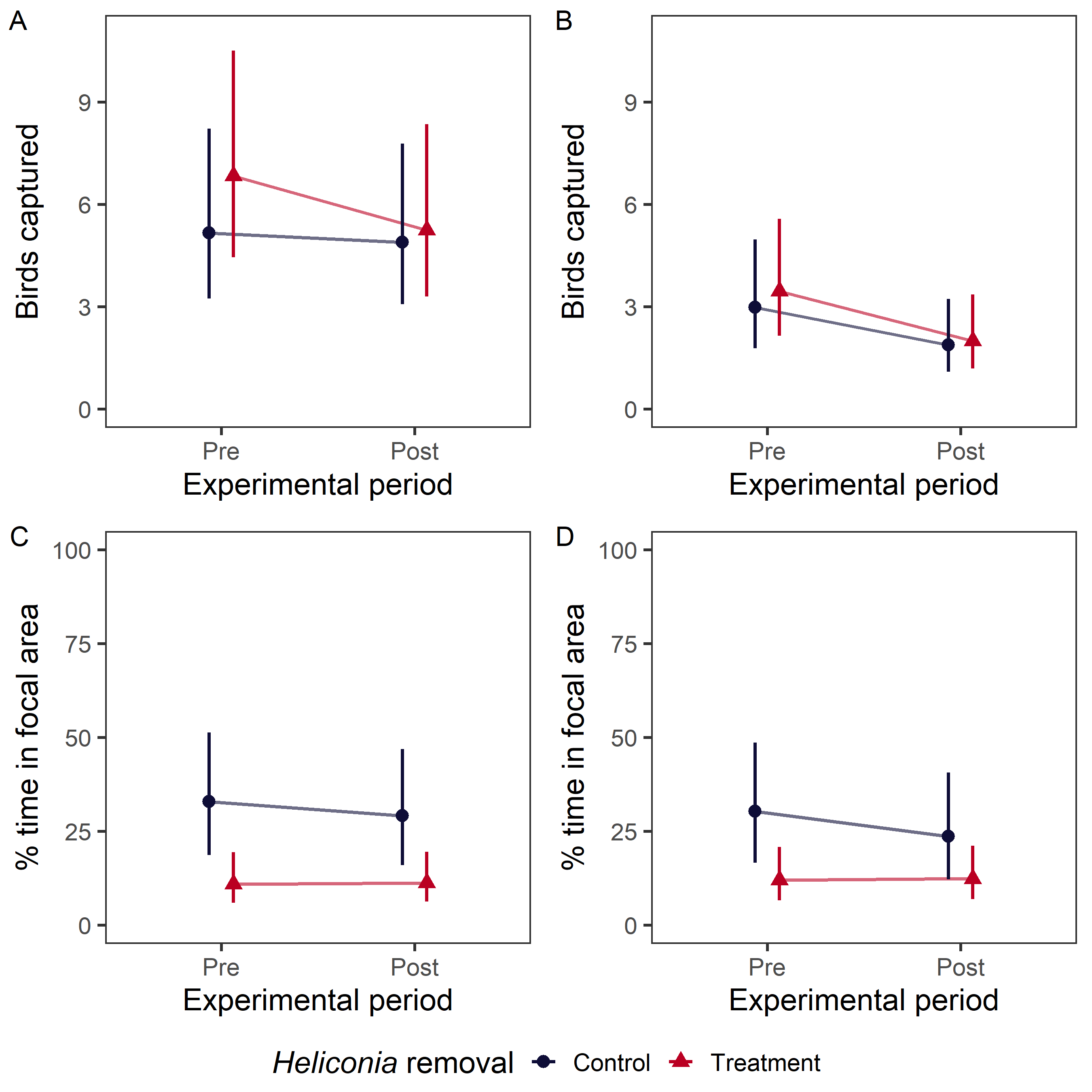

### **Figure S2.** Effects of experimental *Heliconia* removal on hummingbird abundance and space use. Estimated marginal means from GLMMs are presented alongside 95% confidence intervals; conceptually, a treatment effect is indicated by non-parallel lines. (A-B) Estimated number of captured hummingbirds, calculated for the mean capture effort across all replicates. (C-D) Estimated percentage of time that radio-tagged hummingbirds spent in the focal area. Left column = all hummingbird species, right column = green hermits and violet sabrewings only.

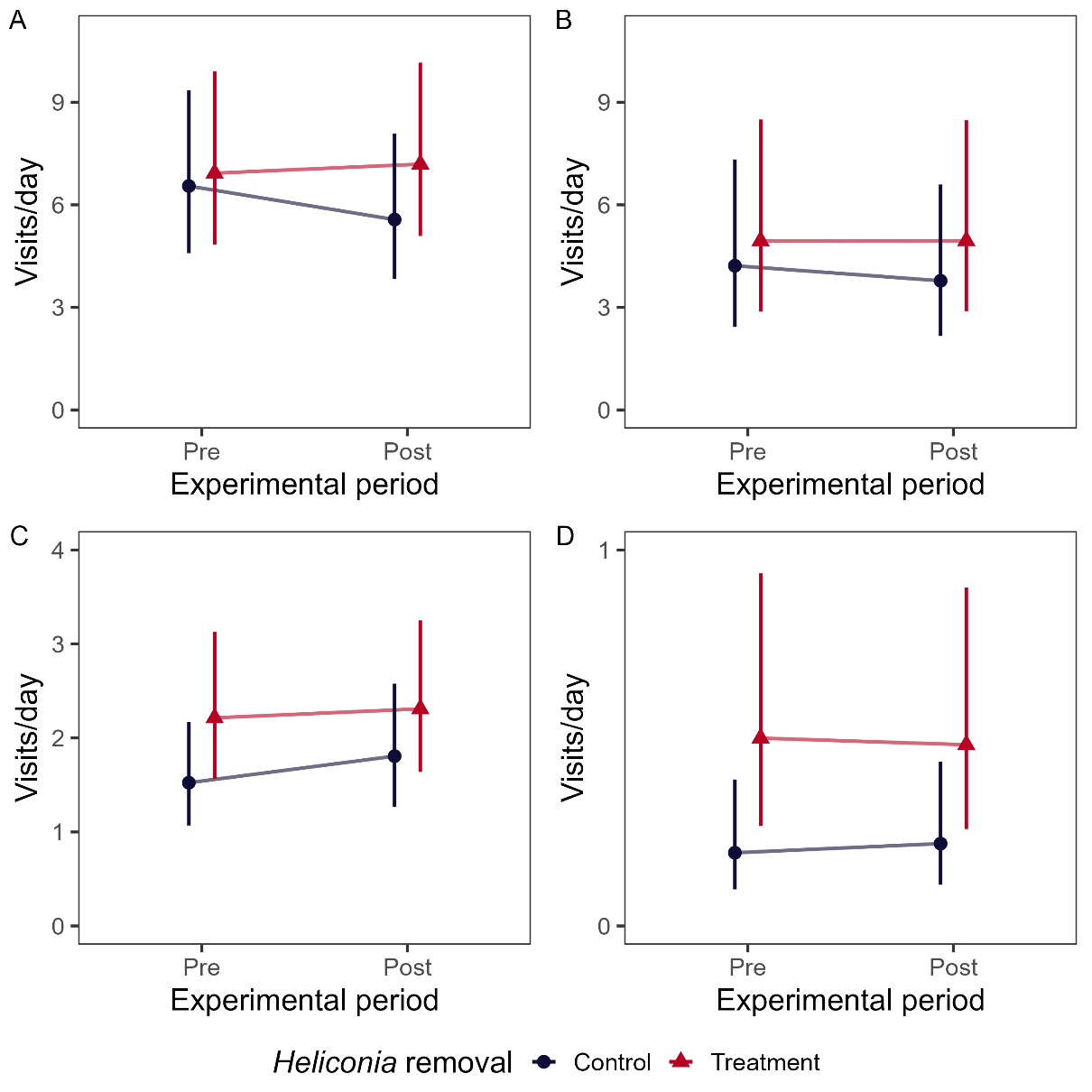

### **Figure S3.** Effects of experimental *Heliconia* removal on hummingbird flower visitation to focal *Heliconia* plants (A-B) and 30 non-*Heliconia* plant species (C-D). Estimated marginal means from GLMMs are presented alongside 95% confidence intervals; conceptually, a treatment effect is indicated by non-parallel lines. *Y*-axis is the estimated number of hummingbird visits, calculated for a 12-hr day. Left column = all hummingbird species, right column = green hermits and violet sabrewings only.

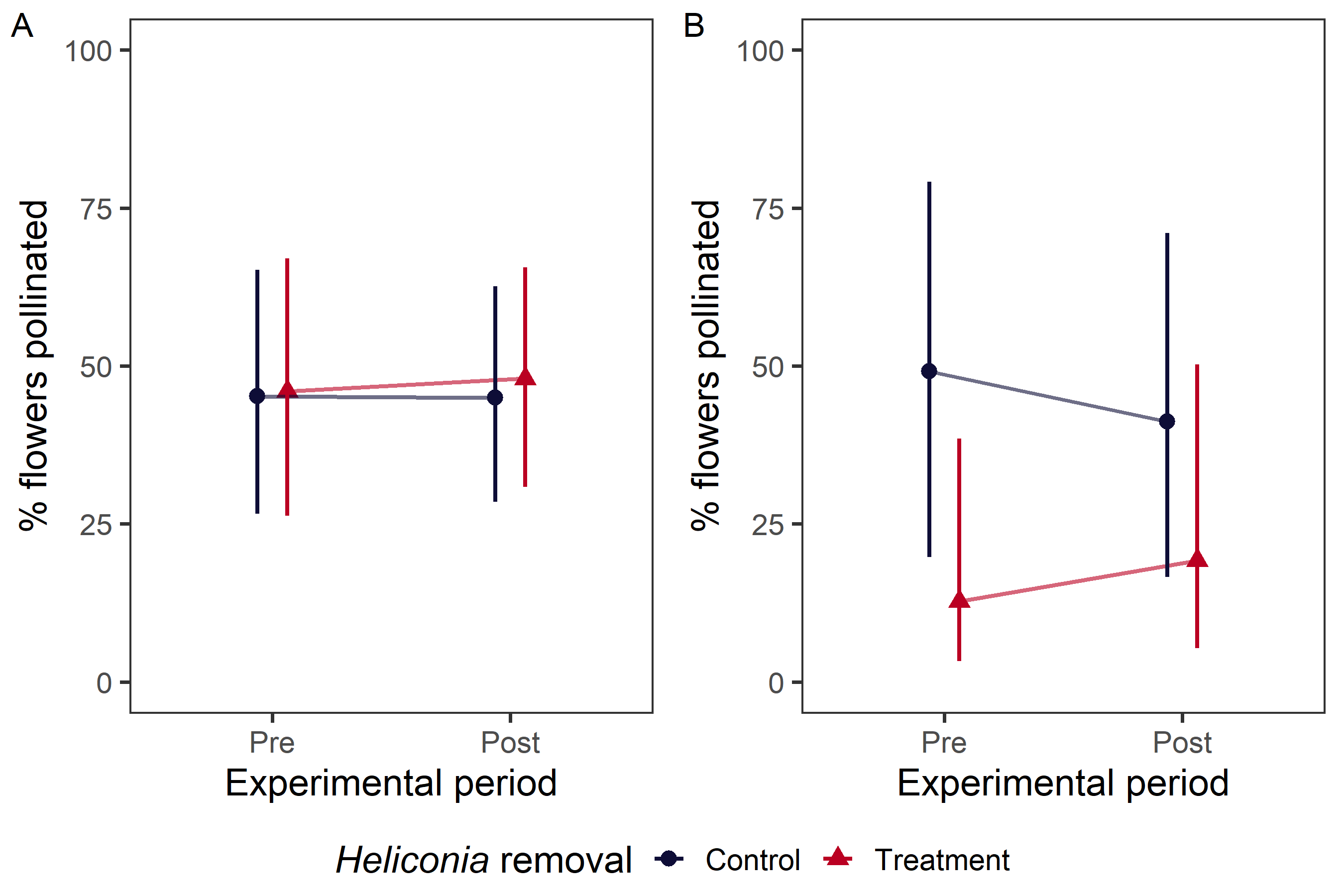

### **Figure S4.** Effects of experimental *Heliconia* removal on plant pollination success. Estimated marginal means from GLMMs are presented alongside 95% confidence intervals; conceptually, a treatment effect is indicated by non-parallel lines. (A) Percentage of flowers pollinated for focal *Heliconia* plants that remained uncovered throughout the experimental period (2 plants/site). (B) Percentage of flowers pollinated for *Hamelia patens*, a potted plant in the floral array (2 floral arrays/site).

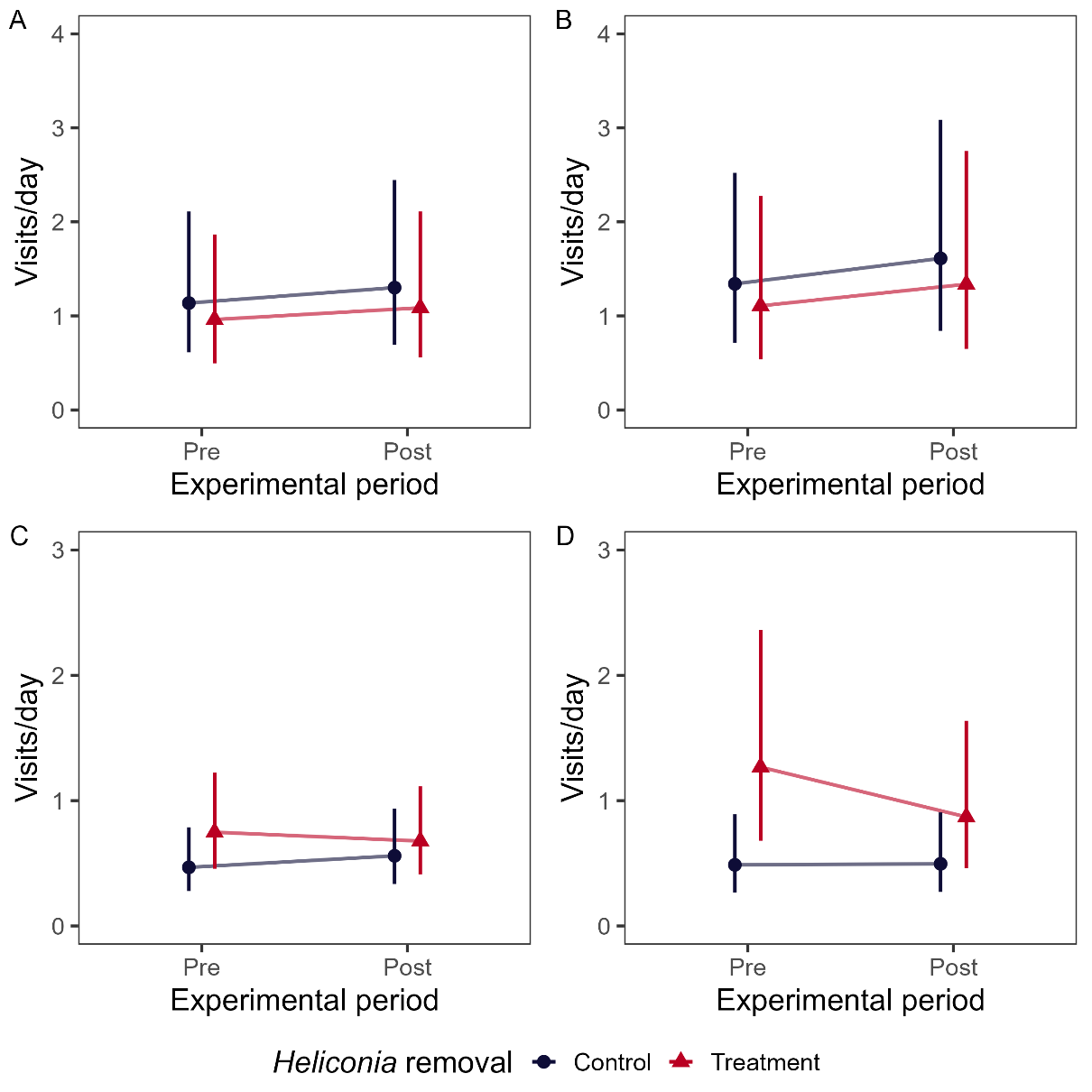

### **Figure S5.** Effects of experimental *Heliconia* removal on hummingbird flower visitation to focal *Heliconia* plants (A-B) and 17 non-*Heliconia* plant species (C-D) by individual hummingbirds marked with nail polish (2017-2018 only). Estimated marginal means from GLMMs are presented alongside 95% confidence intervals; conceptually, a treatment effect is indicated by non-parallel lines. *Y*-axis is the estimated number of hummingbird visits, calculated for a 12-hr day. Left column = all hummingbird species, right column = green hermits and violet sabrewings only.

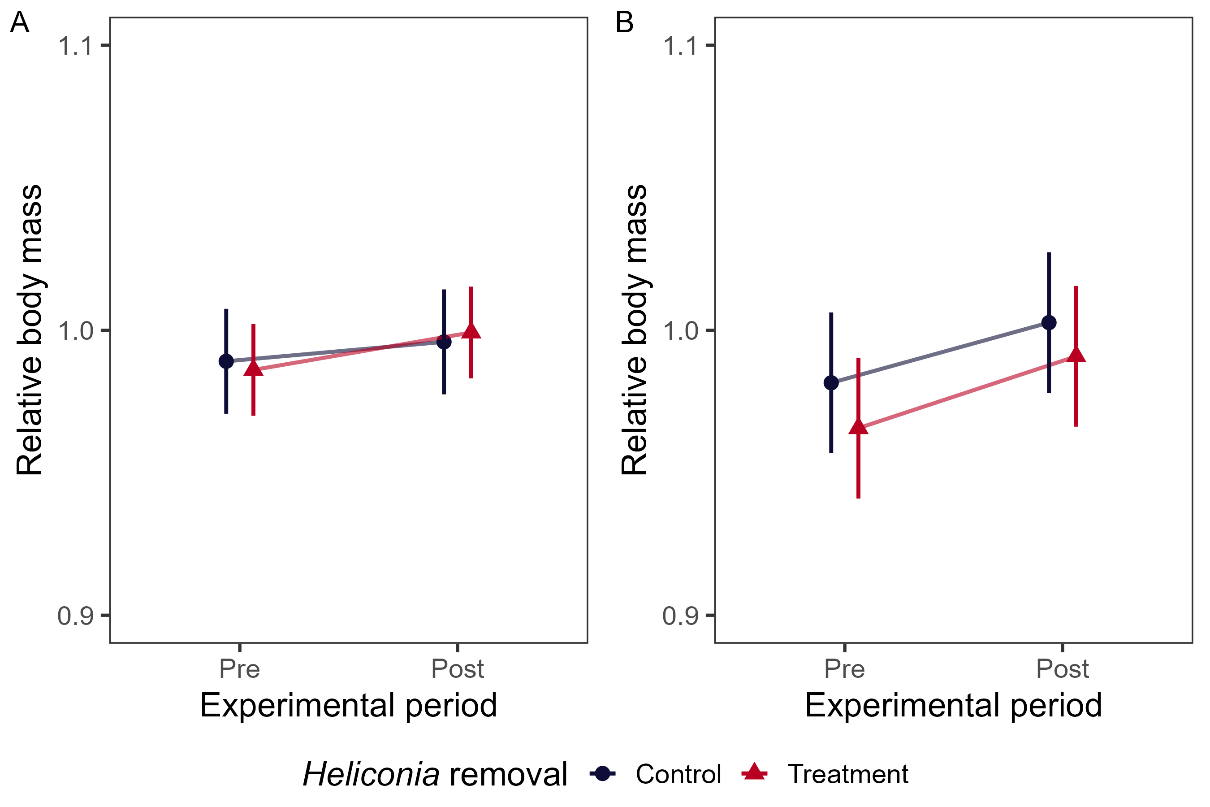

### **Figure S6.** Effects of experimental *Heliconia* removal on the body mass of recaptured hummingbirds, calculated relative to predicted mass based on the bird’s structural size (wing length). Estimated marginal means from GLMMs are presented alongside 95% confidence intervals; conceptually, a treatment effect is indicated by non-parallel lines. (A) All hummingbird species; (B) green hermits and violet sabrewings only.

### **Table S1.** Hummingbird species (*N* = 19) detected in fourteen study sites surrounding the Las Cruces Biological Station, 2016-2018. Cells marked with ‘x’ indicate at least one individual was captured during this study period. *Heliothrix barroti* and *Phaethornis longirostris* were only detected during video observations and are denoted with an asterisk. One captured species, *Glaucis aeneus*, was not detected on camera. Out of 383 total hummingbird captures, 332 were unique individuals. Values are sorted by the number of individuals captured without experimental manipulation, i.e., in control sites or in treatment sites prior to *Heliconia* removal.

| Common name | Scientific name | *Site 10* | *Site 130* | *Site 137* | *Site 200* | *Site 201* | *Site 203* | *Site 204* | *Site 205* | *Site 24* | *Site 29* | *Site 30* | *Site 49* | *Site 58* | *Site 60* | *Number of captures* | *Number of individuals* | *Number of individuals:*  *no manipulation* |
| --- | --- | --- | --- | --- | --- | --- | --- | --- | --- | --- | --- | --- | --- | --- | --- | --- | --- | --- |
| Green hermit | *Phaethornis guy* | x | x | x | x | x | x | x | x | x | x | x | x | x | x | 110 | 86 | 71 |
| Rufous-tailed hummingbird | *Amazilia tzacatl* | x | x | x | x | x | x | x |  | x |  | x | x | x | x | 95 | 89 | 60 |
| Violet sabrewing | *Campylopterus hemileucurus* | x | x |  | x | x | x |  | x | x | x | x |  | x | x | 50 | 45 | 36 |
| Stripe-throated hermit | *Phaethornis striigularis* | x | x |  |  | x | x |  | x | x | x | x | x |  | x | 43 | 37 | 23 |
| White-tailed emerald | *Elvira chionura* |  |  |  | x | x | x |  | x | x | x | x |  | x |  | 19 | 17 | 16 |
| Green-crowned brilliant | *Heliodoxa jacula* |  |  |  | x |  | x |  |  | x | x | x |  |  |  | 22 | 18 | 14 |
| White-tipped sicklebill | *Eutoxeres aquila* |  |  |  |  | x |  |  | x | x |  | x |  |  | x | 11 | 10 | 8 |
| White-throated mountain gem | *Lampornis castaneoventris* |  |  |  |  |  | x |  |  | x |  |  |  |  |  | 11 | 8 | 6 |
| Charming hummingbird | *Amazilia decora* | x |  |  | x | x |  |  |  |  |  |  |  |  |  | 4 | 4 | 3 |
| Garden emerald | *Chlorostilbon assimilis* | x | x |  |  |  |  |  |  | x |  |  |  |  |  | 4 | 4 | 3 |
| Snowy-bellied hummingbird | *Amazilia edward* | x |  | x |  | x |  |  |  |  |  |  |  |  |  | 3 | 3 | 3 |
| Scaly-breasted hummingbird | *Phaeochroa cuvierii* |  |  |  |  | x |  |  |  |  |  |  | x |  |  | 3 | 3 | 2 |
| Long-billed starthroat | *Heliomaster longirostris* | x |  |  |  |  | x |  |  |  |  |  |  |  |  | 2 | 2 | 2 |
| Violet-crowned woodnymph | *Thalurania colombica* |  |  |  |  | x |  |  |  |  |  |  |  |  |  | 2 | 2 | 2 |
| Green-fronted lancebill | *Doryfera ludovicae* |  |  |  |  |  |  |  |  |  | x |  |  |  |  | 2 | 2 | 2 |
| Bronzy hermit | *Glaucis aeneus* |  |  |  |  |  |  |  |  |  |  |  | x |  |  | 1 | 1 | 1 |
| Blue-throated goldentail | *Hylocharis eliciae* | x |  |  |  |  |  |  |  |  |  |  |  |  |  | 1 | 1 | 0 |
| Purple-crowned fairy | *Heliothryx barroti* |  |  |  |  |  |  |  |  |  |  | * |  |  |  | NA | NA | NA |
| Long-billed hermit | *Phaethornis longirostris* |  |  |  | * |  |  |  |  |  |  |  |  |  |  | NA | NA | NA |

### **Table S2.** Estimated percentage of calories removed in treatment sites, calculated as *Heliconia* calories removed divided by total calories available to hummingbirds. Leveraging information about hummingbird visitation patterns in the camera data, we tailored estimates of ‘availability’ to the hummingbird perspective. Values are ordered by the ‘no tailoring’ column, i.e., the lowest overall percentage removed.

|  |  | No tailoring | Subset to plant species visited by each group^1^ | | Subset and weighted by relative visitation rate^2^ | |
| --- | --- | --- | --- | --- | --- | --- |
| Year | Site | All species | All species | *Heliconia* specialists | All species | *Heliconia* specialists |
| 2017 | 201 | 0.2 | 0.9 | 0.9 | 2.2 | 7.9 |
| 2016 | 10 | 0.3 | 0.4 | 16.2 | 1.3 | 9.8 |
| 2017 | 60 | 0.3 | 9.6 | 15.4 | 37.4 | 65.4 |
| 2017 | 203 | 0.4 | 0.4 | 0.4 | 0.4 | 1 |
| 2018 | 60 | 1 | 34.6 | 71.9 | 60.7 | 79.7 |
| 2017 | 24 | 1.6 | 10.2 | 14.7 | 33.2 | 66.9 |
| 2018 | 10 | 2.2 | 13.9 | 17.2 | 14.3 | 78.7 |
| 2017 | 49 | 5.8 | 53.8 | 69.5 | 86.6 | 96.8 |
| 2016 | 137 | 6.3 | 57.4 | 64.5 | 88.7 | 96.1 |
| 2016 | 130 | 15.6 | 15.9 | 19.7 | 21.3 | 57.1 |
| 2016 | 30 | 15.8 | 17.5 | 17.7 | 23.3 | 68.7 |
| 2016 | 58 | 35.6 | 35.6 | 39.2 | 48.5 | 77.7 |
| 2017 | 200 | 38.4 | 40 | 44.7 | 51.3 | 73.3 |
| 2018 | 205 | 46.3 | 46.5 | 47.5 | 50.8 | 86.6 |
| 2018 | 30 | 50.5 | 54.2 | 59.5 | 67.2 | 76.4 |
| 2016 | 204 | 77.4 | 86.5 | 94.3 | 90.3 | 98 |
| ***Min*** | | 0.2 | 0.4 | 0.4 | 0.4 | 1 |
| ***Max*** | | 77.4 | 86.5 | 94.3 | 90.3 | 98 |
| ***Median*** | | 6 | 26 | 29.5 | 43 | 74.8 |
| ***Mean ± SD*** | | 18.6 ± 23.7 | 29.8 ± 25.2 | 37.1 ± 28.4 | 42.3 ± 30.9 | 65 ± 31.4 |

1, Plant species without confirmed hummingbird visitation were excluded.

2, The most frequently visited plant species were assigned a weight of 1; all other resources were downweighted accordingly (see Fig. S1).

### **Table S3.** Summary of visitation to *Heliconia tortuosa* within forest fragments (*N* = 14) surrounding the Las Cruces Biological Station in southern Costa Rica. Based on 1809 hours of video observation, green hermits (*Phaethornis guy*) were the most frequent *H. tortuosa* visitor (75% of 1055 total visits), followed by rufous-tailed hummingbirds (*Amazilia tzacatl,* 10%) and violet sabrewings (*Campylopterus hemileucurus*, 9%). Less frequent visitors comprised only ~5% of total visits. This summary only includes videos from control sites or treatment sites prior to *Heliconia* removal. Visit types were scored based on whether hummingbirds contacted a flower’s reproductive structures (‘honest’) or obtained nectar by bypassing the reproductive structures entirely (‘rob’).

|  |  |  | Visit type | | | |  |
| --- | --- | --- | --- | --- | --- | --- | --- |
| Common name | Scientific name | Sex | Honest | Rob | Honest and rob during same visit | Unknown | Total visits |
| Green hermit | *Phaethornis guy* | All | 725 | 11 | 12 | 44 | **792** |
|  |  | Female | 636 | 6 | 11 | 23 | 676 |
|  |  | Male | 73 | 5 | 1 | 19 | 98 |
|  |  | Unknown | 16 | 0 | 0 | 2 | 18 |
| Rufous-tailed hummingbird | *Amazilia tzacatl* | All | 72 | 7 | 7 | 23 | **109** |
|  |  | Unknown | 72 | 7 | 7 | 23 | 109 |
| Violet sabrewing | *Campylopterus hemileucurus* | All | 97 | 0 | 0 | 0 | **97** |
|  |  | Female | 84 | 0 | 0 | 0 | 84 |
|  |  | Male | 13 | 0 | 0 | 0 | 13 |
| Stripe-throated hermit | *Phaethornis striigularis* | All | 3 | 30 | 5 | 3 | **41** |
|  |  | Unknown | 3 | 30 | 5 | 3 | 41 |
| Scaly-breasted hummingbird | *Phaeochroa cuvierii* | All | 0 | 10 | 0 | 0 | **10** |
|  |  | Unknown | 0 | 10 | 0 | 0 | 10 |
| Purple-crowned fairy | *Heliothryx barroti* | All | 0 | 3 | 0 | 0 | **3** |
|  |  | Female | 0 | 3 | 0 | 0 | 3 |
| Long-billed hermit | *Phaethornis longirostris* | All | 1 | 0 | 0 | 0 | **1** |
|  |  | Unknown | 1 | 0 | 0 | 0 | 1 |
| Unknown species |  | All | 0 | 1 | 0 | 1 | **2** |
|  |  | Unknown | 0 | 1 | 0 | 1 | 2 |

| Analysis | Subanalysis | Bird group | Contrast (treatment) | Contrast (period) | Estimate | 2.5% | 97.5% |
| --- | --- | --- | --- | --- | --- | --- | --- |
| Mist net captures | Captures | All species | treatment / control | post / pre | 0.81 | 0.53 | 1.2 |
|  |  | *Heliconia* specialists | treatment / control | post / pre | 0.92 | 0.47 | 1.8 |
|  | Recaptures | All species | treatment / control | NA | 0.77 | 0.38 | 1.5 |
|  |  | *Heliconia* specialists | treatment / control | NA | 0.88 | 0.36 | 2.1 |
| Radio telemetry | All replicates | All species | treatment / control | post / pre | 0.82 | 0.47 | 1.4 |
|  |  | *Heliconia* specialists | treatment / control | post / pre | 0.95 | 0.48 | 1.9 |
|  | Without outlier replicate | All species | treatment / control | post / pre | 1.2 | 0.71 | 1.9 |
|  |  | *Heliconia* specialists | treatment / control | post / pre | 1.3 | 0.75 | 2.3 |
| Flower visitation (*Heliconia*) | All birds | All species | treatment / control | post / pre | 1.2 | 0.81 | 1.8 |
|  |  | *Heliconia* specialists | treatment / control | post / pre | 1.1 | 0.81 | 1.5 |
|  | Marked birds | All species | treatment / control | post / pre | 0.99 | 0.58 | 1.7 |
|  |  | *Heliconia* specialists | treatment / control | post / pre | 1 | 0.56 | 1.8 |
| Flower visitation (non-*Heliconia*) | All birds | All species | treatment / control | post / pre | 0.88 | 0.71 | 1.1 |
|  |  | *Heliconia* specialists | treatment / control | post / pre | 0.86 | 0.61 | 1.2 |
|  | Marked birds | All species | treatment / control | post / pre | 0.76 | 0.39 | 1.5 |
|  |  | *Heliconia* specialists | treatment / control | post / pre | 0.67 | 0.3 | 1.5 |
| Flower visitation  (non-*Heliconia,*  individual plant species)  All birds | *Calathea crotalifera* | All species | treatment / control | post / pre | 0.46 | 0.18 | 1.2 |
|  | *Scutellaria costaricana* | All species | treatment / control | post / pre | 0.73 | 0.45 | 1.2 |
|  | *Centropogon granulosus* | All species | treatment / control | post / pre | 0.84 | 0.38 | 1.9 |
|  | *Columnea polyantha* | All species | treatment / control | post / pre | 0.47 | 0.12 | 1.8 |
|  | *Hamelia patens* | All species | treatment / control | post / pre | 1.3 | 0.66 | 2.4 |
|  | *Pachystachys lutea* | All species | treatment / control | post / pre | 0.61 | 0.16 | 2.4 |
|  | *Stachytarpheta frantzii* | All species | treatment / control | post / pre | 0.45 | 0.17 | 1.2 |
|  | *Columnea raymondii* | All species | treatment / control | post / pre | 0.66 | 0.36 | 1.2 |
|  | *Costus barbatus* | All species | treatment / control | post / pre | 1.4 | 0.9 | 2.2 |
| Pollination success | *Hamelia patens* | NA | treatment / control | post / pre | 1.8 | 0.53 | 6.1 |
|  | *Heliconia* | NA | treatment / control | post / pre | 1 | 0.55 | 2 |
| Body condition (mass) |  | All species | treatment - control | post - pre | 0.0063 | -0.02 | 0.032 |
|  |  | *Heliconia* specialists | treatment - control | post - pre | 0.0041 | -0.034 | 0.042 |

### **Table S4.** Contrasts for the effect of experimental *Heliconia* removal (treatment *x* period interaction).

### **Table S5.** Summary of Poisson GLMMs (log link) examining the effect of experimental *Heliconia* removal on mist net captures of hummingbirds. Results are presented for a categorical treatment variable, Control/Treatment, and quantitative treatment variable, Ln(calories removed per ha + 1). Standardized coefficient estimates, Wald confidence intervals, Wald *P*-values, and random effect standard deviations are shown. Coefficients of interest are in bold.

| **Number of hummingbirds captured** | | All species | | | | |  | Green hermits & violet sabrewings | | | | |
| --- | --- | --- | --- | --- | --- | --- | --- | --- | --- | --- | --- | --- |
|  |  | Est | 2.5% | 97.5% | *z* | *P* |  | Est | 2.5% | 97.5% | *z* | *P* |
| Categorical treatment | *(Intercept)* | 1.64 | 1.18 | 2.1 | 7.06 | <0.001 |  | 1.09 | 0.59 | 1.59 | 4.27 | <0.001 |
|  | *Treatment* | 0.28 | -0.27 | 0.84 | 1 | 0.319 |  | 0.15 | -0.44 | 0.74 | 0.5 | 0.617 |
|  | *Post* | -0.05 | -0.37 | 0.27 | -0.33 | 0.744 |  | -0.46 | -0.96 | 0.04 | -1.82 | 0.069 |
|  | *Net-hours* | 0.14 | -0.1 | 0.39 | 1.13 | 0.257 |  | 0.15 | -0.1 | 0.4 | 1.18 | 0.238 |
|  | ***Treatment x Post*** | **-0.21** | **-0.63** | **0.2** | **-0.99** | **0.32** |  | **-0.09** | **-0.73** | **0.56** | **-0.26** | **0.792** |
|  | *(Random Intercept) Site* | 0.49 |  |  |  |  |  | 0.57 |  |  |  |  |
|  | *(Random Intercept) Replicate* | 0.38 |  |  |  |  |  | 0.00 |  |  |  |  |
| Quantitative treatment | *(Intercept)* | 1.79 | 1.45 | 2.13 | 10.34 | <0.001 |  | 1.17 | 0.79 | 1.55 | 5.99 | <0.001 |
|  | *Calories removed* | 0.08 | -0.2 | 0.36 | 0.57 | 0.569 |  | 0.04 | -0.24 | 0.32 | 0.26 | 0.793 |
|  | *Post* | -0.18 | -0.38 | 0.03 | -1.69 | 0.091 |  | -0.51 | -0.84 | -0.19 | -3.11 | 0.002 |
|  | *Net-hours* | 0.17 | -0.06 | 0.39 | 1.43 | 0.154 |  | 0.17 | -0.05 | 0.39 | 1.51 | 0.132 |
|  | ***Calories removed x Post*** | **-0.07** | **-0.29** | **0.14** | **-0.67** | **0.506** |  | **0** | **-0.33** | **0.32** | **-0.03** | **0.98** |
|  | *(Random Intercept) Site* | 0.50 |  |  |  |  |  | 0.57 |  |  |  |  |
|  | *(Random Intercept) Replicate* | 0.38 |  |  |  |  |  | 0.00 |  |  |  |  |

### **Table S6.** Summary of betabinomial GLMMs (logit link) examining the effect of experimental *Heliconia* removal on the proportion of time that individual radio-tagged hummingbirds spent in focal areas. Results are presented for a categorical treatment variable, Control/Treatment, and quantitative treatment variable, Ln(calories removed per ha + 1). Standardized coefficient estimates, Wald confidence intervals, Wald *P*-values, and random effect standard deviations are shown. Coefficients of interest are in bold. All replicates, including an outlier, are included in these results.

| **Proportion of time spent in focal area** | | All species | | | | |  | Green hermits & violet sabrewings | | | | |
| --- | --- | --- | --- | --- | --- | --- | --- | --- | --- | --- | --- | --- |
|  |  | Est | 2.5% | 97.5% | *z* | *P* |  | Est | 2.5% | 97.5% | *z* | *P* |
| Categorical treatment | *(Intercept)* | -0.75 | -1.51 | 0.02 | -1.91 | 0.056 |  | -0.8 | -1.56 | -0.05 | -2.08 | 0.037 |
|  | *Treatment* | -1.08 | -1.99 | -0.17 | -2.33 | 0.02 |  | -1.04 | -1.92 | -0.17 | -2.34 | 0.019 |
|  | *Post* | -0.17 | -0.7 | 0.36 | -0.63 | 0.529 |  | -0.31 | -0.94 | 0.32 | -0.98 | 0.329 |
|  | ***Treatment x Post*** | **-0.18** | **-0.89** | **0.52** | **-0.51** | **0.61** |  | **0** | **-0.85** | **0.84** | **-0.01** | **0.994** |
|  | *(Random Intercept) Site* | 0.44 |  |  |  |  |  |  | 0.48 |  |  |  |
|  | *(Random Intercept) Replicate* | 0.02 |  |  |  |  |  |  | 0.03 |  |  |  |
|  | *(Random Intercept) Bird* | 0.98 |  |  |  |  |  |  | 0.71 |  |  |  |
| Quantitative treatment | *(Intercept)* | -1.47 | -1.98 | -0.95 | -5.58 | <0.001 |  | -1.51 | -2.04 | -0.98 | -5.59 | <0.001 |
|  | *Calories removed* | -0.46 | -0.92 | 0 | -1.95 | 0.051 |  | -0.46 | -0.9 | -0.01 | -2.01 | 0.045 |
|  | *Post* | -0.29 | -0.65 | 0.06 | -1.61 | 0.108 |  | -0.31 | -0.74 | 0.11 | -1.45 | 0.147 |
|  | ***Calories removed x Post*** | **-0.1** | **-0.44** | **0.25** | **-0.55** | **0.582** |  | **-0.01** | **-0.42** | **0.4** | **-0.04** | **0.968** |
|  | *(Random Intercept) Site* | 0.46 |  |  |  |  |  | 0.50 |  |  |  |  |
|  | *(Random Intercept) Replicate* | 0.03 |  |  |  |  |  | 0.05 |  |  |  |  |
|  | *(Random Intercept) Bird* | 1.00 |  |  |  |  |  | 0.73 |  |  |  |  |

### **Table S7.** Summary of negative binomial GLMMS (log link) examining the effect of experimental *Heliconia* removal on hummingbird visitation to focal *Heliconia* plants, as ascertained by remote camera observations. Results are presented for a categorical treatment variable, Control/Treatment, and quantitative treatment variable, Ln(calories removed per ha + 1). Standardized coefficient estimates, Wald confidence intervals, Wald *P*-values, and random effect standard deviations are shown. Coefficients of interest are in bold. ‘Flowers’ is the mean number of flowers present per day.

| **Visits to focal *Heliconia* plants** | | All species | | | | |  | Green hermits & violet sabrewings | | | | |
| --- | --- | --- | --- | --- | --- | --- | --- | --- | --- | --- | --- | --- |
|  |  | Est | 2.5% | 97.5% | *z* | *P* |  | Est | 2.5% | 97.5% | *z* | *P* |
| Categorical treatment | *(Intercept)* | -0.61 | -0.95 | -0.26 | -3.41 | 0.001 |  | -1.05 | -1.59 | -0.51 | -3.8 | <0.001 |
|  | *Treatment* | 0.06 | -0.29 | 0.4 | 0.32 | 0.75 |  | 0.16 | -0.31 | 0.63 | 0.66 | 0.508 |
|  | *Post* | -0.16 | -0.46 | 0.14 | -1.06 | 0.287 |  | -0.11 | -0.34 | 0.12 | -0.96 | 0.337 |
|  | *Flowers* | 0.13 | -0.03 | 0.28 | 1.57 | 0.116 |  | 0.14 | 0 | 0.27 | 2.01 | 0.044 |
|  | ***Treatment x Post*** | **0.2** | **-0.2** | **0.6** | **0.97** | **0.332** |  | **0.11** | **-0.2** | **0.42** | **0.7** | **0.483** |
|  | *(Random Intercept) Site* | 0.47 |  |  |  |  |  | 0.79 |  |  |  |  |
|  | *(Random Intercept) Replicate* | 0.22 |  |  |  |  |  | 0.51 |  |  |  |  |
| Quantitative treatment | *(Intercept)* | -0.57 | -0.87 | -0.27 | -3.74 | <0.001 |  | -0.96 | -1.44 | -0.48 | -3.94 | <0.001 |
|  | *Calories removed* | 0.06 | -0.12 | 0.23 | 0.65 | 0.518 |  | 0.13 | -0.11 | 0.37 | 1.08 | 0.28 |
|  | *Post* | -0.06 | -0.26 | 0.14 | -0.6 | 0.55 |  | -0.06 | -0.21 | 0.1 | -0.7 | 0.481 |
|  | *Flowers* | 0.13 | -0.03 | 0.28 | 1.54 | 0.123 |  | 0.14 | 0 | 0.28 | 2 | 0.045 |
|  | ***Calories removed x Post*** | **0.09** | **-0.12** | **0.29** | **0.85** | **0.397** |  | **0.05** | **-0.1** | **0.2** | **0.63** | **0.531** |
|  | *(Random Intercept) Site* | 0.46 |  |  |  |  |  | 0.79 |  |  |  |  |
|  | *(Random Intercept) Replicate* | 0.21 |  |  |  |  |  | 0.50 |  |  |  |  |

### **Table S8.** Summary of negative binomial GLMMS (log link) examining the effect of experimental *Heliconia* removal on hummingbird visitation to non-*Heliconia* plant species, as ascertained by remote camera observations. Results are presented for a categorical treatment variable, Control/Treatment, and quantitative treatment variable, Ln(calories removed per ha + 1). Standardized coefficient estimates, Wald confidence intervals, Wald *P*-values, and random effect standard deviations are shown. Coefficients of interest are in bold. ‘Flowers’ is the mean number of flowers present per day.

| **Visits to non-*Heliconia* plant species** | | All species | | | | |  | Green hermits & violet sabrewings | | | | |
| --- | --- | --- | --- | --- | --- | --- | --- | --- | --- | --- | --- | --- |
|  |  | Est | 2.5% | 97.5% | *z* | *P* |  | Est | 2.5% | 97.5% | *z* | *P* |
| Categorical treatment | *(Intercept)* | -2.06 | -2.42 | -1.71 | -11.46 | <0.001 |  | -4.12 | -4.81 | -3.43 | -11.68 | <0.001 |
|  | *Treatment* | 0.37 | 0.02 | 0.72 | 2.1 | 0.036 |  | 0.94 | 0.17 | 1.72 | 2.38 | 0.017 |
|  | *Post* | 0.17 | 0 | 0.34 | 2 | 0.046 |  | 0.12 | -0.15 | 0.38 | 0.87 | 0.382 |
|  | *Flowers* | 0.15 | 0.03 | 0.26 | 2.48 | 0.013 |  | 0.21 | 0 | 0.42 | 1.98 | 0.048 |
|  | ***Treatment x Post*** | **-0.13** | **-0.35** | **0.09** | **-1.14** | **0.255** |  | **-0.15** | **-0.49** | **0.18** | **-0.9** | **0.369** |
|  | *(Random Intercept) Site* | 0.47 |  |  |  |  |  | 0.58 |  |  |  |  |
|  | *(Random Intercept) Replicate* | 0.14 |  |  |  |  |  | 0.66 |  |  |  |  |
|  | *(Random Intercept) Plant species* | 0.88 |  |  |  |  |  | 1.6 |  |  |  |  |
| Quantitative treatment | *(Intercept)* | -1.87 | -2.17 | -1.57 | -12.12 | <0.001 |  | -3.62 | -4.16 | -3.07 | -13.07 | <0.001 |
|  | *Calories removed* | 0.2 | 0.03 | 0.38 | 2.25 | 0.025 |  | 0.53 | 0.15 | 0.91 | 2.71 | 0.007 |
|  | *Post* | 0.1 | -0.01 | 0.21 | 1.87 | 0.061 |  | 0.04 | -0.13 | 0.2 | 0.45 | 0.656 |
|  | *Flowers* | 0.15 | 0.03 | 0.26 | 2.49 | 0.013 |  | 0.21 | 0 | 0.42 | 1.98 | 0.048 |
|  | ***Calories removed x Post*** | **-0.07** | **-0.18** | **0.04** | **-1.19** | **0.235** |  | **-0.06** | **-0.23** | **0.1** | **-0.77** | **0.441** |
|  | *(Random Intercept) Site* | 0.47 |  |  |  |  |  | 0.65 |  |  |  |  |
|  | *(Random Intercept) Replicate* | 0.12 |  |  |  |  |  | 0.58 |  |  |  |  |
|  | *(Random Intercept) Plant species* | 0.88 |  |  |  |  |  | 1.59 |  |  |  |  |

### **Table S9.** Summary of Poisson and negative binomial GLMMS (log link) examining the effect of experimental *Heliconia* removal on hummingbird visitation to individual non-*Heliconia* plant species, as ascertained by remote camera observations. Standardized coefficient estimates, Wald confidence intervals, Wald *P*-values, and random effect standard deviations are shown. Coefficients of interest are in bold. ‘Flowers’ is the mean number of flowers present per day.

| **Visits to individual non-*Heliconia* plant species** | | All species | | | | |
| --- | --- | --- | --- | --- | --- | --- |
|  |  | Est | 2.5% | 97.5% | *z* | *P* |
| *Calathea marantifolia* | *(Intercept)* | -3.98 | -5.23 | -2.72 | -6.2 | <0.001 |
|  | *Treatment* | 1.83 | 0.37 | 3.28 | 2.46 | 0.014 |
|  | *Post* | 0.46 | -0.24 | 1.16 | 1.3 | 0.194 |
|  | *Flowers* | 0.22 | -0.25 | 0.7 | 0.91 | 0.364 |
|  | ***Treatment x Post*** | **-0.77** | **-1.67** | **0.13** | **-1.68** | **0.093** |
|  | *(Random Intercept) Site* | 0.00 |  |  |  |  |
|  | *(Random Intercept) Replicate* | 1.37 |  |  |  |  |
| *Scutellaria costaricana* | *(Intercept)* | -1.89 | -2.33 | -1.45 | -8.41 | <0.001 |
|  | *Treatment* | 0.54 | 0.04 | 1.03 | 2.12 | 0.034 |
|  | *Post* | 0.04 | -0.31 | 0.4 | 0.24 | 0.81 |
|  | *Flowers* | 0.13 | -0.08 | 0.33 | 1.21 | 0.227 |
|  | ***Treatment x Post*** | **-0.32** | **-0.78** | **0.15** | **-1.34** | **0.18** |
|  | *(Random Intercept) Site* | 0.49 |  |  |  |  |
|  | *(Random Intercept) Replicate* | 0.31 |  |  |  |  |
| *Centropogon granulosus* | *(Intercept)* | -2.76 | -3.5 | -2.02 | -7.32 | <0.001 |
|  | *Treatment* | -0.23 | -0.91 | 0.45 | -0.67 | 0.506 |
|  | *Post* | 0.04 | -0.49 | 0.57 | 0.16 | 0.872 |
|  | *Flowers* | -0.11 | -0.33 | 0.12 | -0.93 | 0.35 |
|  | ***Treatment x Post*** | **-0.17** | **-0.94** | **0.6** | **-0.44** | **0.657** |
|  | *(Random Intercept) Site* | 0.98 |  |  |  |  |
|  | *(Random Intercept) Replicate* | 0.01 |  |  |  |  |
| *Columnea polyantha* | *(Intercept)* | -3.62 | -4.67 | -2.56 | -6.72 | <0.001 |
|  | *Treatment* | 1.93 | 0.72 | 3.13 | 3.12 | 0.002 |
|  | *Post* | 0.75 | -0.31 | 1.8 | 1.38 | 0.166 |
|  | *Flowers* | 0.44 | 0.04 | 0.84 | 2.18 | 0.03 |
|  | ***Treatment x Post*** | **-0.76** | **-2.04** | **0.53** | **-1.16** | **0.247** |
|  | *(Random Intercept) Site* | 0.34 |  |  |  |  |
|  | *(Random Intercept) Replicate* | 0.37 |  |  |  |  |
| *Hamelia patens* | *(Intercept)* | -2.14 | -2.65 | -1.63 | -8.18 | <0.001 |
|  | *Treatment* | 0.18 | -0.4 | 0.76 | 0.6 | 0.548 |
|  | *Post* | 0.27 | -0.18 | 0.73 | 1.17 | 0.243 |
|  | *Flowers* | -0.15 | -0.36 | 0.06 | -1.37 | 0.171 |
|  | ***Treatment x Post*** | **0.23** | **-0.38** | **0.85** | **0.74** | **0.459** |
|  | *(Random Intercept) Site* | 0.52 |  |  |  |  |
|  | *(Random Intercept) Replicate* | 0.00 |  |  |  |  |
| *Pachystachys lutea* | *(Intercept)* | -2.36 | -3.28 | -1.44 | -5.03 | <0.001 |
|  | *Treatment* | 0.87 | -0.26 | 2 | 1.51 | 0.13 |
|  | *Post* | 0.81 | -0.19 | 1.81 | 1.59 | 0.111 |
|  | *Flowers* | -0.24 | -0.74 | 0.26 | -0.95 | 0.34 |
|  | ***Treatment x Post*** | **-0.49** | **-1.78** | **0.8** | **-0.74** | **0.459** |
|  | *(Random Intercept) Site* | 0.00 |  |  |  |  |
|  | *(Random Intercept) Replicate* | 0.54 |  |  |  |  |
| *Stachytarpheta frantzii* | *(Intercept)* | 2.26 | 1.64 | 2.87 | 7.2 | <0.001 |
|  | *Treatment* | 1.04 | 0.32 | 1.76 | 2.82 | 0.005 |
|  | *Post* | 0.33 | -0.38 | 1.04 | 0.91 | 0.365 |
|  | *Flowers^2^* | -0.13 | -0.36 | 0.09 | -1.15 | 0.25 |
|  | *Hours* | 0.69 | 0.38 | 1 | 4.36 | <0.001 |
|  | ***Treatment x Post*** | **-0.79** | **-1.72** | **0.14** | **-1.66** | **0.096** |
|  | *(Random Intercept) Site* | 0.00 |  |  |  |  |
|  | *(Random Intercept) Replicate* | 0.46 |  |  |  |  |
| *Columnea raymondii* | *(Intercept)* | -2.42 | -3.35 | -1.48 | -5.07 | <0.001 |
|  | *Treatment* | 0.1 | -1.1 | 1.3 | 0.17 | 0.868 |
|  | *Post* | 0.12 | -0.23 | 0.47 | 0.69 | 0.492 |
|  | *Flowers* | 0.42 | 0.11 | 0.73 | 2.63 | 0.008 |
|  | ***Treatment x Post*** | **-0.41** | **-0.99** | **0.17** | **-1.37** | **0.17** |
|  | *(Random Intercept) Site* | 0.87 |  |  |  |  |
|  | *(Random Intercept) Replicate* | 0.80 |  |  |  |  |
| *Costus barbatus* | *(Intercept)* | -0.52 | -0.97 | -0.07 | -2.29 | 0.022 |
|  | *Treatment* | -0.32 | -0.89 | 0.24 | -1.12 | 0.264 |
|  | *Post* | -0.3 | -0.65 | 0.04 | -1.71 | 0.087 |
|  | *Flowers* | -0.12 | -0.29 | 0.05 | -1.42 | 0.155 |
|  | ***Treatment x Post*** | **0.34** | **-0.07** | **0.74** | **1.61** | **0.107** |
|  | *(Random Intercept) Site* | 0.28 |  |  |  |  |
|  | *(Random Intercept) Replicate* | 0.29 |  |  |  |  |

### **Table S10.** Summary of binomial GLMMs (logit link) examining the effect of experimental *Heliconia* removal on pollination success in *Heliconia tortuosa* and *Hamelia patens* at stations (focal arrays). Results are presented for a categorical treatment variable, Control/Treatment, and quantitative treatment variable, Ln(calories removed per ha + 1). Standardized coefficient estimates, Wald confidence intervals, Wald *P*-values, and random effect standard deviations are shown. Coefficients of interest are in bold.

| **Proportion of styles with pollen tubes** | | *Heliconia tortuosa* | | | | |  | *Hamelia patens* | | | | |
| --- | --- | --- | --- | --- | --- | --- | --- | --- | --- | --- | --- | --- |
|  |  | Est | 2.5% | 97.5% | *z* | *P* |  | Est | 2.5% | 97.5% | *z* | *P* |
| Categorical treatment | *(Intercept)* | -0.19 | -0.99 | 0.61 | -0.47 | 0.637 |  | -0.03 | -1.31 | 1.24 | -0.05 | 0.959 |
|  | *Treatment* | 0.03 | -1.1 | 1.17 | 0.05 | 0.957 |  | -1.89 | -3.43 | -0.35 | -2.41 | 0.016 |
|  | *Post* | -0.01 | -0.76 | 0.74 | -0.03 | 0.978 |  | -0.32 | -1.34 | 0.69 | -0.62 | 0.532 |
|  | ***Treatment x Post*** | **0.09** | **-1.07** | **1.25** | **0.15** | **0.877** |  | **0.81** | **-0.73** | **2.35** | **1.03** | **0.302** |
|  | *(Random Intercept) Site* | 0.42 |  |  |  |  |  | 1.34 |  |  |  |  |
|  | *(Random Intercept) Replicate* | 0.62 |  |  |  |  |  | 0.00 |  |  |  |  |
| Quantitative treatment | *(Intercept)* | -0.17 | -0.74 | 0.4 | -0.58 | 0.563 |  | -0.85 | -1.88 | 0.18 | -1.61 | 0.107 |
|  | *Calories removed* | 0.19 | -0.41 | 0.79 | 0.63 | 0.528 |  | -0.94 | -1.75 | -0.14 | -2.3 | 0.022 |
|  | *Post* | 0.02 | -0.63 | 0.68 | 0.06 | 0.951 |  | 0.07 | -0.69 | 0.82 | 0.17 | 0.863 |
|  | ***Calories removed x Post*** | **-0.02** | **-0.72** | **0.68** | **-0.07** | **0.948** |  | **0.37** | **-0.42** | **1.16** | **0.92** | **0.357** |
|  | *(Random Intercept) Site* | 0.38 |  |  |  |  |  | 1.28 |  |  |  |  |
|  | *(Random Intercept) Replicate* | 0.58 |  |  |  |  |  | 0.00 |  |  |  |  |

### **Table S11.** Summary of binomial GLMMs (logit link) examining the effect of experimental *Heliconia* removal on recapture probability, i.e., the proportion of individual hummingbirds captured during the ‘pre’ period that were recaptured during the ‘post’ period. Results are presented for a categorical treatment variable, Control/Treatment, and quantitative treatment variable, Ln(calories removed per ha + 1). Standardized coefficient estimates, Wald confidence intervals, Wald *P*-values, and random effect standard deviations are shown. Coefficients of interest are in bold.

| **Proportion of hummingbirds recaptured** | | All species | | | | |  | Green hermits & violet sabrewings | | | | |
| --- | --- | --- | --- | --- | --- | --- | --- | --- | --- | --- | --- | --- |
|  |  | Est | 2.5% | 97.5% | *z* | *P* |  | Est | 2.5% | 97.5% | *z* | *P* |
| Categorical treatment | *(Intercept)* | -1.49 | -2.07 | -0.91 | -5.03 | <0.001 |  | -1.32 | -2.1 | -0.54 | -3.32 | 0.001 |
|  | ***Treatment*** | **-0.31** | **-1.11** | **0.48** | **-0.77** | **0.439** |  | **-0.16** | **-1.2** | **0.88** | **-0.3** | **0.763** |
|  | *(Random Intercept) Site* | 0.00 |  |  |  |  |  | 0.00 |  |  |  |  |
| Quantitative treatment | *(Intercept)* | -1.65 | -2.05 | -1.25 | -8.13 | <0.001 |  | -1.41 | -1.93 | -0.89 | -5.28 | <0.001 |
|  | ***Calories removed*** | **-0.14** | **-0.55** | **0.27** | **-0.66** | **0.511** |  | **-0.04** | **-0.58** | **0.5** | **-0.14** | **0.889** |
|  | *(Random Intercept) Site* | 0.00 |  |  |  |  |  | 0.06 |  |  |  |  |

### **Table S12.** Summary of Poisson GLMMs (log link) examining the effect of experimental *Heliconia* removal on marked hummingbird visitation to focal *Heliconia* plants, as ascertained by remote camera observations. Results are presented for a categorical treatment variable, Control/Treatment, and quantitative treatment variable, Ln(calories removed per ha + 1). Standardized coefficient estimates, Wald confidence intervals, Wald *P*-values, and random effect standard deviations are shown. Coefficients of interest are in bold. ‘Flowers’ is the mean number of flowers present per day.

| **Visits to focal *Heliconia* plants from marked birds (2017-2018 only)** | | All species | | | | |  | Green hermits & violet sabrewings | | | | |
| --- | --- | --- | --- | --- | --- | --- | --- | --- | --- | --- | --- | --- |
|  |  | Est | 2.5% | 97.5% | *z* | *P* |  | Est | 2.5% | 97.5% | *z* | *P* |
| Categorical treatment | *(Intercept)* | -2.36 | -2.96 | -1.76 | -7.68 | <0.001 |  | -2.19 | -2.8 | -1.58 | -7.05 | <0.001 |
|  | *Treatment* | -0.17 | -1.04 | 0.71 | -0.38 | 0.707 |  | -0.19 | -1.11 | 0.72 | -0.41 | 0.682 |
|  | *Post* | 0.13 | -0.28 | 0.55 | 0.64 | 0.524 |  | 0.18 | -0.25 | 0.62 | 0.83 | 0.409 |
|  | *Flowers* | 0.03 | -0.19 | 0.25 | 0.26 | 0.797 |  | 0 | -0.24 | 0.24 | 0 | 0.997 |
|  | ***Treatment x Post*** | **-0.01** | **-0.53** | **0.51** | **-0.05** | **0.962** |  | **0** | **-0.55** | **0.56** | **0.02** | **0.986** |
|  | *(Random Intercept) Site* | 0.00 |  |  |  |  |  | 0.00 |  |  |  |  |
|  | *(Random Intercept) Replicate* | 0.00 |  |  |  |  |  | 0.00 |  |  |  |  |
|  | *(Random Intercept) Bird* | 0.98 |  |  |  |  |  | 0.92 |  |  |  |  |
| Quantitative treatment | *(Intercept)* | -2.44 | -2.88 | -1.99 | -10.76 | <0.001 |  | -2.28 | -2.74 | -1.82 | -9.69 | <0.001 |
|  | *Calories removed* | -0.11 | -0.55 | 0.33 | -0.48 | 0.63 |  | -0.14 | -0.6 | 0.33 | -0.59 | 0.558 |
|  | *Post* | 0.13 | -0.14 | 0.41 | 0.94 | 0.346 |  | 0.19 | -0.1 | 0.48 | 1.27 | 0.204 |
|  | *Flowers* | 0.03 | -0.2 | 0.25 | 0.22 | 0.824 |  | -0.01 | -0.25 | 0.24 | -0.04 | 0.966 |
|  | ***Calories removed x Post*** | **-0.02** | **-0.29** | **0.26** | **-0.13** | **0.899** |  | **-0.01** | **-0.31** | **0.29** | **-0.06** | **0.951** |
|  | *(Random Intercept) Site* | 0.00 |  |  |  |  |  | 0.00 |  |  |  |  |
|  | *(Random Intercept) Replicate* | 0.00 |  |  |  |  |  | 0.00 |  |  |  |  |
|  | *(Random Intercept) Bird* | 0.98 |  |  |  |  |  | 0.92 |  |  |  |  |

### **Table S13.** Summary of negative binomial GLMMS (log link) examining the effect of experimental *Heliconia* removal on marked hummingbird visitation to non-*Heliconia* plant species, as ascertained by remote camera observations. Results are presented for a categorical treatment variable, Control/Treatment, and quantitative treatment variable, Ln(calories removed per ha + 1). Standardized coefficient estimates, Wald confidence intervals, Wald *P*-values, and random effect standard deviations are shown. Coefficients of interest are in bold. ‘Flowers’ is the mean number of flowers present per day.

| **Visits to non-*Heliconia* plant species from marked birds (2017-2018 only)** | | All species | | | | |  | Green hermits & violet sabrewings | | | | |
| --- | --- | --- | --- | --- | --- | --- | --- | --- | --- | --- | --- | --- |
|  |  | Est | 2.5% | 97.5% | *z* | *P* |  | Est | 2.5% | 97.5% | *z* | *P* |
| Categorical treatment | *(Intercept)* | -3.25 | -3.76 | -2.73 | -12.32 | <0.001 |  | -3.2 | -3.8 | -2.61 | -10.55 | <0.001 |
|  | *Treatment* | 0.47 | -0.21 | 1.15 | 1.35 | 0.178 |  | 0.96 | 0.16 | 1.75 | 2.36 | 0.018 |
|  | *Post* | 0.18 | -0.34 | 0.7 | 0.68 | 0.5 |  | 0.02 | -0.57 | 0.6 | 0.06 | 0.955 |
|  | *Flowers* | 0.03 | -0.18 | 0.25 | 0.3 | 0.76 |  | 0 | -0.29 | 0.3 | 0.02 | 0.981 |
|  | ***Treatment x Post*** | **-0.28** | **-0.94** | **0.39** | **-0.82** | **0.413** |  | **-0.39** | **-1.19** | **0.4** | **-0.97** | **0.333** |
|  | *(Random Intercept) Site* | 0.00 |  |  |  |  |  | 0.00 |  |  |  |  |
|  | *(Random Intercept) Replicate* | 0.00 |  |  |  |  |  | 0.00 |  |  |  |  |
|  | *(Random Intercept) Plant species* | 0.52 |  |  |  |  |  | 0.70 |  |  |  |  |
|  | *(Random Intercept) Bird* | 0.47 |  |  |  |  |  | 0.22 |  |  |  |  |
| Quantitative treatment | *(Intercept)* | -2.99 | -3.36 | -2.62 | -15.89 | <0.001 |  | -2.79 | -3.24 | -2.34 | -12.11 | <0.001 |
|  | *Calories removed* | 0.27 | -0.06 | 0.6 | 1.58 | 0.113 |  | 0.47 | 0.08 | 0.87 | 2.33 | 0.02 |
|  | *Post* | 0.04 | -0.29 | 0.36 | 0.21 | 0.833 |  | -0.15 | -0.55 | 0.24 | -0.75 | 0.451 |
|  | *Flowers* | 0.04 | -0.18 | 0.25 | 0.33 | 0.744 |  | 0.01 | -0.29 | 0.3 | 0.05 | 0.957 |
|  | ***Calories removed x Post*** | **-0.18** | **-0.5** | **0.15** | **-1.06** | **0.291** |  | **-0.21** | **-0.6** | **0.18** | **-1.05** | **0.293** |
|  | *(Random Intercept) Site* | 0.00 |  |  |  |  |  | 0.00 |  |  |  |  |
|  | *(Random Intercept) Replicate* | 0.00 |  |  |  |  |  | 0.00 |  |  |  |  |
|  | *(Random Intercept) Plant species* | 0.51 |  |  |  |  |  | 0.71 |  |  |  |  |
|  | *(Random Intercept) Bird* | 0.45 |  |  |  |  |  | 0.27 |  |  |  |  |

### **Table S14.** Summary of LMMs examining the effect of experimental *Heliconia* removal on relative body mass for individual hummingbirds captured during each capture session (pre and post). Relative body mass controls for differences in structural size among hummingbirds and was calculated by subtracting observed body mass from the body mass predicted by species-specific allometric equations [i.e., Ln(body mass) ~ Ln(wing length)]. Results are presented for a categorical treatment variable, Control/Treatment, and quantitative treatment variable, Ln(calories removed per ha + 1). Standardized coefficient estimates, Wald confidence intervals, Wald *P*-values, and random effect standard deviations are shown. Coefficients of interest are in bold. Random effects for Site and Replicate ID were not included due to small sample size and problems with model convergence. Coefficients of interest are in bold.

| **Hummingbird body mass** | | All species  (30 individuals) | | | | |  | Green hermits & violet sabrewings  (16 individuals) | | | | |
| --- | --- | --- | --- | --- | --- | --- | --- | --- | --- | --- | --- | --- |
|  |  | Est | 2.5% | 97.5% | *z* | *P* |  | Est | 2.5% | 97.5% | *z* | *P* |
| Categorical treatment | *(Intercept)* | 0.99 | 0.97 | 1.01 | 107.72 | <0.001 |  | 0.98 | 0.96 | 1.01 | 81.72 | <0.001 |
|  | *Treatment* | 0 | -0.03 | 0.02 | -0.25 | 0.805 |  | -0.02 | -0.05 | 0.02 | -0.94 | 0.349 |
|  | *Post* | 0.01 | -0.01 | 0.03 | 0.7 | 0.484 |  | 0.02 | 0 | 0.05 | 1.61 | 0.107 |
|  | ***Treatment x Post*** | **0.01** | **-0.02** | **0.03** | **0.49** | **0.626** |  | **0** | **-0.03** | **0.04** | **0.22** | **0.826** |
|  | *(Random Intercept) Bird* | 0.02 |  |  |  |  |  | 0.02 |  |  |  |  |
| Quantitative treatment | *(Intercept)* | 0.99 | 0.98 | 1 | 164.16 | <0.001 |  | 0.97 | 0.96 | 0.99 | 115.03 | <0.001 |
|  | *Calories removed* | -0.01 | -0.02 | 0.01 | -0.85 | 0.393 |  | -0.01 | -0.03 | 0.01 | -1.03 | 0.301 |
|  | *Post* | 0.01 | 0 | 0.02 | 1.63 | 0.103 |  | 0.02 | 0.01 | 0.04 | 2.51 | 0.012 |
|  | ***Calories removed x Post*** | **0** | **-0.01** | **0.02** | **0.74** | **0.457** |  | **0** | **-0.02** | **0.02** | **0.29** | **0.776** |
|  | *(Random Intercept) Bird* | 0.02 |  |  |  |  |  | 0.02 |  |  |  |  |

### **Table S15.** Natural variation in *H. tortuosa* density within 14 forest fragments in southern Costa Rica, measured 1-3 times across three years (2016-2018).

|  |  | *H. tortuosa* density | | |
| --- | --- | --- | --- | --- |
| Site | Replicate | Inflorescences per ha | Flowers per ha  (estimated) | Calories per ha  (estimated)^1^ |
| 10 | 2016_10 | 169 | 135 | 7462 |
| 10 | 2017_10 | 35 | 19 | 1045 |
| 10 | 2018_10 | 155 | 109 | 6013 |
| 24 | 2017_24 | 118 | 48 | 2650 |
| 24 | 2018_24 | 235 | 164 | 9022 |
| 29 | 2018_29 | 440 | 283 | 15568 |
| 30 | 2016_30 | 221 | 157 | 8664 |
| 30 | 2017_30 | 189 | 81 | 4482 |
| 30 | 2018_30 | 448 | 356 | 19618 |
| 49 | 2017_49 | 197 | 124 | 6849 |
| 58 | 2016_58 | 201 | 111 | 6124 |
| 58 | 2017_58 | 52 | 11 | 620 |
| 58 | 2018_58 | 103 | 64 | 3520 |
| 60 | 2017_60 | 67 | 30 | 1678 |
| 60 | 2018_60 | 335 | 186 | 10265 |
| 130 | 2016_130 | 67 | 35 | 1923 |
| 130 | 2017_130 | 41 | 15 | 842 |
| 137 | 2016_137 | 219 | 160 | 8802 |
| 137 | 2017_137 | 72 | 29 | 1616 |
| 137 | 2018_137 | 134 | 129 | 7112 |
| 200 | 2017_200 | 165 | 58 | 3189 |
| 201 | 2017_201 | 11 | 5 | 287 |
| 203 | 2017_203 | 3 | 2 | 134 |
| 204 | 2016_204 | 15 | 12 | 646 |
| 204 | 2017_204 | 16 | 7 | 404 |
| 205 | 2018_205 | 1131 | 372 | 20481 |
| ***Min*** | | 3 | 2 | 134 |
| ***Max*** | | 1131 | 372 | 20481 |
| ***Median*** | | 144 | 72 | 4001 |
| ***Mean ± SD*** | | 186 ± 228 | 104 ± 104 | 5731 ± 5729 |

1, Unit is calories, not kilocalories.
